# A trehalase-derived MAMP triggers LecRK-V-mediated immune responses in Arabidopsis

**DOI:** 10.1101/2025.02.02.636166

**Authors:** Erika Iino, Yasuhiro Kadota, Noriko Maki, Erika Ono, Kazuki Sato, Nobuaki Ishihama, Bruno Pok Man Ngou, Marc W Schmid, Takamasa Suzuki, Taketo Uehara, Ken Shirasu

**Affiliations:** RIKEN Center for Sustainable Resource Science (CSRS), Suehiro-cho 1-7-22 Tsurumi-ku, Yokohama, Kanagawa, 230-0045, Japan; Graduate school of Science, The University of Tokyo, 7-3-1, Hongo, Bunkyo-ku, Tokyo, 113-8654, Japan; MWSchmid GmbH, Glarus, Switzerland; College of Bioscience and Biotechnology, Chubu University, Kasugai, 487-0027, Japan; Institute for Plant Protection, National Agriculture and Food Research Organization, Tsukuba, Japan

## Abstract

Plant-parasitic nematodes (PPNs) cause major agricultural losses worldwide, yet the molecular basis of plant immunity against these pathogens remains poorly understood. To investigate how plants recognize PPNs, we aimed to identify microbe-associated molecular patterns (MAMPs) from nematodes and the corresponding plant immune components. Due to the limited availability of material from obligate PPNs, we used *Caenorhabditis elegans*, a free-living nematode, as a MAMP source. *C. elegans* extracts activated MAMP-triggered immune responses in *Arabidopsis* Col-0. Through chromatography-based purification, we identified a secreted trehalase and pinpointed a conserved peptide region essential for its MAMP activity. A corresponding peptide from root-knot nematode trehalase enabled the identification of lectin receptor kinases LecRK-V.5 and LecRK-V.6 as key components in immune induction. Notably, this peptide region is conserved across insect and fungal pathogens, with LecRK-Vs required for immune responses to these peptides, highlighting the role of LecRK-V-mediated mechanism for broad-spectrum pathogen detection via trehalase-derived peptides.

**Teaser:** Trehalase-derived peptides trigger LecRK-V-mediated immune responses in plants, enabling broad detection of nematodes, insects, and fungi.

## Introduction

Plant-parasitic nematodes (PPNs) rank among the most destructive agricultural pests, causing an estimated annual global crop loss of around 175 billion USD (*1, 2*). The most economically significant PPNs are sedentary endoparasites, including root-knot nematodes (RKNs) and cyst nematodes (CNs) (*1*). RKNs exhibit a broad host range, infecting thousands of plant species (*1*), while CNs, though more selective, target important crops such as potatoes and soybeans (*1*). Upon root penetration, these endoparasites secrete a suite of effectors, from their subventral gland cells into the plant apoplast to facilitate tissue migration (*3*). Notably, CNs migrate intracellularly, while RKNs move intercellularly. Once reaching the vascular tissue, these nematodes secrete effectors from the dorsal gland cell, reprogramming plant cells into specialized feeding structures, such as giant cells or syncytia, which sustain them throughout their sedentary stage (*4*).

Plants detect microbes through the perception of microbe-associated molecular patterns (MAMPs) via surface-localized pattern recognition receptors (PRRs), leading to the activation of MAMP-triggered immunity (MTI) (*5*). A well-characterized example of a MAMP is flg22, a peptide derived from flagellin originally identified in the plant pathogenic *Pseudomonas* (*6*), which is recognized by the PRR FLS2 (*7*). Another notable MAMP is elf18, a peptide from bacterial elongation factor Tu, isolated from non-pathogenic *E. coli* (*8*), and perceived by the PRR EFR (*9*). Similarly, N-acetylchitooligosaccharides (chitin), fungal MAMPs (*10*), are detected by the PRR CERK1 (*11*). Activation of such PRRs triggers a series of downstream immune responses, including calcium influx (*12, 13*) and reactive oxygen species (ROS) bursts (*14, 15*), cell wall fortification through lignin and callose deposition, activation of mitogen-activated protein kinases (MAPKs), phytoalexin biosynthesis, and the induction of defense-related genes (*16*). Since nematodes maintain close physical contact with plant cells during both migratory and sedentary stages, MTI likely plays a crucial role in plant defense against these parasites (*17*). Thus, understanding the molecular basis of nematode recognition, likely via their MAMPs, is essential for advancing our knowledge of plant immunity against PPNs.

Previous studies have identified the conserved nematode pheromone ascaroside (ascr#18) as the first nematode-derived MAMP (*18*). Ascr#18 was shown to be recognized by the PRR NEMATODE-INDUCED LEUCINE RICH REPEAT-RECEPTOR LIKE KINASE (LRR-RLK) 1 (NILR1), which was originally identified to be involved in MTI induction in response to nematode-derived compound (*19–21*). Although this is the only currently proposed MAMP-PRR pair in nematode-plant interactions, many more MAMPs and their corresponding PRRs likely remain to be discovered. Discovering new MAMP-PRR pairs in nematode-plant interactions will be key to understanding how plants defend themselves against these persistent agricultural pests.

In this study, we describe the identification of a novel MAMP peptide derived from secreted trehalase in nematodes using a biochemical approach. Utilizing the natural variation in *Arabidopsis thaliana,* we successfully mapped the locus responsible for immune responses triggered by the MAMP peptide. This locus contains a gene cluster encoding L-type Lectin Receptor Kinases (LecRKs), and knockout analysis confirmed that *LecRK-V.5* and *LecRK-V.6* as key components required for the MAMP-inducible immune responses. Intriguingly, the MAMP sequence was conserved not only in nematodes but also across a range of other plant pathogens, suggesting that secreted trehalase may be a common target for plant immune recognition. This conservation highlights the significance of trehalase, a trehalose-degrading enzyme, as a crucial factor for diverse pathogens employing distinct infection strategies. These findings expand our understanding of MAMP-triggered plant immunity and suggest broad implications for developing resistance strategies against multiple pathogens.

## Results

### MAMP potential in nematode extracts

To elucidate the molecular mechanisms underlying plant recognition of nematodes, we sought to identify MAMPs from PPNs and corresponding plant genes that are required for MAMP-triggered immune responses. However, the obligate parasitism of PPNs presents significant technical challenges in obtaining sufficient material for the biochemical identification of MAMPs. To overcome this, we used the free-living nematode *Caenorhabditis elegans*, leveraging the conserved nature of MAMPs among pathogenic and non-pathogenic organisms. Using *Arabidopsis* expressing the *GUS* reporter system driven by the promoter of the immunity marker gene *CYP71A12* (*pCYP71A12:GUS*) (*22*), we explored responses triggered by *C. elegans* extracts. To exclude potential contamination of bacterial MAMPs, we employed a *cerk1 fls2 pCYP71A12:GUS* line. Notably, *C. elegans* extracts induced *GUS* expression in the root elongation zone (Fig. 1A), similar to the responses triggered by the MAMP flg22 and *Meloidogyne incognita* extracts as well as its infection (*22, 23*). Additionally, we observed root pigmentation (Fig. 1B), as well as lignin accumulation (fig. S1), both hallmarks of the immune responses in resistant plants under nematode attack (*17, 24*). RNA-seq analysis revealed that *C. elegans* extracts up-regulated a significant number of genes in the *efr fls2 cerk1* triple PRR mutant, many of which were also induced by *M. incognita* extracts (Fig. 1C and Tables, S1 and S2), suggesting a shared response. Gene Ontology (GO) enrichment analysis showed that these up-regulated genes were mostly associated with immunity-related pathways (Table S3). In addition, *C. elegans* extracts induced over half of the genes activated by flg22 (*25*) or chitin (*26*), highlighting transcriptional overlaps among these treatments (Fig. 1D and Table S4). These findings demonstrate the existence of potential nematode MAMP(s) in *C. elegans* extracts.

**Fig. 1.**
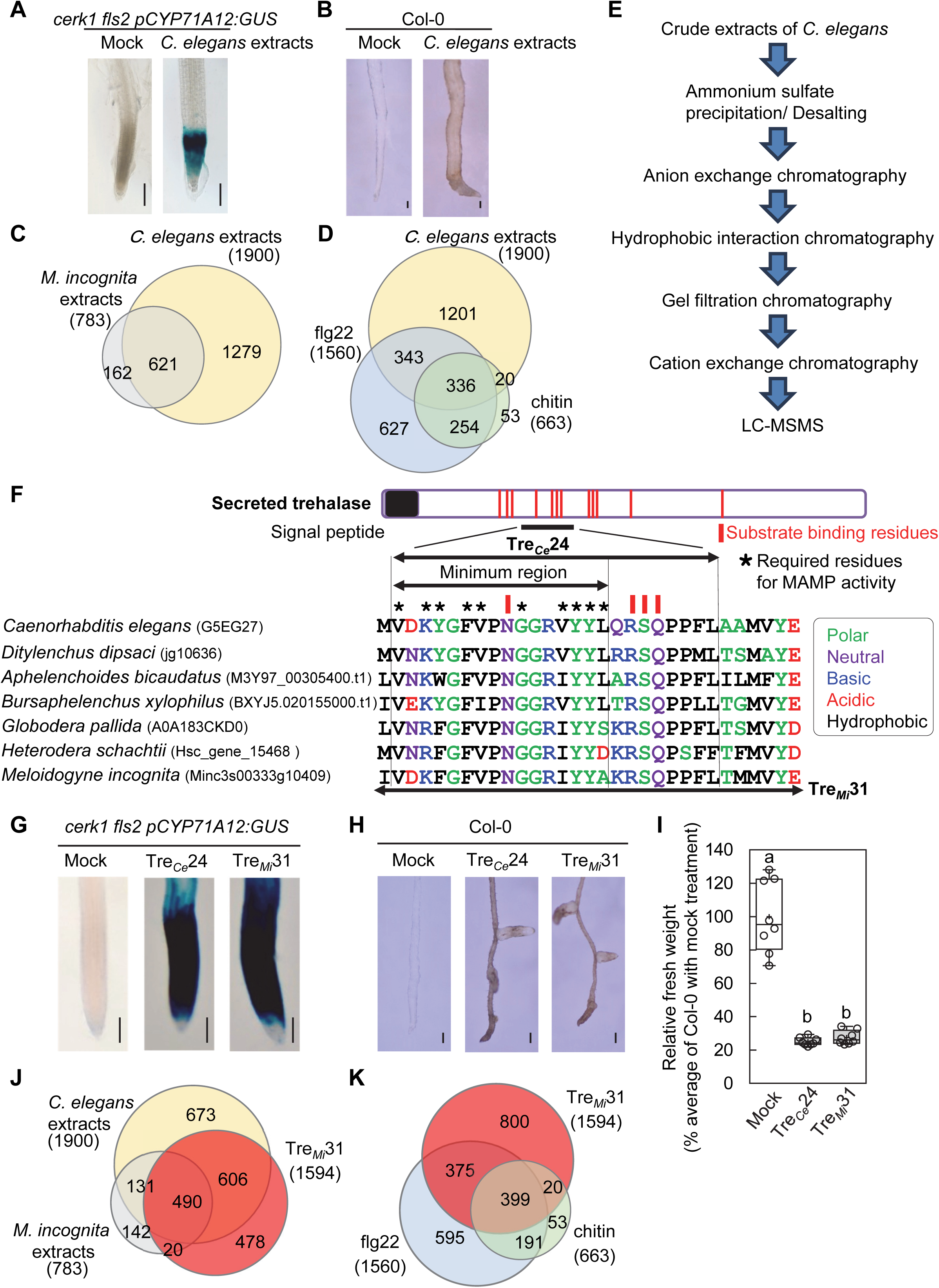
Purification and characterization of MAMPs from *Caenorhabditis elegans*. (**A** and **B**) *C. elegans* extracts (100 µg/mL) trigger MAMP-inducible responses in *Arabidopsis* Col-0, including *CYP71A12* expression (**A**) and root pigmentation (**B**). *CYP71A12* expression was assessed using the *cerk1 fls2 pCYP71A12:GUS* reporter line. Black bars represent 100 µm. Experiments were repeated more than three times with consistent results. (**C**), Venn diagram showing overlapping RNA-seq data of genes upregulated (logFC ≥ 1, adjusted p-value ≤ 0.01) in *efr fls2 cerk1* triple mutants treated with *C. elegans* and *Meloidogyne incognita* extracts (100 µg/mL) for 12 hours. (**D**) *C. elegans* extracts induce transcription of genes responsive to flg2*2* and chitin. Venn diagrams show *Arabidopsis* genes upregulated by *C. elegans* extracts, flg22(*25*), and chitin (*26*). (**E**) Workflow for MAMP identification from *C. elegans* extracts via chromatography and LC-MS/MS. (**F**) Sequence alignment of trehalase peptides from *C. elegans* and PPNs. Red lines highlight substrate-binding residues. Stars indicate essential residues for MAMP activity, determined via the *CYP71A12* expression assay (see Extended Data Fig. 3). Amino acids are color-coded based on their chemical properties. (**G** to **I**) 50 µM Tre*_Ce_*24 and Tre*_Mi_*31 peptides induce *CYP71A12* expression (**G**), root pigmentation (**H**), and seedling growth inhibition (**I**) in Col-0. Black bars in (**G** and **H**) represent 100 µm. Box plots show the 25th to 75th percentile range, with the median represented by a central line, the mean by a black cross, and whiskers indicating the range. Each open circle represents one data point from 8 samples (with one seedling per sample). Different letters denote significant differences (p ≤ 0.00001, one-way ANOVA with Tukey’s post hoc test). Experiments were repeated three times with consistent results. (**J**) Tre*_Mi_*31 upregulates genes similarly to *C. elegans* and *M. incognita* extracts. Venn diagrams show genes upregulated in Col-0 upon treatment with 50 µM Tre*_Mi_*31 or 100 µg/mL *C. elegans* and *M. incognita* extracts (logFC ≥ 1, adjusted p-value ≤ 0.01) for 12 hours. (**K**) Tre*_Mi_*31 induces genes responsive to flg22 and chitin. Venn diagrams show the overlap of genes upregulated by Tre*_Mi_*31, flg22 (*25*), and chitin (*26*).

### MAMP purification from *C. elegans*

For biochemical identification of MAMP(s), we propagated a large quantity of *C. elegans* in liquid culture and monitored the *GUS*-inducing activity at each purification step using the *cerk1 fls2 pCYP71A12:GUS* line. The activity was initially detected in the 40-70% ammonium sulfate precipitate (Fig.1E and fig. S2). After desalting, the potential MAMP(s) was purified through a series of chromatography techniques, including anion exchange, hydrophobic interaction, gel filtration, and cation exchange chromatography. Active fractions from the final cation exchange step were analyzed by liquid chromatography-mass spectrometry (LC-MS/MS). We conducted two independent purification experiments, consistently identifying the same set of 25 proteins (Table S5). Of these, 9 proteins were selected for further analysis based on the following criteria: (i) high conservation in PPNs, (ii) isoelectric point of 5.5-8.0, consistent with the migration pattern of *GUS*-inducing activity in ion exchange chromatography, and (iii) predicted molecular mass of 20-100 kDa, aligning with the behavior of *GUS*-inducing activity in gel filtration chromatography. Based on sequence conservation among PPNs, we synthesized 19 peptides from these 9 candidate proteins to test their *GUS*-inducing activity (fig. S3). Of the 19 peptides, only one, a 24-amino-acid peptide derived from secreted trehalase (designated as Tre*_Ce_*24, Fig. 1F), induced *GUS* expression in the *cerk1 fls2 pCYP71A12:GUS* line (Fig. 1G).

### Sequence specificity of trehalase-derived peptides

Trehalase is an enzyme that catalyzes the hydrolysis of trehalose, a disaccharide, into glucose molecules. *C. elegans* has five trehalase-encoding genes (*CeTRE1* to *CeTRE5*, fig. S4). The peptide Tre*_Ce_*24 is derived from *Ce*TRE3, and multiple specific peptides from *Ce*TRE3 were identified through LC-MS/MS analysis (fig. S4 and Table S6). Tre*_Ce_*24 contains key residues essential for substrate binding and is highly conserved among PPNs (Fig. 1F and fig. S5). Peptides with longer C-terminal sequences (Tre*_Ce_*26 and Tre*_Ce_*32), demonstrated stronger activity, while those with shorter C-terminal sequences (Tre*_Ce_*16 and Tre*_Ce_*15) exhibited reduced or no *GUS* expression, indicating the importance of the glutamine (17^th^) and leucine (16^th^) residues (fig. S6A). Similarly, peptides with truncated N-terminal sequences (Tre*_Ce_*24-2, Tre*_Ce_*24-4, Tre*_Ce_*24-6, and Tre*_Ce_*24-8) also showed reduced activity (fig. S6B), highlighting the critical role of the N-terminal valine, aspartic acid, lysine, and tyrosine residues. These results establish that the sequence VDKYGFVPNGGRVYYL represents the minimal region required for MAMP activity, with peptides containing a longer C-terminus exhibiting higher activity. Alanine scanning of the Tre*_Ce_*19 sequence further identified 10 essential residues for *GUS* expression, demonstrating clear sequence specificity (Fig. 1F and fig. S6C). Importantly, corresponding peptides from homologous secreted trehalases in parasitic species, including *Ditylenchus dipsaci* (stem and bulb nematode), *Aphelenchoides bicaudatus* (mycetophagous nematode), *Bursaphelenchus xylophilus* (pine wood nematode), *Globodera pallida* and *Heterodera schachtii* (CNs), as well as *M. incognita* and *M. enterolobii* (RKNs), all exhibited *GUS*-induction activity (Fig. 1G and figs. S5 and S7A). We selected *M. incognita*-derived peptide for further experiments due to its economic importance in crop production. The region of Tre*_Mi_* corresponding to the minimal MAMP-active region of Tre*_Ce_* (Tre*_Mi_*16, VDKFGFVPNGGRIYYA) showed weak *GUS*-induction activity, while the longer one (Tre*_Mi_*31) exhibited stronger response (fig. S7B), mirroring sequence specificity of peptides from *C. elegans*. The *M. incognita* genome possesses seven trehalase-encoding genes, one of which encodes both a signal peptide and the Tre*_Mi_*31 sequence (fig. S8). Another gene *Minc3s00692g16151* also encodes the Tre*_Mi_*31 sequence but lacks a signal peptide, suggesting that Arabidopsis may recognize peptides derived from both secreted and non-secreted trehalases, potentially released during infection.

### Tre*_Mi_*31 activates a series of MAMP responses

Next, we examined plant responses induced by Tre*_Mi_*31. Tre*_Mi_*31 induced root pigmentation and growth abnormalities, including root tip bending, lateral root formation, and inhibition of primary root growth, which became evident approximately 7 days after treatment. These phenotypes resemble those triggered by the phytocytokine SCOOP12 (*27*), which is recognized by the PRR MIK2 (*28, 29*) (Fig. 1H). In addition, Tre*_Mi_*31 induced lignin accumulation and inhibited seedling growth in a manner comparable to *C. elegans* extracts, Tre*_Ce_* 24, known MAMPs (*30*), and nematode culture filtrates (*19*) (fig. S9 and Fig. 1I). RNA-seq analysis revealed that Tre*_Mi_*31 upregulated 1,594 genes (Table S7), including a significant portion of the genes induced by *C. elegans* and *M. incognita* extracts (Fig. 1J). Furthermore, Tre*_Mi_*31 activated more than half of the genes upregulated by flg22 (*25*) and chitin (*26*) (Fig. 1K). GO enrichment analysis showed that the upregulated genes were predominantly associated with immunity, phytoalexin biosynthesis, and cell wall thickening, particularly lignin biosynthesis, while downregulated genes were linked to root growth and development (Table S8). In addition, Kyoto Encyclopedia of Genes and Genomes pathway analysis of genes differentially expressed in response to Tre*_Mi_*31, but not to flg22 (*31*), revealed activation of pathways involved in phenylalanine, tyrosine, and tryptophan biosynthesis, as well as phenylpropanoid biosynthesis (fig. S10, A and B), consistent with the observed lignin accumulation (fig. S10C). Moreover, Tre*_Mi_*31 strongly activated genes such as the immunity marker *CYP71A12* which is involved in the biosynthesis of tryptophan-derived secondary metabolites such as camalexin, indole glucosinolates, and 4-hydroxy indole-3-carbonyl nitrile (*32*) (fig. S10D).

### LecRK-V.5 and LecRK-V.6 are required for Tre*_Mi_*31 signaling

To elucidate the molecular mechanisms involved in Tre*_Mi_*31 recognition, we first investigated if known signaling components, including the nematode receptor NILR1 and the co-receptors such as BAK1, BKK1, and SOBIR1, were required. Tre*_Mi_*31-induced responses remained unchanged in the *nilr1-2* (*19*)*, bak1-5 bkk1* (*33, 34*), and *sobir1-13* (*35*) *mutants* (fig. S11). Then, we assessed root pigmentation and abnormality as well as seedling growth inhibition across various *Arabidopsis* accessions, identifying several accessions insensitive to Tre*_Mi_*31 (figs. S12 and S13). Among them, Cvi-0 failed to induce root pigmentation and abnormality, seedling growth inhibition, and *CYP71A12* expression in response to Tre*_Mi_*31 (Fig. 2, A to C), suggesting that Tre*_Mi_*31 recognition or its downstream signaling is specifically disrupted in this accession. A cross between Col-0 and Cvi-0 showed that all F1 plants responded to Tre*_Mi_*31, and the F2 population exhibited a 3:1 segregation ratio, indicating that a single dominant locus in Col-0 controls the response to Tre*_Mi_*31. To identify this locus, we collected 200 non-responsive F2 seedlings and performed whole-genome bulk sequencing (Fig. 2D). Single nucleotide polymorphism (SNP) analysis revealed significant enrichment of Cvi-0-derived SNPs at the distal end of chromosome 3 (fig. S14). Further phenotyping of 365 recombinant inbred lines (RILs) between Col-0 and Cvi-0 identified the locus *c3_22147* as most closely linked to the causal gene, along with the nearby locus *c3_20729* (*36*) (Table S9), aligning with the chromosomal region identified by bulk sequencing. While most RILs containing Cvi-0 alleles at these loci were insensitive to Tre*_Mi_*31 and those with Col-0 alleles were sensitive, a few exceptions arose due to homologous recombination between these loci. SNP analysis of these exceptional RILs via Sanger sequencing further narrowed the genomic region to 78 Kb (Table S10), containing 20 genes. Among these, four potential candidates were identified, all encoding highly similar lectin receptor kinases: LecRK-V.5, LecRK-V.6, LecRK-V.7, and LecRK-V.8 (fig. S15). T-DNA insertion mutants for candidate genes were tested (Table S11), and the mutant SALK_133163 (*lecrk-V.5-3*) (*37*), which has a T-DNA insertion in the exon of *LecRK-V.5*, exhibited reduced root pigmentation (Fig. 2D). Notably, *LecRK-V.5* is transcriptionally upregulated by Tre*_Mi_*31 (fig. S16A), a characteristic feature of PRRs, and shows strong co-expression with *NILR1* (fig. S16B). In contrast, T-DNA insertions in *LecRK-V.6*, *LecRK-V.7*, and *LecRK-V.8* did not affect root pigmentation, seedling growth inhibition, or *CYP71A12* expression upon Tre*_Mi_*31 treatment (fig. S17). These findings suggest that *LecRK-V.5* plays a major role in Tre*_Mi_*31-induced responses.

**Fig. 2.**
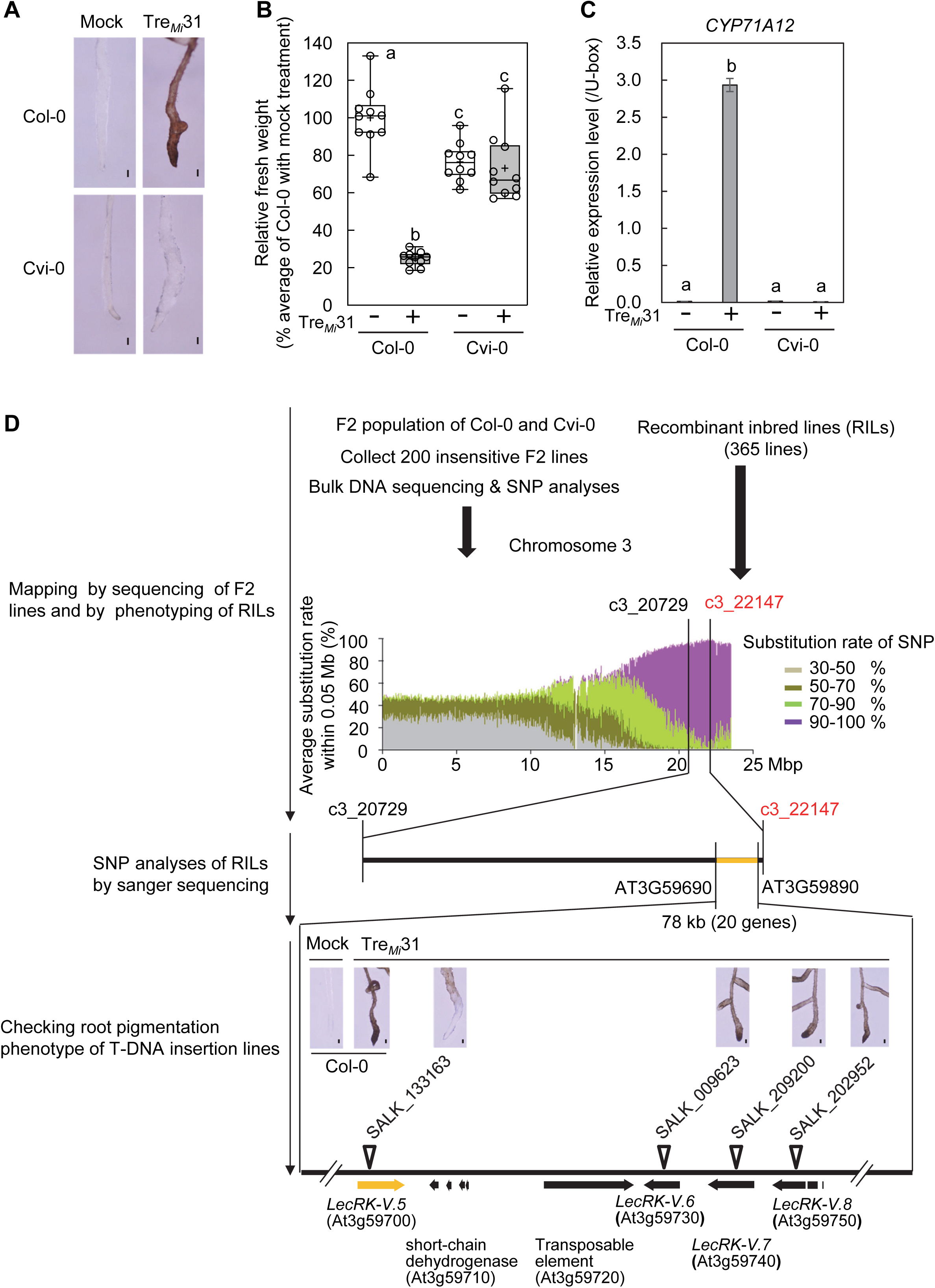
Identification of the genomic region conferring Tre*_Mi_*31 insensitivity in Cvi-0. (**A** to **C**) *Arabidopsis* accession Cvi-0 is insensitive to Tre*_Mi_*31, unlike Col-0. Col-0 seedlings, but not Cvi-0, display root pigmentation (**A**), growth inhibition (**B**), and *CYP71A12* expression (**C**) in response to 50 µM Tre*_Mi_*31. Black bars in (**A**) represent 100 µm. In the box plot (**B**), the 25th**-**75th percentiles are shown, with the median indicated by a central line, the mean by a black cross, and whiskers representing the full range. Each open circle represents one data point from 10 samples (with one seedling per sample). Different letters denote significant differences (p ≤ 0.01, one-way ANOVA with Tukey’s post hoc test). In the bar chart (**C**), transcript levels of *CYP71A12* in Col-0 and Cvi-0 seedlings with or without treatment with 50 µM Tre*_Mi_*31 for 6 hours were measured by RT-qPCR after normalization to the *U-box* housekeeping gene transcript (*AT5G15400*). Values are presented as mean ± standard error (SE) of three technical replicates, with different letters indicating significant differences (p ≤ 0.05, one-way ANOVA with Tukey’s post hoc test). Experiments were repeated three times with consistent results. (**D**) Genomic sequencing of F2 populations with insensitivity from Col-0 × Cvi-0 crosses, along with phenotyping of recombinant inbred lines (RILs), identified a genomic region linked to Tre*_Mi_*31 insensitivity in Cvi-0. Significant accumulation of SNPs from Cvi-0 was detected near the end of chromosome 3 (fig. S14). The substitution ratio of SNPs (Col-0 to Cvi-0) was categorized into four levels (30-50%, 50-70%, 70-90%, and 90-100%) and color-coded accordingly. The vertical axis represents the average substitution ratio of SNPs (Cvi-0 / (Col-0 + Cvi-0)) within a 0.05 Mb window. RIL phenotyping identified C3_22147 as the closest marker and C3_20729 as the second closest marker to the causative gene. Sanger sequencing narrowed the candidate region to 78 kb between At3g59690 and At3g59890, containing 20 genes (Table S10). Screening T-DNA insertion lines revealed that the *lecrk-V.5* mutant (SALK_133163: *lecrk-V.5-3*) shows reduced sensitivity to 50 µM Tre*_Mi_*31 (Table S11). Black bars represent 100 µm.

To confirm the role of *LecRK-V.5* in Tre*_Mi_*31 signaling, we examined the phenotypes of two mutant alleles, *lecrk-V.5-3* and *lecrk-V.5-2* (GK_623G01) (*38*) (fig. S18, A and B). The *lecrk-V.5-2* mutant displayed an almost complete loss of sensitivity to Tre*_Mi_*31, while *lecrk-V.5-3* exhibited a partial response, characterized by reduced root pigmentation, restored seedling growth inhibition, and diminished *CYP71A12* expression (Fig. 3, A to C, and fig. S18C). In contrast, Col-0, Cvi-0, *lecrk-V.5-2, and lecrk-V.5-3* mutants induce root pigmentation in response to SCOOP12 (Fig. 3A). Notably, *lecrk-V.5-2* contains a T-DNA insertion in the 3’ half of the gene, whereas *lecrk-V.5-3* has an insertion in the 5’ end (fig. S18A). Interestingly, *lecrk-V.5-2* showed higher expression of the 5’ region, encoding the ectodomain with the transmembrane region but lacking the kinase domain (fig. S18D). This truncated form may act in a dominant-negative manner, potentially through heterodimerization with other LecRKs or competition for ligand binding, whereas *lecrk-V.5-3* is a true knockout mutant. RNA-seq analysis further demonstrated that nearly all Tre*_Mi_*31-induced genes were downregulated in the *lecrk-V.5-3* mutant (Fig. 3D), indicating that *LecRK-V.5* is important for the recognition or very early stages of the Tre*_Mi_*31 signaling pathway (Table S12). This conclusion was further supported by the restoration of Tre*_Mi_*31 responses in the *lecrk-V.5-2* mutant following complementation with *LecRK-V.5-3xHA* under its native promoter, with the complementation lines exhibiting even stronger responsiveness, likely due to higher expression levels (Fig. 3, E and F, and fig. S18E). To further explore the role of *LecRK-V.5* in downstream signaling events, we compared the effects of Tre*_Mi_*31 and flg22 on MAPK activation. Tre*_Mi_*31 induced a slower and more prolonged MAPK activation than flg22 (Fig. 3G) and did not trigger ROS production (fig. S18F), suggesting that Tre*_Mi_*31 induces downstream pathways that are, at least partially, distinct from most identified MAMP receptors.

**Fig. 3.**
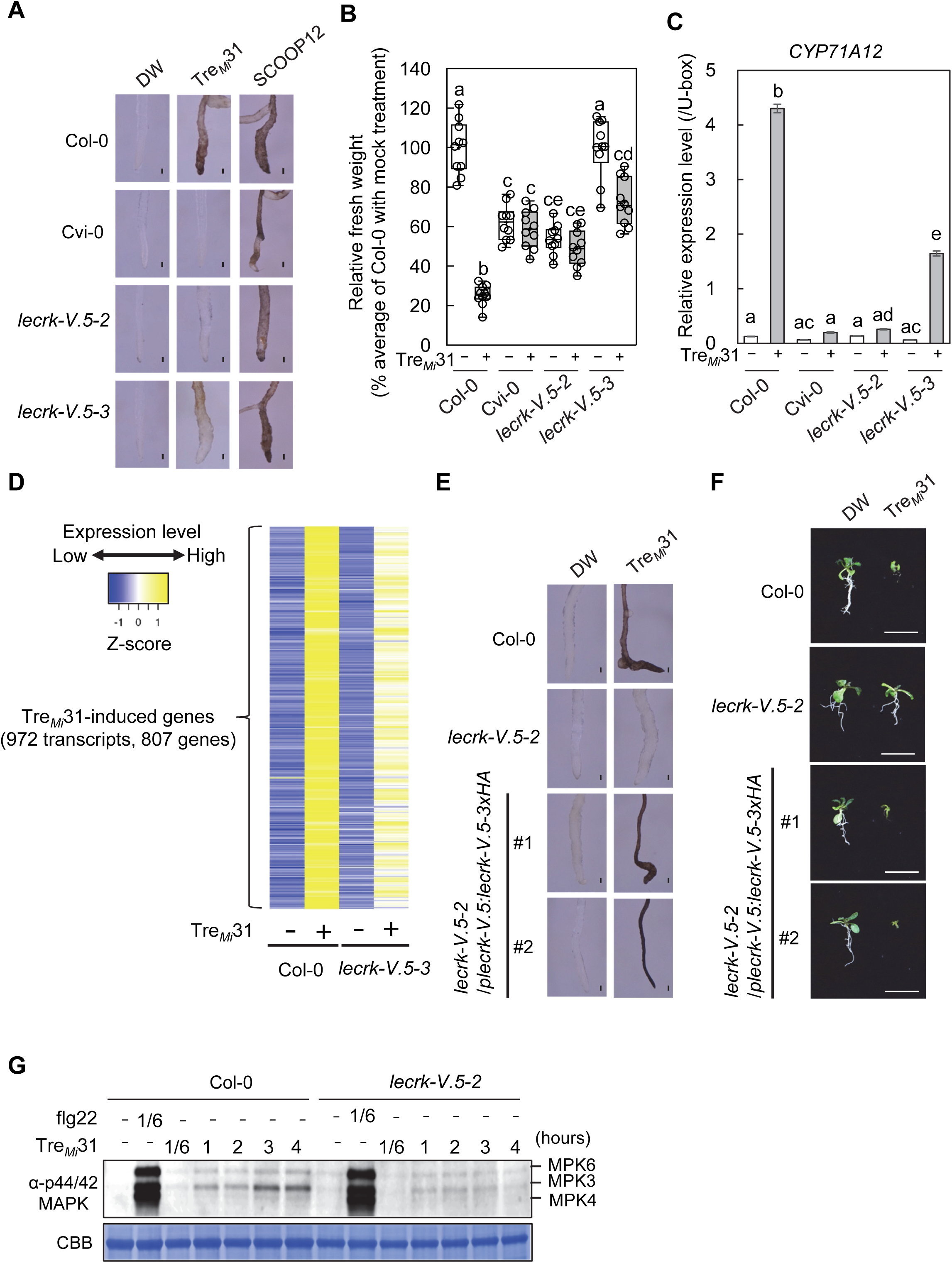
*lecrk-V.5 mutants have defects in* Tre*_Mi_*31 signaling. (**A** to **C**) *lecrk-V.5-3* mutants show reduced responses to Tre*_Mi_*31, whereas *lecrk-V.5-2* mutants do not respond. Col-0 seedlings, but not Cvi-0 or *lecrk-V.5-2* mutants, exhibit root pigmentation (**A**), growth inhibition (**B**), and *CYP71A12* expression (**C**) in response to 50 μM Tre*_Mi_*31, while Col-0, Cvi-0, *lecrk-V.5-2, and lecrk-V.5-3* mutants induce root pigmentation in response to 2 μM SCOOP12. Black bars in (**A**) represent 100 µm. In the box plot (**B**), the 25th**-**75th percentiles are shown, with the median indicated by a central line, the mean by a black cross, and whiskers representing the full range. Each open circle represents one data point from 10 samples (with one seedling per sample). Different letters denote significant differences (p ≤ 0.01, one-way ANOVA with Tukey’s post hoc test). In the bar chart (**C**), transcript levels of *CYP71A12* in the seedlings with or without treatment with 50 µM Tre*_Mi_*31 were measured by RT-qPCR after normalization to the *U-box* housekeeping gene transcript (*AT5G15400*). Values are presented as mean ± SE of three technical replicates, with different letters indicating significant differences (p ≤ 0.05, one-way ANOVA with Tukey’s post hoc test). Experiments were repeated three times with consistent results. (**D**) RNA-seq analysis reveals suppression of most upregulated genes (logFC ≥ 1, adjusted p-value ≤ 0.01) by the treatment with 30 µM Tre*_Mi_*31 for 12 hours in *lecrk-V.5-3* mutants compared to Col-0. (**E** and **F**) The expression of *LecRK-V.5-3xHA* under its native promoter restores root pigmentation (**E**) and seedling growth inhibition (**F**) in *lecrk-V.5-2* mutants upon treatment with 50 µM Tre*_Mi_*31. T2 complementation lines with the transgene were used for the assay. Black bars in (**E**) represent 100 µm. White bars in (**F**) represent 1 cm. Experiments were repeated three times with consistent results. (**G**) Col-0 seedlings, but not *lecrk-V.5-2* mutants, show slow and sustained activation of MAPKs upon 50 µM Tre*_Mi_*31 treatment. 1 μM flg22 was used as a positive control. Experiments were repeated three times with consistent results.

As the *lecrk-V.5-3* mutant did not completely abolish the Tre*_Mi_*31 response, we hypothesized that the other LecRK-V(s) might also contribute to the signaling pathway. To investigate this, we generated CRISPR-mediated *lecrk-V.5-3/lecrk-V.78-d* mutants, which lack the region between *LecRK-V.7* and *LecRK-V.8* in the *lecrk-V.5-3* background (fig. S19). These mutants exhibited a weak yet noticeable response to Tre*_Mi_*31 including root pigmentation and shoot growth inhibition, suggesting a role for *LecRK-V.6* in Tre*_Mi_*31 recognition (Fig. 4A and fig. S20A). In contrast, *lecrk-V.8/lecrk-V.56-d* (lacking regions from *LecRK-V.5* to *LecRK-V.6* in *lecrk-V.8* background) and *lecrk-V.567-d* (spanning *LecRK-V.5* to *LecRK-V.7*) showed no response to Tre*_Mi_*31, indicating that *LecRK-V.7* and *LecRK-V.8* are not involved in Tre*_Mi_*31 recognition (Fig.4, A to C, and fig. S20A). Additionally, *lecrk-V.567-d* did not respond to Tre*_Ce_*24 and trehalase-derived peptides from PPNs (fig. S20, B and C), suggesting that *LecRK-V.5* and *LecRK-V.6* are required for the response to Tre*_Mi_*31 and other PPN trehalase-derived peptides.

**Fig. 4.**
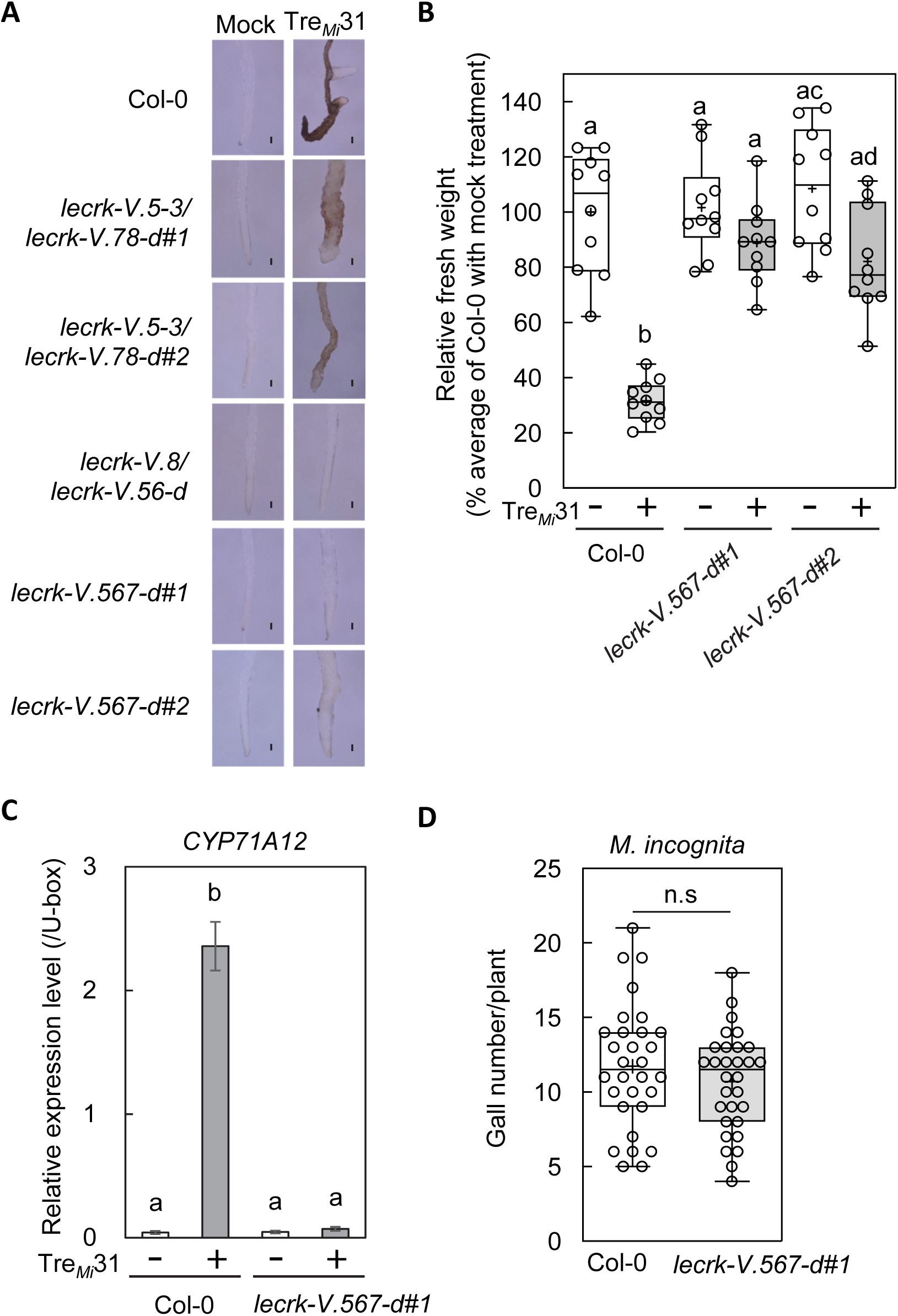
*LecRK-V.5* and *LecRK-V.6* are involved in Tre*_Mi_*31 recognition. (**A**) CRISPR deletion line, *lecrk-V.5-3/lecrk-V.78-d* shows reduced but detectable root pigmentation upon 50 µM Tre*_Mi_*31 treatment, while *lecrk-V.567-d* and *lecrk-V.8/lecrk-V.56-d* mutants do not exhibit root pigmentation. Black bars represent 100 µm. (**B** and **C**) *lecrk-V.567-d* mutants do not show seedling growth inhibition (**B**) and *CYP71A12* expression (**C**) upon treatment with 50 µM Tre*_Mi_*31. In the box plot (**B**), the 25th**-**75th percentiles are shown, with the median indicated by a central line, the mean by a black cross, and whiskers representing the full range. Each open circle represents one data point from 10 samples (with one seedling per sample). Different letters denote significant differences (p ≤ 0.05, one-way ANOVA with Tukey’s post hoc test). In the bar chart (**C**), transcript levels of *CYP71A12* in the seedlings upon treatment with 50 µM Tre*_Mi_*31 for 6 hours were measured by RT-qPCR after normalization to the *U-box* housekeeping gene transcript (*AT5G15400*). Values are presented as mean ± SE of three biological replicates, with different letters indicating significant differences (p ≤ 0.0001, one-way ANOVA with Tukey’s post hoc test). Experiments were repeated three times with consistent results. (**D**) Number of galls formed in the *lecrk-V.567-d* line upon infection with *M. incognita*. No significant (n.s.) difference was observed compared to Col-0 plants by Student’s t-test (n=30). Experiments were repeated three times with consistent results.

Phylogenetic analysis of their ectodomains reveals that LecRK-V.5 and LecRK-V.6, along with the closely related LecRK-Vs (3, 4, 7, 8), are exclusive to the Brassicales (fig. S21), indicating that the recognition system for trehalase-derived peptides by LecRK-Vs is unique to this lineage. To evaluate whether Tre*_Mi_*31 recognition contributes to resistance against PPNs, we compared the number of galls formed by *M. incognita* in Col-0 and *lecrk-V.567-d* mutants. No significant difference was observed (Fig. 4D), suggesting that Tre*_Mi_*31 recognition plays only a modest role in resistance. This limited contribution may be due to the lower expression of secreted trehalases in *M. incognita* during infection compared to CNs (*39*) or the ability of *M. incognita* to overcome Tre*_Mi_*31-mediated defenses through virulence effectors.

### LecRK-V.5 and LecRK-V.6 are required for the induction of immune responses by trehalase-derived peptides from pathogenic insects and fungi

Given the high conservation of secreted trehalases across higher organisms, we conducted a BLAST search to identify sequences similar to Tre*_Mi_*31 in other plant pathogenic organisms. Remarkably, several insects, fungal, and oomycetes pathogens, and were found to possess sequences corresponding to Tre*_Mi_*31 (Fig. 5A and fig. S22). We synthesized these peptides and evaluated their MAMP activity by using the *cerk1 fls2 pCYP71A12:GUS* line. The results showed clear MAMP activity in peptides derived from the fungal pathogen *Colletotrichum fructicola* and plant-pathogenic insects, including aphids (*Aphis glycines* and *Acyrthosiphon pisum*) and the Mediterranean fruit fly (*Ceratitis capitata*) (fig. S23A). Sequence comparison revealed that the MAMP-active peptides tended to have more positively charged residues (K, R, H) at the fourth position from the N-terminus, while non-active peptides were more enriched in negatively charged residues (E) (Fig. 5A). Consistently, substituting positively charged residues with negatively charged ones in Tre*_Ce_*19 and Tre*_Mi_*19 abolished their MAMP activity (fig. S23, B and C). Conversely, introducing a reverse-charge mutation (negative to positive) in the non-active peptide from the oomycete pathogen *Albugo laibachii* activated its MAMP activity (Fig. 5B), underscoring the importance of the positively charged residue at this position. Furthermore, the *lecrk-V.567-d* mutant failed to induce root pigmentation and *CYP71A12* expression in response to peptides from insects and fungal pathogens, highlighting the essential role of *LecRK-V.5* and *LecRK-V.6* in mediating broad-spectrum pathogen recognition through trehalase-derived peptides (Fig. 5, C and D, and fig. S23D).

**Fig. 5.**
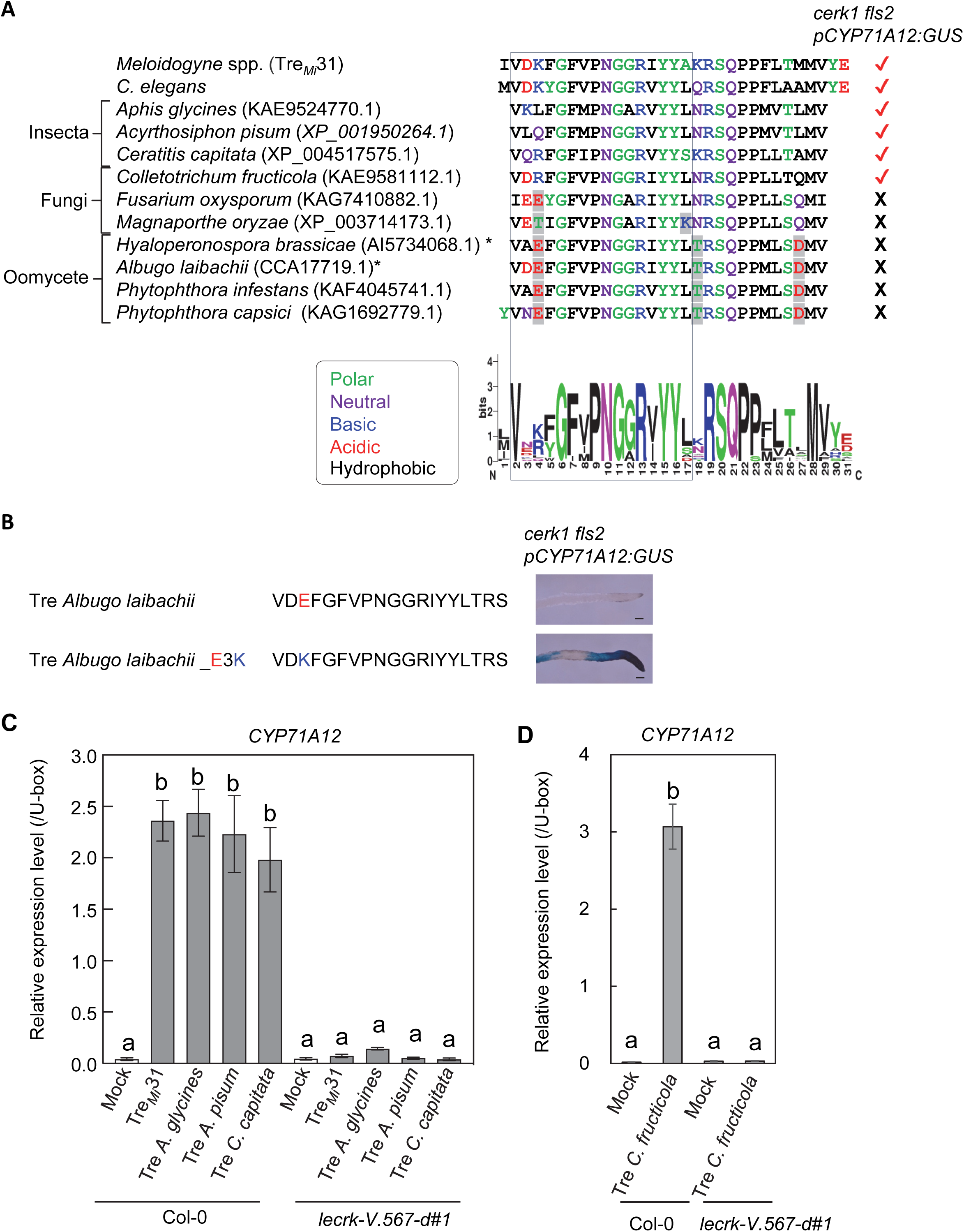
*LecRK-V.5* and *LecRK-V.6* are involved in the recognition of trehalase peptides from pathogenic insects and fungi. (**A**) Sequence alignment of trehalase peptides from *C. elegans*, *M. incognita*, plant pathogenic insects, fungi, and oomycetes, with their MAMP activities determined via the *CYP71A12* expression assay (see Extended Fig. 9A). The sequence logo highlights conserved residues in MAMP-active peptides from nematodes (Extended Fig. 4A), insects, and fungi. Trehalases in *H. brassicae* (AI5734068.1) and *A. laibachii* (CCA17719.1) do not have signal peptides predicted by SignalP-6.0 (**\***). Amino acid residues highlighted in gray differ in properties from those found in MAMP-active peptides. (**B**) A charge-reverse substitution of the third glutamic acid residue to lysine (E3K) in a MAMP-inactive trehalase peptide from *A. laibachii* restores MAMP activity. *CYP71A12* expression was analyzed upon treatment with 50 µM peptide in the *cerk1 fls2 pCYP71A12:GUS* line. Black bars represent 100 µm. (**C** and **D**) Col-0 seedlings, but not the *lecrk-V.567-d* line, show *CYP71A12* expression in response to trehalase-derived peptides from insects (**C**) and fungi (**D**). Transcript levels of *CYP71A12* in the seedlings upon treatment with 50 µM Trehalase-derived peptides were measured by RT-qPCR after normalization to the *U-box* housekeeping gene transcript (*AT5G15400*). Values are presented as mean ± SE of three biological replicates, with different letters indicating significant differences (p ≤ 0.0001, one-way ANOVA with Tukey’s post hoc test). Experiments were repeated three times with consistent results.

## Discussion

This study elucidates how plants recognize PPNs through trehalase-derived peptides, identifying *LecRK-V.5* and *LecRK-V.6* as essential components of this immune signaling pathway (fig. S24). The conservation of this peptide region in trehalases across PPNs, as well as in insects and fungal pathogens, underscores the significant role of LecRK-Vs in response to broad-spectrum pathogens.

Trehalose plays diverse and essential roles in various organisms, including bacteria, fungi, nematodes, and insects. It serves as an energy and carbon source and acts as a stress protectant, stabilizing proteins and membranes under extreme conditions such as heat, cold, desiccation, and osmotic stress (*40*). Trehalose is particularly critical for pathogenic organisms, as mutants of bacterial pathogens (*41, 42*) and fungal pathogens (*43*) defective in trehalose biosynthesis exhibit significantly reduced virulence in plants, suggesting its possible protective effects against host immunity. Pathogenic organisms also induce trehalose accumulation in the host by activating host trehalose biosynthesis pathways (*41, 44*). Enhanced host trehalose biosynthesis increases susceptibility to oomycete pathogens, while inhibition of trehalose biosynthesis suppresses disease (*44*), highlighting trehalose’s role as a susceptibility factor. Accumulated trehalose may support pathogen survival under host immune defenses and provide energy during parasitism (*45*), facilitating the development and colonization of pathogenic organisms. Similarly, PPNs induce trehalose accumulation in specialized feeding structures, such as galls (*46*) and syncytia (*47*), likely aiding their growth and survival. Notably, transgenic soybean hairy roots overexpressing non-secreted trehalase, potentially reducing trehalose levels, showed increased resistance to CNs, emphasizing the importance of trehalose levels in plant tissues in determining susceptibility to nematode infections (International patent publication number: WO2008095919A1).

Consistently, trehalase enzymes are crucial for the survival of many microorganisms, serving diverse roles essential for their physiology and pathogenicity. The significance of trehalases in microbial survival is highlighted by their widespread use as targets in agricultural fungicides and insecticides (*48*). For example, the trehalase inhibitor validamycin is commonly employed to control rice sheath blight caused by *Rhizoctonia solani* (*48*). Trehalases are also implicated in virulence and parasitism of PPNs. Host-induced gene silencing of non-secreted trehalase in *M. incognita* within transgenic tobacco significantly reduces egg production and root-knot formation (*49*), suggesting a critical role for non-secreted trehalase in parasitism, potentially through its involvement in chitin biosynthesis required for eggshell formation.

While the roles of non-secreted trehalases in parasitism are well-documented, the function of secreted trehalases remains less explored. Secreted trehalases are conserved across diverse taxa, including bacteria, fungi, oomycetes, nematodes, and insects, suggesting important roles for their survival. Interestingly, pathogenic *Candida* species secrete trehalases during host colonization, whereas their mammalian hosts do not produce trehalose (*50–53*). Knockout mutants of *Candida* trehalases show significantly reduced infectivity, demonstrating their essential role in virulence. These secreted trehalases likely exploit trehalose released from other microbial sources or dying cells within the host environment, providing a readily accessible nutrient and energy source during infection.

In PPNs, secreted trehalases are released during infection to utilize host-derived trehalose. For example, in pine wood nematodes, secreted trehalase levels in the secretome increase upon treatment with pine extract (*54, 55*). In CNs, a secreted trehalase (encoded by *Hsc_gene_15468*) has been identified as a virulence effector candidate, produced in subventral gland cells and most probably secreted into the apoplast during the early stages of infection (*56–58*). Notably, *Hsc_gene_15468* transcripts accumulate significantly in gland cells during the migratory, parasitic second-stage juvenile (J2) phase, with expression levels about 14 thousand times higher than in the cyst phase and 14 times higher than in the pre-parasitic J2 phase (*57, 58*). This higher expression of secreted trehalase during the early infection stage in CNs, coupled with the recognition of secreted trehalases by *Arabidopsis*, suggests an important role for these enzymes in nematode survival and parasitism within host plants, while the precise role of apoplastic trehalose in plant-PPN interactions remains unclear.

Some LecRKs function as immune receptors, recognizing MAMPs and damage-associated molecular patterns (DAMPs). For instance, DORN1 (LecRK-I.9) recognizes extracellular ATP (*59*), while LecRK-VI.2, is a potential receptor for extracellular NAD^+^ and NAD^+^ phosphate (*60*). Similarly, LecRK-VI.8 may be involved in the recognition of eNAD^+^ but not eNADP^+^ (*61*). Additionally, LecRK-I.1 and LecRK-I.8 are implicated in the perception of insect eggs, although the specific egg-derived ligands remain unidentified (*62, 63*). Our findings indicate that LecRK-V.5 and LecRK-V.6 participate in immune responses to trehalase-derived peptides from PPNs, insect parasites, and fungal pathogens. This suggests secreted trehalases are highly conserved MAMPs across species, while they may also function as virulence effectors in some pathogenic organisms like CNs. Notably, *LecRK-V.5* is transcriptionally upregulated by Tre*_Mi_*31 (fig. S16A) and is closely co-expressed with *NILR1* (fig. S16B). Moreover, *LecRK-V.5* is transcriptionally upregulated in *Arabidopsis* tissues surrounding CNs during their migratory stage (*19*), aligning well with the significant upregulation of secreted trehalase (*Hsc_gene_15468*) during this phase. These findings suggest a possibility that LecRK-V.5 and LecRK-V.6 may function as receptors or components of receptor complexes for trehalase-derived peptides, though further studies are needed to confirm direct interactions. Another possibility is that the trehalase MAMP is directly recognized by another receptor, such as an LRR-RLK that often binds to peptide ligands, and LecRK-V.5 and LecRK-V.6 are activated as downstream components. Since the recognition of trehalase-derived peptides by LecRK-V.5 and LecRK-V.6 is unique to Brassicales (fig. S21), this discovery opens the possibility of transferring this signaling mechanism to other plant species, providing a promising strategy to enhance immunity against diverse pathogens in economically important crops (*64*).

## Materials and Methods

### Plant materials and growth conditions

Seedlings of *A. thaliana* (L.) Heynh. were grown on a half-strength MS medium containing 1% sucrose with or without 0.8% agarose under a continuous light photoperiod or a long-day photoperiod (16 hours light and 8 hours dark) at 23°C. The light is provided by light-emitting diodes (65-75 μE m−2 s−1 for Arabidopsis). The humidity was maintained at 60-70%.

### Extraction of *C. elegans* and *M. incognita* for MAMP activity assays

*C. elegans* and *M. incognita* were dissolved in the Extraction Buffer (20 mM MES, pH 5.6, 0.1% CHAPS (FUJIFILM Wako Pure Chemical Corporation, Osaka, Japan), and proteins were extracted by sonication. The crude extract was centrifuged at 12,000 g for 10 min and repeated more than three times to remove debris. The protein concentration was determined by Bradford protein assay (Bio-Rad, Hercules, CA). An equivalent volume of the extraction buffer was treated to the samples as mock control.

### GUS assay of cerk1 fls2 pCYP71A12:GUS line

GUS staining was performed on 10-to 14-day-old seedlings of the *cerk1 fls2 pCYP71A12:GUS* reporter line after treatment with extract or peptides for 24 hours. Seedlings were vacuum infiltrated with staining solution (50 mM sodium phosphate buffer, pH 7.0, 0.5% Triton X-100, 1 mM potassium ferrocyanide, 1 mM potassium ferricyanide, 5% methanol, 1 mM 5-bromo-4-chloro-3-indolyl D-Glucuronide (X-gluc)) and then incubated at 37℃ for 30 min to 1 hour.

### Seedling growth inhibition and root pigmentation assay

Arabidopsis seeds were sown on half-strength MS agar plates containing 1% sucrose and 0.8% agar. After cold treatment for 2 to 3 days at 4°C, plates were transferred to a growth chamber under continuous light at 23°C. Five-day-old seedlings were then moved to liquid half-strength MS medium with 1% sucrose and treated with extracts or peptides for 7 days. Fresh weight was measured using an analytical balance (Mettler Toledo, Zurich, Switzerland), and root pigmentation was observed under a light microscope.

### Peptide Synthesis

All peptides were synthesized by GenScript (Piscataway, NJ) with purities exceeding 70%. Trifluoroacetic acid was exchanged for phosphate. The pH of each peptide solution was checked and neutralized as necessary.

### RNA-Seq and differential gene expression analyses

Ten-day-old *Arabidopsis* seedlings of Col-0 or *efr fls2 cerk1* mutants, grown in liquid half-strength MS medium with 1% sucrose, were treated with 100 µg/mL *C. elegans* extract for 0.5, 1, 3, 6, and 12 hours (Fig. 1, C and D, and Table S1); with 100 µg/mL *M. incognita* extract for 12 hours (Table S2); and with 50 µM Tre*_Mi_*31 for 12 hours (Fig. 1, J and K, and Table S7), with three biological replicates per condition. Additionally, ten-day-old seedlings of Col-0, *lecrk-V.5-3* (Fig. 3D, Table S12) were treated with 30 µM Tre*_Mi_*31 for 12 hours with four biological replicates per genotype. Transcript levels were analyzed by RNA-Seq, with library preparation performed using the BradSeq protocol (*65*). Single-end 86-bp reads were sequenced on an Illumina NextSeq 500 platform and mapped to the Arabidopsis cDNA reference based on TAIR10 using Bowtie v0.12.9 (*66*). Read counts were obtained per transcript model (*67*) and Reads Per Million mapped reads (PPM) were calculated. Differentially expressed genes (DEGs) with a false discovery rate (FDR) ≤ 0.01 were identified using edgeR. Sequencing reads have been deposited in the DNA Data Bank of Japan (DDBJ) under accession number PRJDB19783 (BioProject: SAMD00851270-SAMD00851330).

### Identification of genomic region for Tre*_Mi_*31 insensitivity in Cvi-0 by sequencing

Cvi-0 and Col-0 were crossed to produce F1 plants, which were then used to generate an F2 population. Root pigmentation assays with 50 µM Tre*_Mi_*31 peptide were performed on the F2 population. A total of 200 F2 lines insensitive to 50 µM Tre*_Mi_*31 peptide were pooled, and their genomic DNA (gDNA) was extracted using Qiagen DNeasy Plant Mini Kits (Qiagen, Venlo, The Netherlands). For DNA library preparation, gDNA was sheared to a size of approximately 450**-**500 bp using the Covaris S2 system (Covaris, Woburn, Massachusetts), and DNA libraries were prepared using the Ovation Rapid DR Multiplex system (NuGEN Technology, Männedorf, Switzerland) according to the manufacturer’s instructions. DNA sequencing was performed with the NextSeq500 (Illumina, San Diego, CA), and potential causal SNPs were identified as previously described (*68*).

### Identification of genomic marker closest to causal mutation by RIL lines

The Cvi-0 x Col-0 RIL population was obtained from INRA Versailles (*36*). Root pigmentation assays with 50 µM Tre*_Mi_*31 peptide were performed for 365 RIL lines. The positions of genetic markers linked to the phenotype were narrowed down with the genotyping and phenotyping data of RILs (Table S9).

### T-DNA insertion lines

T-DNA insertion mutant lines, *lecrk-V.5-2* (GK_623G01) (*38*), *lecrk-V.5-3* (SALK_133163C), *lecrk-V.6* (SALK_009623C), *lecrk-V.7* (SALK_209200C), *lecrk-V.8* (SALK_202952C) and other lines listed in Table S11 were obtained from the Arabidopsis Biological Resource Center at the Ohio State University. Previously published lines were: *bak1-5 bkk1* (*33, 34*), *fls2* (*7*), *cerk1* (*11*), *nilr1-2* (*19*), *sobir1* (*35*). Homozygous *cerk1 fls2 pCYP71A12:GUS* line was generated by crossing homozygous lines and then selection by genotyping.

Other methods are provided in the Supplementary Methods.

### Statistics

Statistical significance was determined using t-tests and one-way ANOVA in GraphPad Prism6 software (GraphPad Software, San Diego, CA, USA).

## Acknowledgments

We appreciate Saki Noda, Dr. Keiko Kuwata, and Prof. Yoshikatsu Matsubayashi for supporting LC-MS/MS analyses. We thank all members of the Shirasu lab for valuable discussions. We are grateful to Ayami Furuta, Naomi Watanabe, Mamiko Kouzai, Mizuki Yamamoto, and Yoko Nagai for their assistance with this project. We also thank Prof. Shinichiro Sawa and Prof. Ryoji Shinya for technical advice and materials, and MolGen B.V. for supplying a large quantity of *C. elegans*.

## Funding

Cabinet Office, Government of Japan, cross-ministerial Strategic Innovation Promotion Program (SIP), “Technologies for Creating Next-Generation Agriculture, Forestry and Fisheries” (funding agency, Bio-oriented Technology Research Advancement Institution, NARO) (to TU and YK) MEXT KAKENHI Grant Numbers JP16H06186 (to Y.K.)

MEXT KAKENHI Grant Numbers JP16KT0037 (to Y.K.)

MEXT KAKENHI Grant Numbers JP20H02994 (to Y.K.)

MEXT KAKENHI Grant Numbers JP21K19128 (to Y.K.)

MEXT KAKENHI Grant Numbers JP24K01764 (to Y.K.)

MEXT KAKENHI Grant Numbers JP17H06172 (to K.Sh.)

MEXT KAKENHI Grant Numbers JP22H00364 (to K.Sh.)

Japan Society for the Promotion of Science KAKENHI Grant Numbers JP21J20881 (to E.I), Japan Society for the Promotion of Science KAKENHI Grant Numbers JP21F21793 (to B.P.M.N.), Japan Society for the Promotion of Science KAKENHI Grant Numbers JP20H05909 (to K.Sh.).

Japan Science and Technology Agency (JST) Program GteX (JPMJGX23B2) (to K.Sh) Japan Science and Technology Agency (JST) ASPIRE Program (JPMJAP2306) (to K.Sh) RIKEN TRIP initiative Field Omics (to K.Sh).

## Author contributions

Conceptualization: YK, KSh

Methodology: YK, KSh

Investigation: EI, YK, NM, EO, KS, NI, TS, TU

Bioinformatics analyses: BPMN, MWS, KS

Visualization: EI, YK, BPMN, KS

Funding acquisition: EI, YK, KSh

Supervision: YK, KSh

Writing—original draft: EI, YK, KSh

Writing—review & editing: EI, YK, NM, EO, KS, NI, BPMN, MWS, TS, TU, KSh

## Data and materials availability

All data generated or analyzed during this study are included in the article or supplementary materials. Sequencing reads of RNA-Seq analyses have been deposited in the DNA Data Bank of Japan (DDBJ) under accession number PRJDB19783 (BioProject: SAMD00851270-SAMD00851330). For requests for materials, contact Yasuhiro Kadota or Ken Shirasu.

## References

1. J. T. Jones et al., Top 10 plant-parasitic nematodes in molecular plant pathology. Mol Plant Pathol 14, 946–961 (2013).

2. M. R. Khan, in Novel Biological and Biotechnological Applications in Plant Nematode Management, M. R. Khan, Ed. (Springer Nature Singapore, Singapore, 2023), pp. 3–45.

3. B. Molloy, T. Baum, S. Eves-van den Akker, Unlocking the development- and physiology-altering ’effector toolbox’ of plant-parasitic nematodes. Trends Parasitol 39, 732–738 (2023).

4. S. Siddique, F. M. Grundler, Parasitic nematodes manipulate plant development to establish feeding sites. Curr Opin Microbiol 46, 102–108 (2018).

5. D. Couto, C. Zipfel, Regulation of pattern recognition receptor signalling in plants. Nat Rev Immunol 16, 537–552 (2016).

6. G. Felix, J. D. Duran, S. Volko, T. Boller, Plants have a sensitive perception system for the most conserved domain of bacterial flagellin. Plant J 18, 265–276 (1999).

7. L. Gomez-Gomez, T. Boller, FLS2: an LRR receptor-like kinase involved in the perception of the bacterial elicitor flagellin in Arabidopsis. Mol Cell 5, 1003–1011 (2000).

8. G. Kunze et al., The N terminus of bacterial elongation factor Tu elicits innate immunity in Arabidopsis plants. Plant Cell 16, 3496–3507 (2004).

9. C. Zipfel et al., Perception of the bacterial PAMP EF-Tu by the receptor EFR restricts Agrobacterium-mediated transformation. Cell 125, 749–760 (2006).

10. A. Yamada, N. Shibuya, O. Kodama, T. Akatsuka, Induction of Phytoalexin Formation in Suspension-cultured Rice Cells by N-Acetyl-chitooligosaccharides. Bioscience, Biotechnology, and Biochemistry 57, 405–409 (1993).

11. A. Miya et al., CERK1, a LysM receptor kinase, is essential for chitin elicitor signaling in Arabidopsis. Proc Natl Acad Sci U S A 104, 19613–19618 (2007).

12. K. Thor et al., The calcium-permeable channel OSCA1.3 regulates plant stomatal immunity. Nature 585, 569–573 (2020).

13. W. Tian et al., A calmodulin-gated calcium channel links pathogen patterns to plant immunity. Nature 572, 131-+ (2019).

14. Y. Kadota et al., Direct regulation of the NADPH oxidase RBOHD by the PRR-associated kinase BIK1 during plant immunity. Mol Cell 54, 43–55 (2014).

15. L. Li et al., The FLS2-Associated Kinase BIK1 Directly Phosphorylates the NADPH Oxidase RbohD to Control Plant Immunity. Cell Host & Microbe 15, 329–338 (2014).

16. A. P. Macho, C. Zipfel, Plant PRRs and the activation of innate immune signaling. Mol Cell 54, 263–272 (2014).

17. K. Sato, Y. Kadota, K. Shirasu, Plant Immune Responses to Parasitic Nematodes. Front Plant Sci 10, 1165 (2019).

18. P. Manosalva et al., Conserved nematode signalling molecules elicit plant defenses and pathogen resistance. Nat Commun 6, 7795 (2015).

19. B. Mendy et al., Arabidopsis leucine-rich repeat receptor-like kinase NILR1 is required for induction of innate immunity to parasitic nematodes. PLoS Pathog 13, e1006284 (2017).

20. L. Huang et al., NILR1 perceives a nematode ascaroside triggering immune signaling and resistance. Curr Biol 33, 3992–3997 e3993 (2023).

21. L. Huang et al., A receptor for dual ligands governs plant immunity and hormone response and is targeted by a nematode effector. Proc Natl Acad Sci U S A 121, e2412016121 (2024).

22. Y. A. Millet et al., Innate immune responses activated in Arabidopsis roots by microbe-associated molecular patterns. Plant Cell 22, 973–990 (2010).

23. M. A. Teixeira, L. Wei, I. Kaloshian, Root-knot nematodes induce pattern-triggered immunity in Arabidopsis thaliana roots. New Phytol 211, 276–287 (2016).

24. K. Sato et al., Transcriptomic Analysis of Resistant and Susceptible Responses in a New Model Root-Knot Nematode Infection System Using Solanum torvum and Meloidogyne arenaria. Front Plant Sci 12, 680151 (2021).

25. M. Safaeizadeh, T. Boller, C. Becker, Comparative RNA-seq analysis of Arabidopsis thaliana response to AtPep1 and flg22, reveals the identification of PP2-B13 and ACLP1 as new members in pattern-triggered immunity. PLoS One 19, e0297124 (2024).

26. J. Wan et al., A LysM receptor-like kinase plays a critical role in chitin signaling and fungal resistance in Arabidopsis. Plant Cell 20, 471–481 (2008).

27. M. C. Guillou et al., The peptide SCOOP12 acts on reactive oxygen species homeostasis to modulate cell division and elongation in Arabidopsis primary root. J Exp Bot 73, 6115–6132 (2022).

28. S. Hou et al., The Arabidopsis MIK2 receptor elicits immunity by sensing a conserved signature from phytocytokines and microbes. Nat Commun 12, 5494 (2021).

29. J. Rhodes et al., Perception of a divergent family of phytocytokines by the Arabidopsis receptor kinase MIK2. Nat Commun 12, 705 (2021).

30. L. Gomez-Gomez, Z. Bauer, T. Boller, Both the extracellular leucine-rich repeat domain and the kinase activity of FSL2 are required for flagellin binding and signaling in Arabidopsis. Plant Cell 13, 1155–1163 (2001).

31. M. Kanehisa, Y. Sato, M. Kawashima, KEGG mapping tools for uncovering hidden features in biological data. Protein Sci 31, 47–53 (2022).

32. J. Rajniak, B. Barco, N. K. Clay, E. S. Sattely, A new cyanogenic metabolite in Arabidopsis required for inducible pathogen defence. Nature 525, 376–379 (2015).

33. B. Schwessinger et al., Phosphorylation-dependent differential regulation of plant growth, cell death, and innate immunity by the regulatory receptor-like kinase BAK1. PLoS Genet 7, e1002046 (2011).

34. M. Roux et al., The Arabidopsis leucine-rich repeat receptor-like kinases BAK1/SERK3 and BKK1/SERK4 are required for innate immunity to hemibiotrophic and biotrophic pathogens. Plant Cell 23, 2440–2455 (2011).

35. M. Gao et al., Regulation of cell death and innate immunity by two receptor-like kinases in Arabidopsis. Cell Host Microbe 6, 34–44 (2009).

36. M. Simon et al., Quantitative trait loci mapping in five new large recombinant inbred line populations of Arabidopsis thaliana genotyped with consensus single-nucleotide polymorphism markers. Genetics 178, 2253–2264 (2008).

37. Y. Wang, K. Bouwmeester, P. Beseh, W. Shan, F. Govers, Phenotypic analyses of Arabidopsis T-DNA insertion lines and expression profiling reveal that multiple L-type lectin receptor kinases are involved in plant immunity. Mol Plant Microbe Interact 27, 1390–1402 (2014).

38. M. Desclos-Theveniau et al., The Arabidopsis lectin receptor kinase LecRK-V.5 represses stomatal immunity induced by Pseudomonas syringae pv. tomato DC3000. PLoS Pathog 8, e1002513 (2012).

39. M. Da Rocha et al., Genome Expression Dynamics Reveal the Parasitism Regulatory Landscape of the Root-Knot Nematode Meloidogyne incognita and a Promoter Motif Associated with Effector Genes. Genes (Basel) 12, (2021).

40. D. Kuczynska-Wisnik, K. Stojowska-Swedrzynska, E. Laskowska, Intracellular Protective Functions and Therapeutical Potential of Trehalose. Molecules 29, (2024).

41. A. M. MacIntyre et al., Trehalose Synthesis Contributes to Osmotic Stress Tolerance and Virulence of the Bacterial Wilt Pathogen Ralstonia solanacearum. Mol Plant Microbe Interact 33, 462–473 (2020).

42. S. Djonovic et al., Trehalose biosynthesis promotes Pseudomonas aeruginosa pathogenicity in plants. PLoS Pathog 9, e1003217 (2013).

43. A. J. Foster, J. M. Jenkinson, N. J. Talbot, Trehalose synthesis and metabolism are required at different stages of plant infection by Magnaporthe grisea. EMBO J 22, 225–235 (2003).

44. X. Zhu et al., Phytophthora sojae boosts host trehalose accumulation to acquire carbon and initiate infection. Nat Microbiol 8, 1561–1573 (2023).

45. M. Vanaporn, R. W. Titball, Trehalose and bacterial virulence. Virulence 11, 1192–1202 (2020).

46. F. Baldacci-Cresp et al., Maturation of nematode-induced galls in Medicago truncatula is related to water status and primary metabolism modifications. Plant Sci 232, 77–85 (2015).

47. J. Hofmann et al., Metabolic profiling reveals local and systemic responses of host plants to nematode parasitism. Plant J 62, 1058–1071 (2010).

48. M. D. Garcia, J. C. Arguelles, Trehalase inhibition by validamycin A may be a promising target to design new fungicides and insecticides. Pest Manag Sci 77, 3832–3835 (2021).

49. V. Mani et al., Chitin Biosynthesis Inhibition of Meloidogyne incognita by RNAi-Mediated Gene Silencing Increases Resistance to Transgenic Tobacco Plants. Int J Mol Sci 21, (2020).

50. M. Van Ende et al., The involvement of the Candida glabrata trehalase enzymes in stress resistance and gut colonization. Virulence 12, 329–345 (2021).

51. Y. Pedreno et al., Disruption of the Candida albicans ATC1 gene encoding a cell-linked acid trehalase decreases hypha formation and infectivity without affecting resistance to oxidative stress. Microbiology (Reading) 153, 1372–1381 (2007).

52. R. Sanchez-Fresneda, M. Martinez-Esparza, S. Maicas, J. C. Arguelles, E. Valentin, In Candida parapsilosis the ATC1 gene encodes for an acid trehalase involved in trehalose hydrolysis, stress resistance and virulence. PLoS One 9, e99113 (2014).

53. R. G. Lopes et al., The secreted acid trehalase encoded by the CgATH1 gene is involved in Candida glabrata virulence. Mem Inst Oswaldo Cruz 115, e200401 (2020).

54. R. Shinya et al., Secretome Analysis of the Pine Wood Nematode Bursaphelenchus xylophilus Reveals the Tangled Roots of Parasitism and Its Potential for Molecular Mimicry. PLoS One 8, e67377 (2013).

55. H. Silva et al., Comparative Analysis of Bursaphelenchus xylophilus Secretome Under Pinus pinaster and P. pinea Stimuli. Front Plant Sci 12, 668064 (2021).

56. S. Siddique et al., The genome and lifestage-specific transcriptomes of a plant-parasitic nematode and its host reveal susceptibility genes involved in trans-kingdom synthesis of vitamin B5. Nat Commun 13, 6190 (2022).

57. B. Molloy et al., The origin, deployment, and evolution of a plant-parasitic nematode effectorome. PLoS Pathog 20, e1012395 (2024).

58. C. Pellegrin et al., The SUbventral-Gland master Regulator (SUGR) of nematode virulence. *bioRxiv*, 2024.2001.2022.576598 (2024).

59. J. Choi et al., Identification of a plant receptor for extracellular ATP. Science 343, 290–294 (2014).

60. C. Wang et al., Extracellular pyridine nucleotides trigger plant systemic immunity through a lectin receptor kinase/BAK1 complex. Nat Commun 10, 4810 (2019).

61. Q. Li, M. Zhou, F. Harris, Z. Mou, A group of L-type lectin receptor kinases function redundantly in mediating extracellular NAD(P) signaling in Arabidopsis. Plant Physiol 195, 2524–2527 (2024).

62. C. Gouhier-Darimont, E. Stahl, G. Glauser, P. Reymond, The Arabidopsis Lectin Receptor Kinase LecRK-I.8 Is Involved in Insect Egg Perception. Front Plant Sci 10, 623 (2019).

63. R. Groux et al., Arabidopsis natural variation in insect egg-induced cell death reveals a role for LECTIN RECEPTOR KINASE-I.1. Plant Physiol 185, 240–255 (2021).

64. S. Lacombe et al., Interfamily transfer of a plant pattern-recognition receptor confers broad-spectrum bacterial resistance. Nat Biotechnol 28, 365–369 (2010).

65. B. T. Townsley, M. F. Covington, Y. Ichihashi, K. Zumstein, N. R. Sinha, BrAD-seq: Breath Adapter Directional sequencing: a streamlined, ultra-simple and fast library preparation protocol for strand specific mRNA library construction. Frontiers in Plant Science 6, 366 (2015).

66. B. Langmead, C. Trapnell, M. Pop, S. L. Salzberg, Ultrafast and memory-efficient alignment of short DNA sequences to the human genome. Genome Biol 10, R25 (2009).

67. T. Suzuki et al., The DROL1 subunit of U5 snRNP in the spliceosome is specifically required to splice AT-AC-type introns in Arabidopsis. Plant J 109, 633–648 (2022).

68. T. Suzuki et al., Development of the Mitsucal computer system to identify causal mutation with a high-throughput sequencer. Plant Reprod 31, 117–128 (2018).

