## Supplemental methods, Supplemental Figures for "A trehalase-derived MAMP triggers LecRK-V-mediated immune responses in Arabidopsis"

**This PDF file includes:**

Supplementary Methods,
Figs. S1 to S24
Legends for tables S1 to S13
References

**Other Supplementary Materials for this manuscript include the following:**

Tables S1 to S13

### Supplementary Methods

#### RNA-Seq and differential gene expression analyses

Ten-day-old *Arabidopsis* seedlings of Col-0 or *efr fls2 cerk1* mutants, grown in liquid half-strength MS medium with 1% sucrose, were treated with 100 µg/mL *C. elegans* extract for 0.5, 1, 3, 6, and 12 hours (Fig. 1, C and D, and Table S1); with 100 µg/mL *M. incognita* extract for 12 hours (Table S2); and with 50 µM $Tre_{Mi}31$ for 12 hours (Fig. 1, J and K, and Table S7), with three biological replicates per condition. Additionally, ten-day-old seedlings of Col-0, *lecrk-V.5-3* (Fig. 3D and Table S12) were treated with 30 µM $Tre_{Mi}31$ for 12 hours with four biological replicates per genotype. Transcript levels were analyzed by RNA-Seq, with library preparation performed using the BradSeq protocol (*1*). Single-end 86-bp reads were sequenced on an Illumina NextSeq 500 platform and mapped to the Arabidopsis cDNA reference based on TAIR10 using Bowtie v0.12.9 (*2*). Read counts were obtained per transcript model (*3*) and Reads Per Million mapped reads (PPM) were calculated. Differentially expressed genes (DEGs) with an FDR ≤ 0.01 were identified using edgeR. Sequencing reads have been deposited in the DNA Data Bank of Japan (DDBJ) under accession number PRJDB19783 (BioProject: SAMD00851270-SAMD00851330).

#### Venn diagram analysis of transcript levels upon treatment with $Tre_{Mi}31$, flg22, and Chitin

Venn diagram analyses were performed using BioVenn (*4*). Genes upregulated or downregulated after treatment with flg22 (*5*), or chitin (*6*) were used.

#### Gene ontology (GO) term enrichment analysis

GO term enrichment analyses were performed using PANTHER (https://geneontology.org/) from the Gene Ontology Consortium.

#### Heatmap of transcript levels in Col-0 and *lecrk-V.5-3* upon treatment with $Tre_{Mi}31$

Heatmap was made using Heatmapper (*7*) (http://www.heatmapper.ca/).

#### Histochemistry of lignin deposition

Lignin deposition was visualized by phloroglucinol-HCl staining as previously described (*8*). Eight-day-old *Arabidopsis* Col-0 seedlings were treated with 100 µg/mL *C. elegans* extract, 50 µM $Tre_{Mi}31$, or distilled water (mock). Seedlings were fixed in a solution of ethanol : acetic acid (9:1, v/v) for 2 hours after a brief vacuum, washed twice with 90% ethanol for 30 min, and incubated overnight in distilled water. Samples were then stained with 2% phloroglucinol (w/v) in 20% HCl. Microphotographs were taken, processed as described and combined manually.

#### *C. elegans* culturing

*C. elegans* was cultured at 25°C on 6 cm Petri dishes containing Nematode Growth Medium (NGM) agar with a lawn of *E. coli* strain OP50. After 4 days on NGM agar plates, *C. elegans* were transferred to S medium supplemented with Streptomycin-Penicillin solution (100 units/mL, FUJIFILM Wako Pure Chemical Corporation, Osaka, Japan) and freeze-dried *E. coli* OP50 powder (2 g/500 mL, LabTIE, MOLGEN Veenendaal, The Netherlands) and grown on a rotary shaker at 25°C. On the fifth day, an additional 2 g/500 mL of freeze-dried *E. coli* OP50 powder was added, and the culture was incubated for another 3 days. After the incubation period, the liquid culture was cooled on ice, and *C. elegans* was collected by gravity sedimentation. The worms were then thoroughly washed with water, filtered through filter paper by aspiration, and frozen using liquid nitrogen.

**Purification of MAMPs from *C. elegans* by chromatography**

*C. elegans* was dissolved in the Extraction Buffer (20 mM Tris, pH 8.0, 0.1% CHAPS (FUJIFILM Wako Pure Chemical Corporation), cOmplete Ultra Protease Inhibitor Cocktail (Merck, Rahway, New Jersey), and proteins were extracted by sonication. The crude extract was centrifuged at 12,000 g for 10 min and repeated more than three times to remove debris. Immunogenic proteins in the cleared crude extract were precipitated with 40–70% ammonium sulfate. The resulting precipitate was dissolved in Extraction Buffer, followed by debris and aggregate removal through centrifugation at 12,000 g for 10 min and membrane filtration using syringe filters (5 µm, 1.2 µm, 0.8 µm, and 0.45 µm; Minisart, Sartorius, Goettingen, Germany) before purification on an AKTA FPLC system (Cytiva, Marlborough, MA). The crude extract was first desalted using a HiPrep 26/10 desalting column (Cytiva) and Desalting Buffer (same as Extraction Buffer). The desalted extract was then applied to a HiTrap Q anion exchange chromatography column pre-equilibrated with Anion Exchange Binding Buffer (20 mM Tris, pH 8.0, 0.1% CHAPS). Bound proteins were eluted with increasing salt concentrations using Anion Exchange Elution Buffer (20 mM Tris, pH 8.0, 0.1% CHAPS, 1 M NaCl). Each fraction was desalted using an ultrafiltration column (10 kDa MWCO) with 10 mM MES, pH 5.6, and subsequently tested for immunogenic activity by GUS assay with the *cerk1 fls2 pCYP71A12:GUS* line. Fractions with immunogenic activity were then mixed with Hydrophobic Interaction Binding Buffer (20 mM HEPES, pH 7.0, 2 M ammonium sulfate) and applied to a HiTrap butyl hydrophobic interaction chromatography column (Cytiva). Proteins were eluted by gradually decreasing the ammonium sulfate concentration using Hydrophobic Interaction Elution Buffer (20 mM HEPES, pH 7.0), and the immunogenic activity of each fraction was analyzed by GUS assay. Fractions with immunogenic activity were concentrated using an ultrafiltration column (10 kDa MWCO) and subjected to Superdex 200 gel filtration chromatography in Gel Filtration Buffer (20 mM Tris, pH 8.0, 150 mM NaCl). The buffer of fractions with immunogenic activity was exchanged to Cation Exchange Binding Buffer (20 mM MES, pH 5.5, 0.1% CHAPS) using an ultrafiltration column (10 kDa MWCO) and then applied to a HiTrap SP cation exchange chromatography column. The final fractions with immunogenic activity were submitted for LC-MS/MS analysis to identify proteins.

**LC-MS/MS analyses**

Proteins were separated by SDS-PAGE (NuPAGe ®, Invitrogen) for 5-10 min, and then the gel areas containing proteins were cut out and digested using an improved in-gel digestion method (*9*) followed by nano-LC-MS/MS using a Q Exactive LC mass spectrometer coupled to an Ultimate 3000 nano-LC system (Thermo Fisher Scientific, Waltham, MA).

**RT-qPCR assay**

RT-qPCR was performed as described previously (*10*). Total RNA was extracted from Arabidopsis seedlings using an RNeasy Plant Mini Kit (Qiagen, Hilden, Germany) or Maxwell RSC Plant RNA Kit (Promega, WI, USA) according to the manufacturer's instructions. RNA was reverse transcribed with a ReverTraAce qPCR RT Kit (Toyobo, Osaka, Japan) according to the manufacturer's instructions. One µg of total RNA was used as a template for cDNA synthesis. RT-qPCR was carried out using Thunderbird SYBR qPCR Mix (Toyobo) with a Stratagene Mx3000p real-time thermal cycler (Agilent, CA, USA). Relative transcript levels were calculated against a standard curve with normalization to the *U-box* housekeeping gene transcript (*At5g15400*). Primers used for the RT-qPCR are listed in Table S13.

**KEGG pathway analysis**

KEGG pathway analysis was conducted using KEGG Mapper (https://www.genome.jp/kegg/mapper/).

**Vector construction for the complementation line and CRISPR/Cas9-mediated genomic deletion lines**

To generate the complementation line, *lecrk-V.5-2/pLecRK-V.5:LecRK-V.5-3×HA*, the genomic region of *LecRK-V.5* was amplified by PCR using KOD One (Toyobo). The resulting PCR products were cloned into epiGreenB5(3xHA) between the *EcoRI* and *BamHI* restriction sites using an In-Fusion HD Cloning Kit (Clontech, CA, USA). *LecRK-V.5-3×HA* in epiGreenB5 was again amplified by PCR using KOD One and the resulting PCR products were cloned into pBin19g vector between the *EcoRI* and *BamHI* restriction sites using an In-Fusion HD Cloning Kit. CRISPR/Cas9-mediated genomic deletion lines (*lecrk-V.567-d*, *lecrk-V.5-3/lecrk-V.78-d*, *lecrk-V.8/lecrk-V.56-d*) were generated using the pKAMA-ITACHI vector (pKI1.1R) as previously described(*11*). Optimal sgRNA sequences were identified using CRISPR-P (http://crispr.hzau.edu.cn/CRISPR2/), CRISPR-PLANT v2 (http://omap.org/crispr2/), CRISPOR (http://crispor.gi.ucsc.edu/), and Chopchop (https://chopchop.cbu.uib.no/). To insert an additional cassette containing the *AtU6.26* promoter, sgRNA, and polyT sequence into pKI1.1R, the cassette from pCRIPSR was amplified by PCR with FW and RV primers targeting two different sequences. The amplified fragments were inserted into the *AarI*-cut pKI1.1R vector using the In-Fusion HD Cloning Kit. Constructs were introduced into *Arabidopsis* via Agrobacterium-mediated transformation using floral dip and floral drop methods. The complimentation lines of *lecrk-V.5-2/pLecRK-V.5:LecRK-V.5-3×HA (pBin19g)* were generated by direct transformation of *pLecRK-V.5:LecRK-V.5-3×HA (pBin19g)* to *lecrk-V.5-2* mutant. To select CRISPR/Cas9-mediated genomic deletion lines, T1 seeds were screened for OLE1-RFP fluorescence, and genomic deletions were confirmed by PCR. In the T2 generation, seeds lacking the OLE1-RFP signal were selected to exclude transgene-harboring plants. Primers used for sgRNA construction and genotyping are listed in Table S13.

**Propagation of *M. incognita* and infection assay**

*M. incognita* was propagated on *Solanum lycopersicum* cultivar "Micro-Tom" in a greenhouse. Nematode eggs were isolated from infected roots and hatched at 25°C. Freshly hatched J2 juveniles were collected and transferred to a Kimwipe filter placed over a glass beaker filled with sterilized distilled water containing 100 µg/mL Streptomycin-Penicillin and 10 µg/mL nystatin overnight. Active J2s passed through the filter were collected and surface-sterilized with 0.002% mercuric chloride, 0.002% sodium azide, and 0.001% Triton X-100 for 10 minutes, followed by three rinses with distilled water (*12*). Sterilized J2s were further filtered by Kimwipes to obtain only active J2s. Arabidopsis seeds were sown and grown on half-strength MS medium with 1% sucrose and 0.8% agar under long-day conditions at 23°C. Four or five-day-old seedlings were transferred to quarter-strength MS medium with 0.5% sucrose and 0.6% phytagel pH 5.7. Twelve to thirteen-day-old seedlings were inoculated with approximately 80 nematodes per plant, and the roots were covered with black paper to simulate below-ground conditions. Gall numbers were counted six weeks after the inoculation under the microscope.

**ROS burst assay**

Eight ten-day-old seedlings grown in 96 well plates and a solution containing 1 µM L-012 (FUJIFILM Wako, Tokyo, Japan) and 20 µg/mL horseradish peroxidase (HRP) (Sigma-Aldrich, St. Louis, MO, USA) were used. Luminescence was measured using a Tristar2 multimode reader (Berthold Technologies, Bad Wildbad, Germany).

**MAPK activation assay**

MAPK activation assays were performed as described previously (*13*). Ten-day-old *Arabidopsis* seedlings were flash-frozen in liquid nitrogen, and proteins were extracted using protein extraction buffer (50 mM Tris-HCl, pH 7.5, 150 mM NaCl, 10% glycerol, 2 mM EDTA, 5 mM DTT, 1x EDTA-free cOmplete Protease Inhibitor Cocktail (Merck), 0.1% IGEPAL CA630 (Merck), 0.5 mM PMSF, 1 mM $Na_2MoO_4$, 1 mM NaF, 0.5 mM $Na_3VO_4$, 20 mM β-glycerophosphate). The extract was centrifuged at 16,000 × g to remove insoluble material and protein concentration was determined by the Bradford method (Bio-Rad Laboratories, Hercules, CA). Proteins were separated by SDS-PAGE and transferred onto a PVDF membrane (Transblot, Bio-Rad Laboratories) following the manufacturer's instructions. Membranes were blocked overnight at 4°C in 5% (w/v) skim milk (FUJIFILM Wako Pure Chemical Corporation) in TBS-T. Phosphorylated MAPKs were detected using α-phospho-p44/42 MAPK (Erk1/2) (Thr202/Tyr204) rabbit monoclonal antibody (1:2000, Cell Signaling Technology, Danvers, MA) for 1 hour at room temperature in 5% (w/v) BSA (Sigma-Aldrich) in TBS-T, followed by a 1-h incubation with α-rabbit IgG-HRP secondary antibody (1:10000, Roche, Basel, Germany) in 5% skim milk in TBS-T. Detection was performed using SuperSignal West Femto Maximum Sensitivity Substrate (Thermo Fisher Scientific) with a LAS 4000 system (GE Healthcare, Chicago, IL, USA). PVDF membranes were stained with Coomassie Brilliant Blue (CBB) to confirm equal loading.

**Phylogenetic analyses**

The phylogenetic trees in Fig. S21 were drawn using the previously published data and visualized and pruned, and figures were generated with iTOL (*14, 15*). For alignment and phylogeny methods, refer to https://github.com/MWSchmid/Ngou-et-al.-2022. Alignment files and tree files were taken from https://doi.org/10.5281/zenodo.7017981.

**Co-expression gene network analysis**

Genes that co-express with *LecRK-V.5* in *Arabidopsis* were analyzed using ATTED-II (https://atted.jp) (*16*).

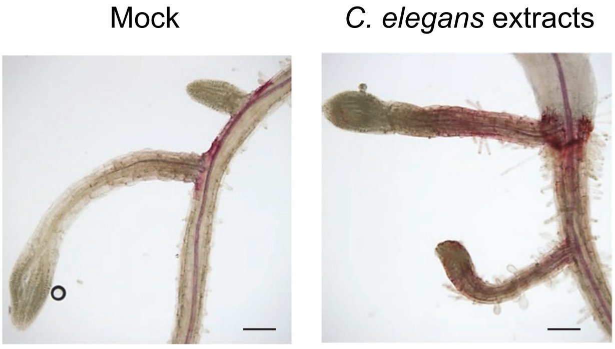


**Fig. S1. *C. elegans* extracts induce ectopic lignin accumulation in Col-0 roots.**
Col-0 seedlings were treated with 100 µg/mL *C. elegans* extracts for 24 h, followed by lignin staining using phloroglucinol. Experiments were repeated three times with consistent results. Black bars represent 100 µm.

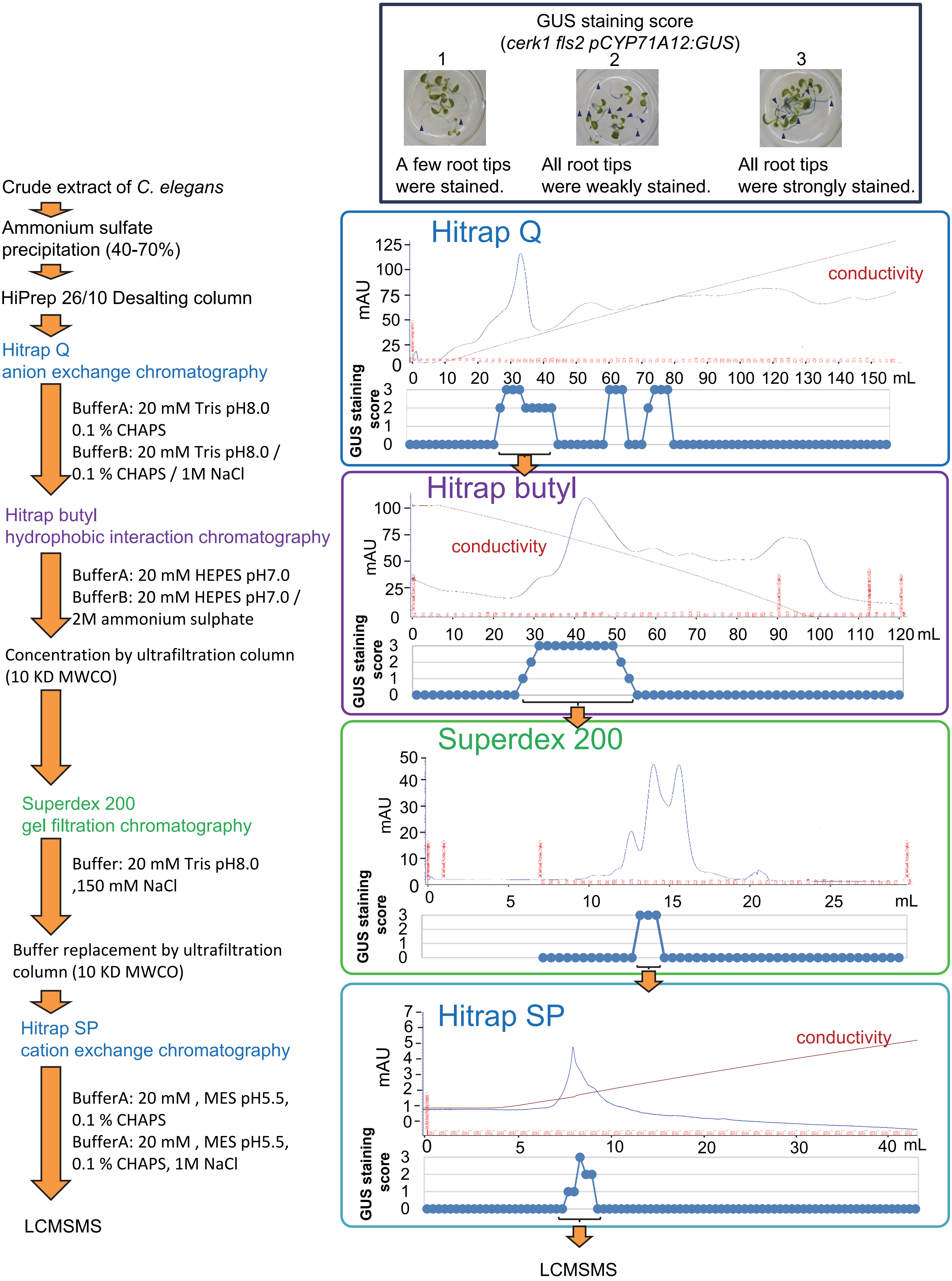


**Fig. S2. Workflow for MAMP identification from *C. elegans* extracts using chromatography and LC-MS/MS.** MAMP activity in each fraction was assessed using the *CYP71A12* expression assay. Scores were assigned based on the number of stained root tips and staining intensities. The detailed protocol is provided in the Supplementary Methods.

**A**

| | Candidate MAMPs | Synthesized peptides |
|---|---|---|
| 1 | | RYSLLYVPNSFIVPGGRFREFYYWDAYWIIKGLIASDM |
| 2 | | VDKYGFVPNGGRVYYLQRSQPPFL |
| 3 | Trehalase (F5GUD3, G5EG27) | DLASAAESGWDFSTRW |
| 4 | | GVFTYPGGIPTSMSQESDQQWDFPNGWSPNNHMIIEGLRKSAN |
| 5 | | GGEYDVQDGFGWSNGAILDLLLTYNDRL |
| 6 | Aspartic protease (G5EEI4) | ITLGTPPQPATVVLDTGSSNLWVID |
| 7 | | SWILGDTFIRQYCNVYDIGNGQIGFATAE |
| 8 | | IRDQSDCGSCWAFAAAEAISDRTCIASNG |
| 9 | Cathepsin B (P43509) | DNGTPYWLVANSWNVAWGEKGYFRIIRGLNECGIEH |
| 10 | | LVANSWNVAWGEKGYFRIIRGLN |
| 11 | Cathepsin B (P43508) | PYWLVANSWNVNWGENGYFRIIRGTN |
| 12 | | PTDPFWRQVNLTFAQLTGIYDAY |
| 13 | | PGHIVTFSGYPGVLISTDDYTITSAGLTSIETTIAIFNQTLY |
| 14 | PLB-like 3-1 (Q9BL07) | FGRYNSGTYNNQWTVLDWKQFTPEKELPD |
| 15 | | YLKKYTYFASYNIPFLPKVSEISGF |
| 16 | | PRARIFDRDHSKVTDIDSLTKLMRYNDYTHEEFARCKCTP |
| 17 | Neprilysin-1 (Q18673) | AFPAGILQQPFFDARFPKALNYGGIGAVIGHEITHGFDD |
| 18 | GILT-like protein (O17861) | CQHGEEECSINKFE |
| 19 | Uncharacterized protein(Q9XTU5) | PDKIKLVTLGQPRTGDYAFA |

**B**

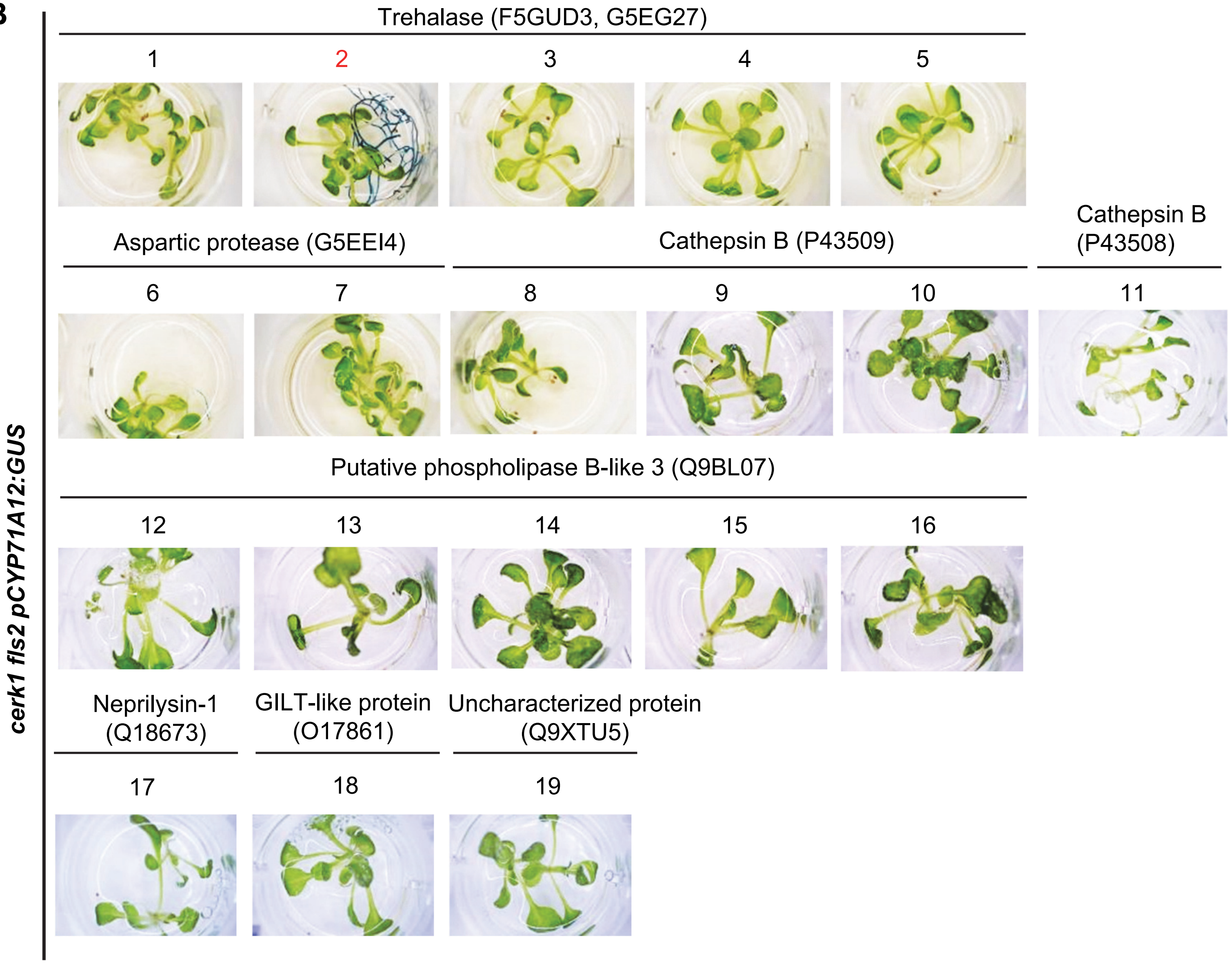


**Fig. S3. Candidate MAMPs peptides.**
(**A**) A list of candidate MAMPs peptides synthesized using highly conserved regions between *C. elegans* and plant-parasitic nematodes (PPNs). (**B**) *CYP71A12* expression in roots in *cerk1 fls2 pCYP71A12:GUS* line upon treatment with 60 µM peptides from candidate MAMPs. Experiments were repeated three times with consistent results. MAMP-active peptide was shown in red.

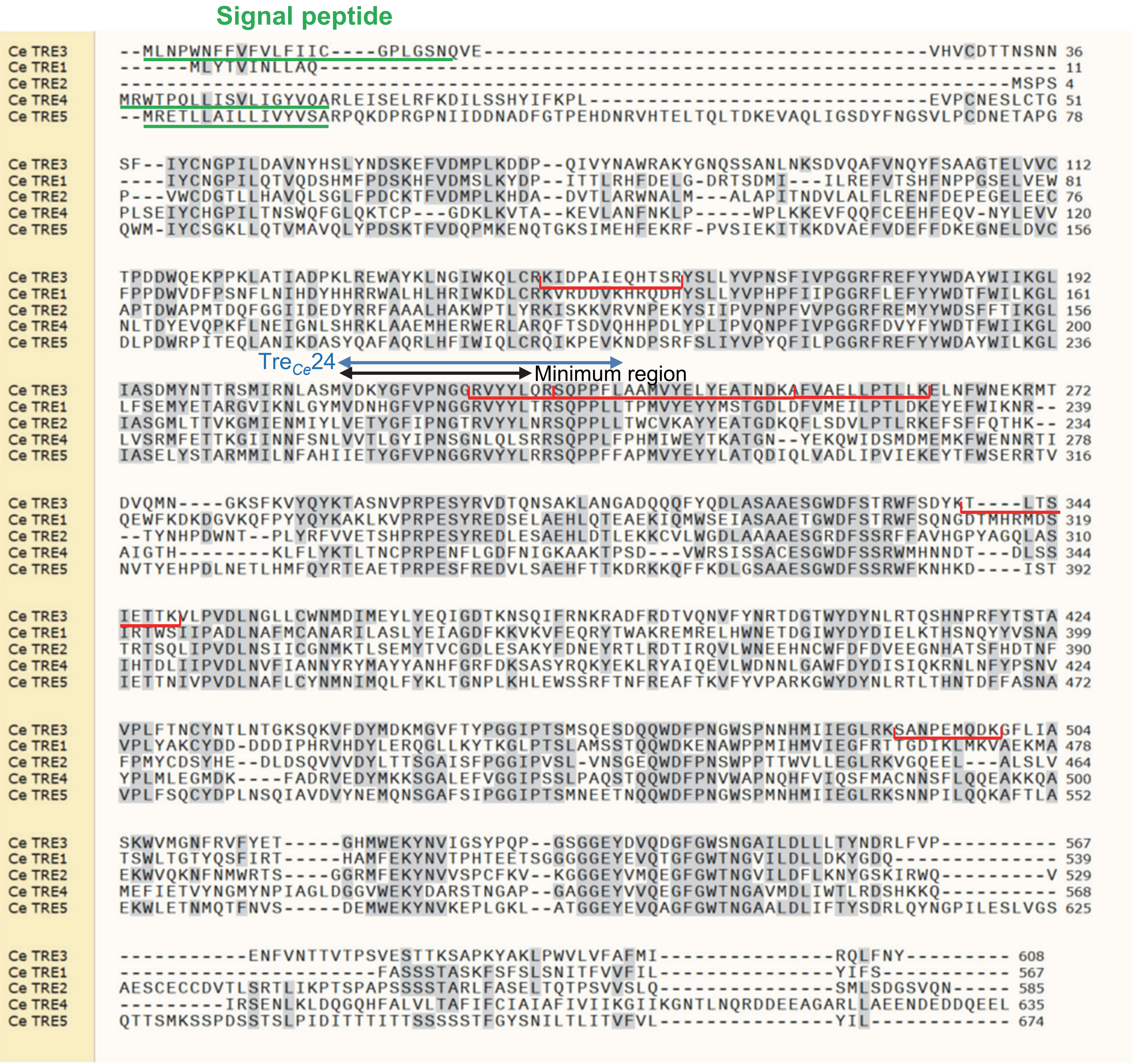


**Fig. S4. Amino acid sequence alignment of trehalase proteins in *C. elegans*.**
Specific peptides for *Ce*TRE3 identified by LC-MS/MS are indicated with red lines. Regions corresponding to predicted signal peptides by SignalP-6.0 (https://services.healthtech.dtu.dk/services/SignalP-6.0/), $\mathrm{Tre}_{Ce}24$, and the minimal region required for MAMP activity are also highlighted.

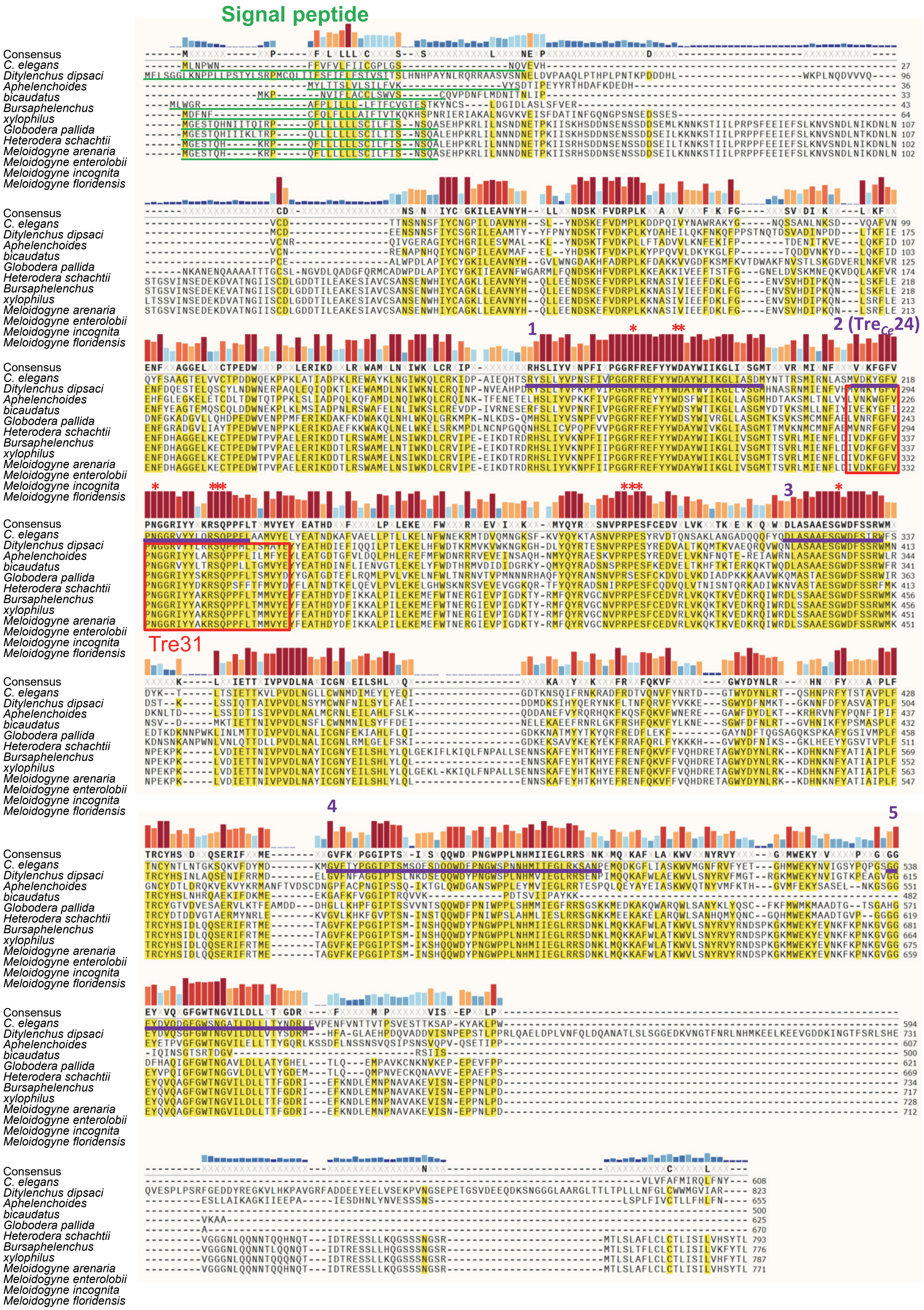


**Fig. S5. Amino acid sequence alignment of secreted trehalase proteins of *C. elegans* and PPNs.** Five highly conserved regions between *C. elegans* and PPNs, tested for MAMP activity, are highlighted with purple lines (see fig. S3). Regions corresponding to predicted signal peptides by SignalP-6.0 are highlighted with green lines. The 31 amino acid residues of PPNs (Tre31) shown in Fig. 1F are enclosed in a red box. The degree of conservation of each residue among sequences is displayed as colored bars above the alignment.

**A**

*cerk1 fls2 pCYP71A12:GUS*

| Name | Sequence | 10 μM | 30 μM | 50 μM |
|---|---|---|---|---|
| Tre$_{Ce}$32 | VDKYGFVPNGGRVYYLQRSQPPFLAAMVYELY | | | |
| Tre$_{Ce}$26 | VDKYGFVPNGGRVYYLQRSQPPFLAA | | | |
| Tre$_{Ce}$24 | VDKYGFVPNGGRVYYLQRSQPPFL | | | |
| Tre$_{Ce}$23 | VDKYGFVPNGGRVYYLQRSQPPF | | | |
| Tre$_{Ce}$21 | VDKYGFVPNGGRVYYLQRSQP | | | |
| Tre$_{Ce}$20 | VDKYGFVPNGGRVYYLQRSQ | | | |
| Tre$_{Ce}$19 | VDKYGFVPNGGRVYYLQRS | | | |
| Tre$_{Ce}$18 | VDKYGFVPNGGRVYYLQR | | | |
| Tre$_{Ce}$17 | VDKYGFVPNGGRVYYLQ | | | |
| Tre$_{Ce}$16 | VDKYGFVPNGGRVYYL | | | |
| Tre$_{Ce}$15 | VDKYGFVPNGGRVYY | | | |

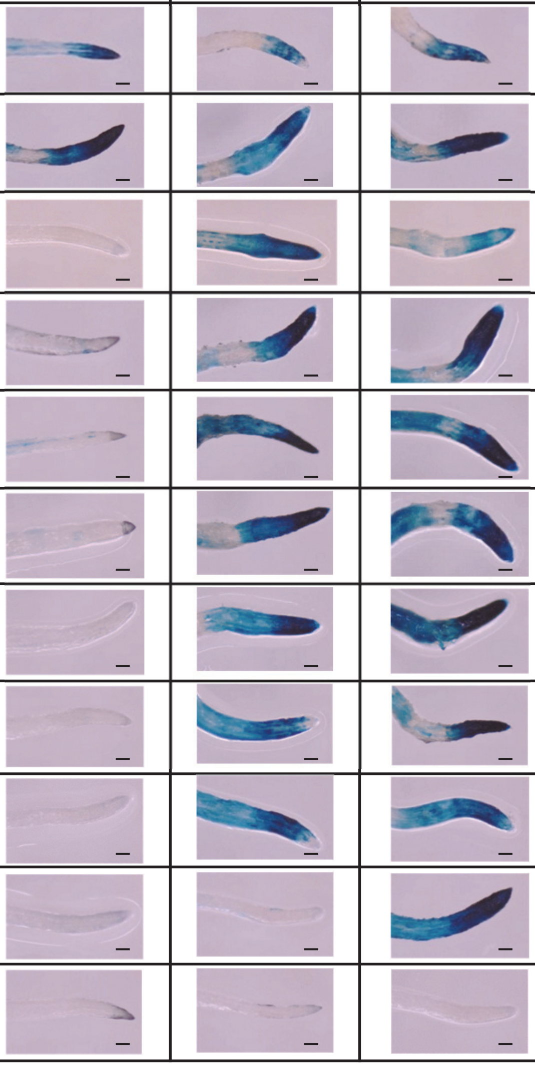

**B**

*cerk1 fls2 pCYP71A12:GUS*

| Name | Sequence | 10 μM | 30 μM | 50 μM |
|---|---|---|---|---|
| Tre$_{Ce}$24-2 | KYGFVPNGGRVYYLQRSQPPFL | | | |
| Tre$_{Ce}$24-4 | GFVPNGGRVYYLQRSQPPFL | | | |
| Tre$_{Ce}$24-6 | VPNGGRVYYLQRSQPPFL | | | |
| Tre$_{Ce}$24-8 | NGGRVYYLQRSQPPFL | | | |

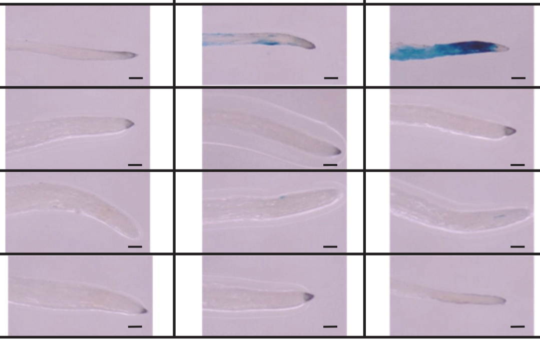

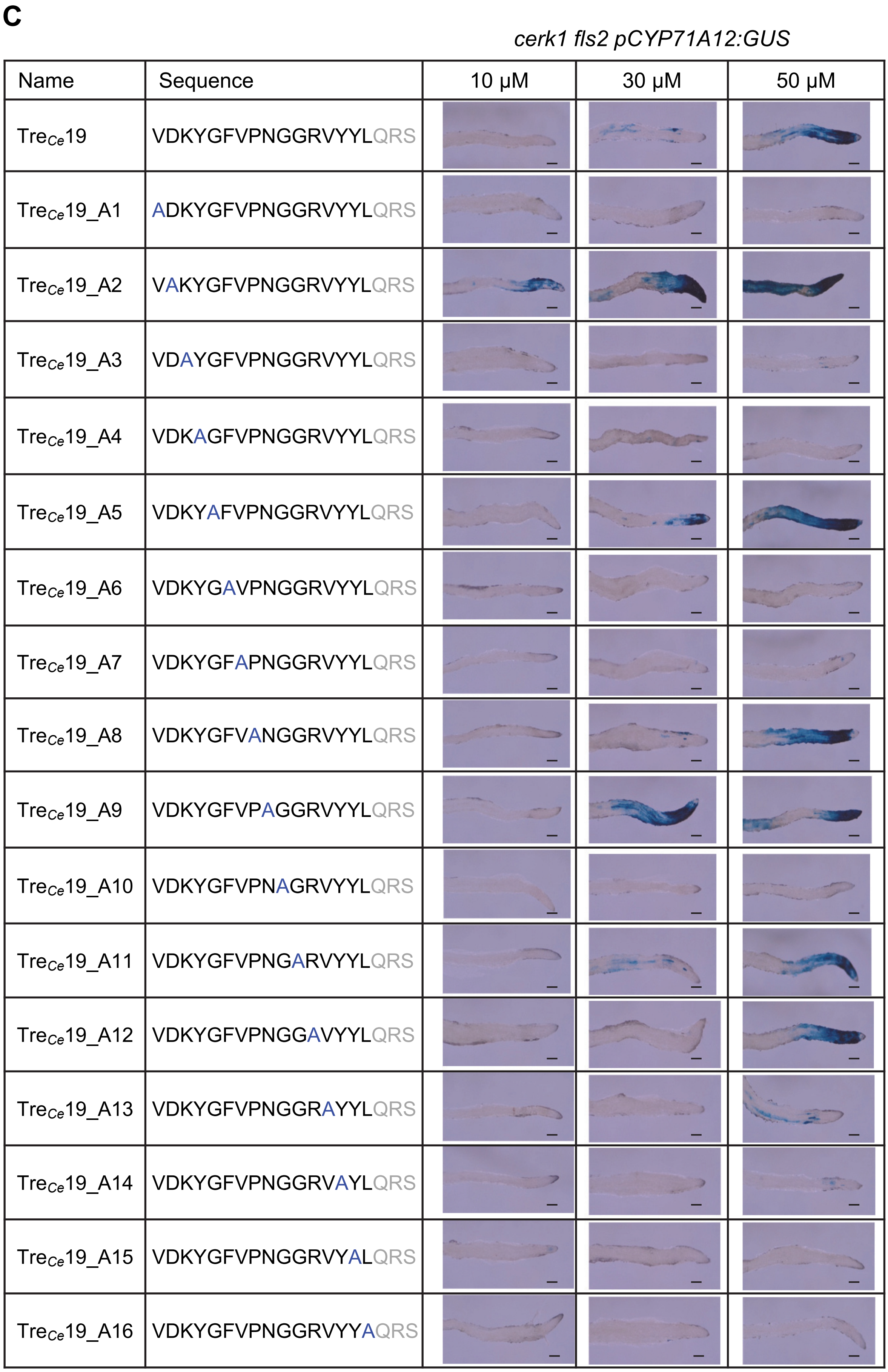


**Fig. S6. Minimal region and essential residues for MAMP activity of the *C. elegans* trehalase peptide.** (**A**) MAMP activity of C-terminally truncated trehalase peptides and Tre$_{Ce}$24 with hydroxylated prolines, determined via the *CYP71A12* expression assay using *cerk1 fls2 pCYP71A12:GUS* line. (**B**) MAMP activity of N-terminally truncated trehalase peptides. (**C**) Alanine scanning of the Tre$_{Ce}$19 peptide. Black bars represent 100 µm. Experiments were repeated three times with consistent results.

**A**

*cerk1 fls2 pCYP71A12:GUS*

| Name | | 50 µM | 100 µM |
|---|---|---|---|
| Tre *Ditylenchus dipsaci* | MVNKYGFVPNGGRVYYLRRSQP | 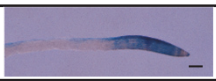 | |
| Tre *Aphelenchoides bicaudatus* | LVNKWGFVPNGGRIYYLARSQP | 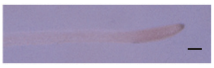 | 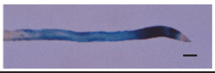 |
| Tre *Bursaphelenchus xylophilus* | IVEKYGFIPNGGRVYYLTRSQP | 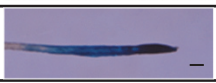 | |
| Tre *Globodera pallida* | LVNRFGFVPNGGRIYYSKRSQP | 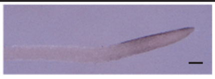 | 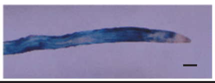 |
| Tre *Heterodera schachtii* | MVNRFGFVPNGGRIYYDKRSQP | 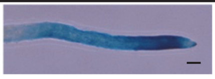 | |

**B**

*cerk1 fls2 pCYP71A12:GUS*

| Name | Sequence | 10 µM | 30 µM | 50 µM |
|---|---|---|---|---|
| Tre$_{Mi}$31 | IVDKFGFVPNGGRIYYAKRSQPPFLTMMVYE | 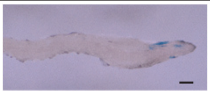 | 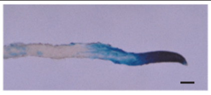 | 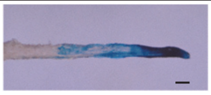 |
| Tre$_{Mi}$16 | VDKFGFVPNGGRIYYA | 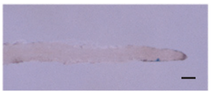 | 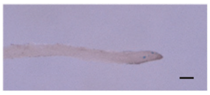 | 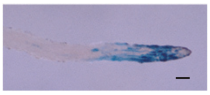 |

**Fig. S7. MAMP activity of trehalase peptides from plant parasitic nematodes.**
(**A**) MAMP activity of trehalase peptides from plant parasitic nematodes, assessed via the *CYP71A12* expression assay using *cerk1 fls2 pCYP71A12:GUS* line. (**B**) MAMP activity of Tre$_{Mi}$16 and Tre$_{Mi}$31 peptide. Black bars represent 100 µm. Experiments were repeated three times with consistent results.

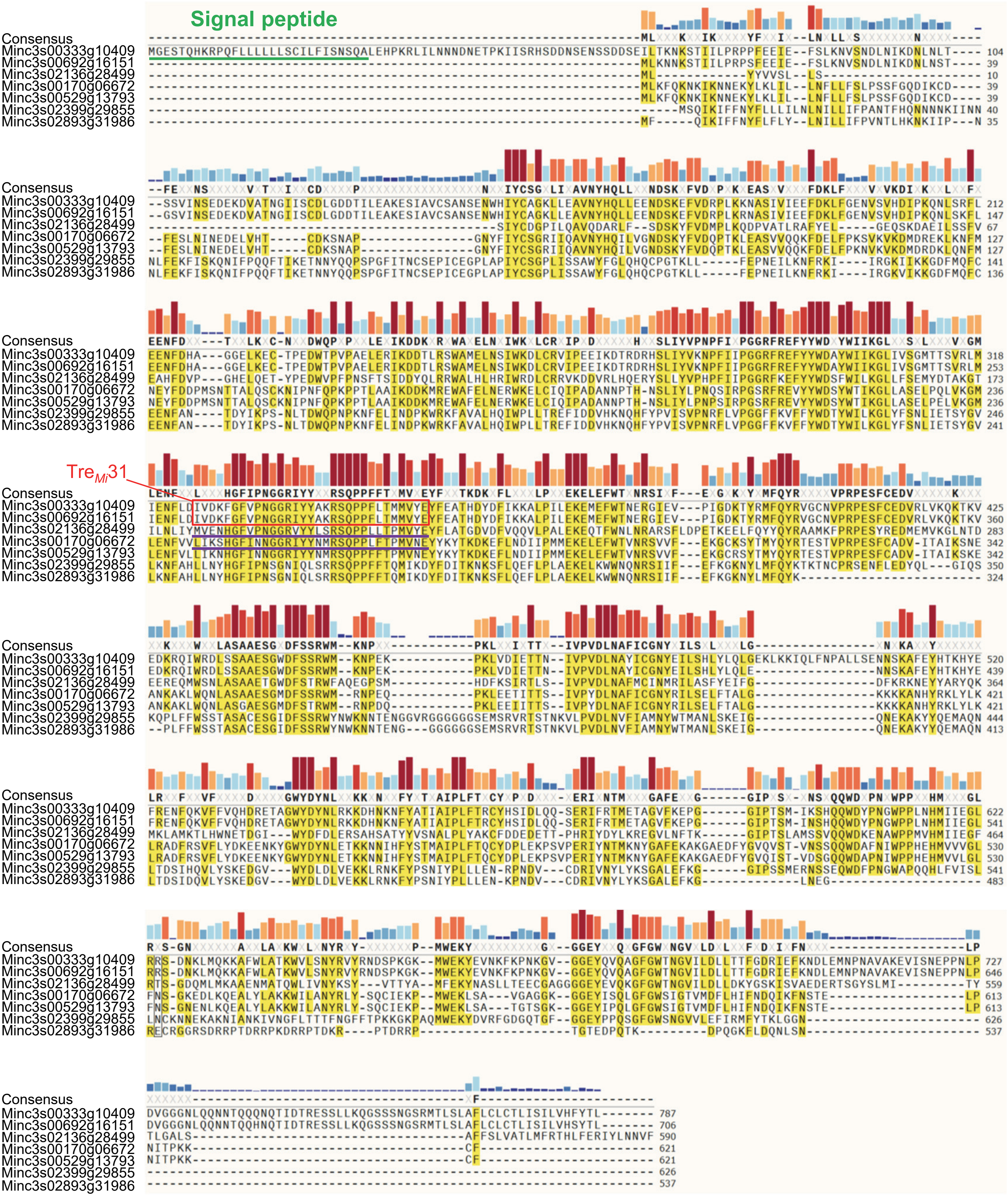


**Fig. S8. Amino acid sequence alignment of trehalase proteins in *M. incognita*.**

Amino acid sequence alignment of trehalase homologs longer than 500 amino acid residues. Regions corresponding to the predicted signal peptide by SignalP-6.0 and Tre$_{Mi}$31 are highlighted with a green line and a red box, respectively. The degree of conservation for each residue among sequences is displayed as colored bars above the alignment.

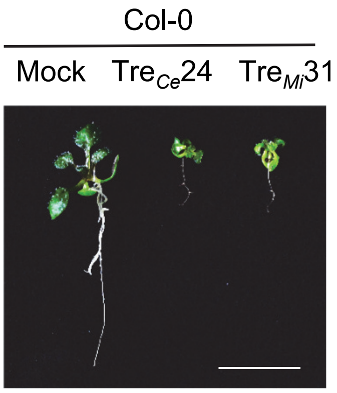


**Fig. S9. Characterization of trehalase peptide-induced responses.**
50 μM $Tre_{Ce}24$ and $Tre_{Mi}31$ induce seedling growth inhibition in Col-0. The white bar represents 1 cm.

**A** **Phe, Tyr, and Trp biosynthesis**

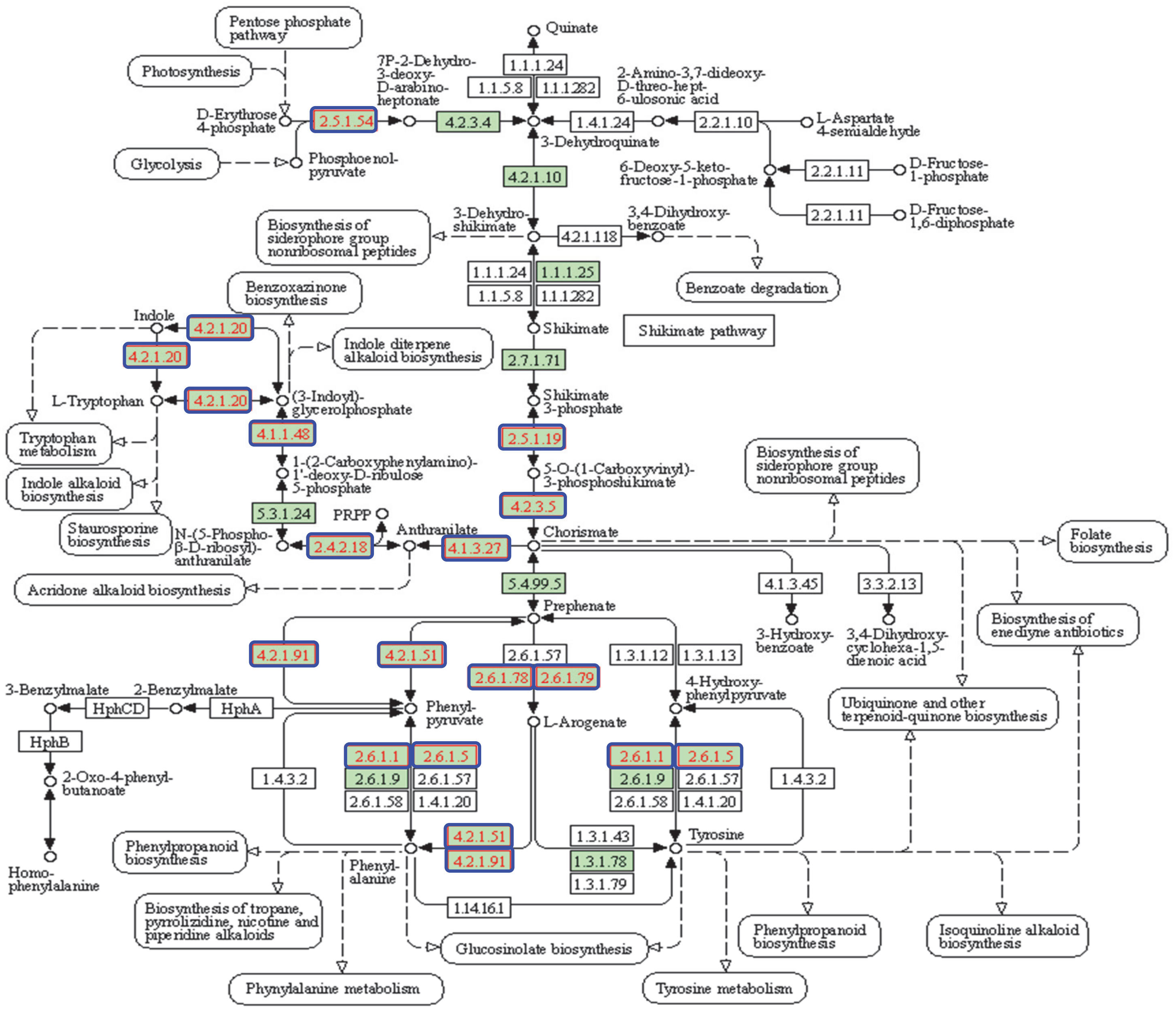


Expression upon Tre$_{Mi}$31 treatment for 12 hours

| | Gene ID | logFC | adj_ pvalue |
|---|---|---|---|
| Class-II DAHP synthetase | AT1G22410 | 2.07 | 2.64E-20 |
| DHS1 | AT4G39980 | 1.87 | 1.42E-40 |
| TSA1 | AT3G54640 | 3.41 | 1.79E-145 |
| IGPS | AT2G04400 | 2.98 | 7.49E-88 |
| EPSP synthase | AT2G45300 | 1.61 | 1.31E-18 |
| chorismate synthase | AT1G48850 | 1.56 | 5.56E-25 |
| Glutamine amidotransferase | AT1G24807 | 3.51 | 2.44E-38 |
| ADT4 | AT3G44720 | 1.98 | 2.61E-18 |
| AAT | AT2G22250 | 1.64 | 3.95E-18 |
| Tyrosine transaminase | AT4G28420 | 8.85 | 2.36E-19 |

**B** **Phenylpropanoid biosynthesis**

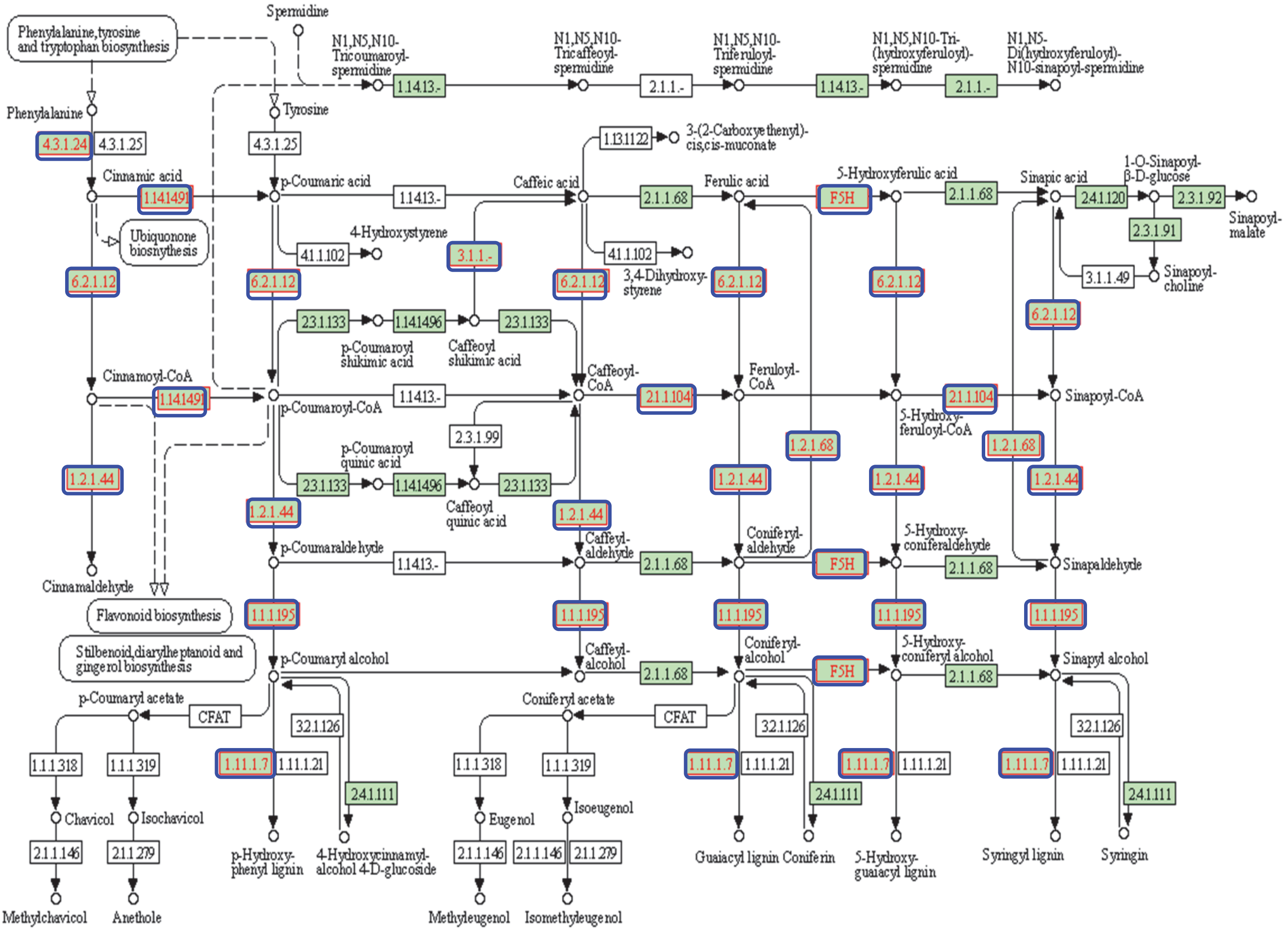


Expression upon Tre$_{Mi}$31 treatment for 12 hours

| | Gene ID | logFC | adj_pvalue |
|---|---|---|---|
| PAL | AT2G37040 | 1.84 | 5.32E-15 |
| C4H | AT2G30490 | 1.73 | 5.16E-20 |
| F5H,FAH1 | AT4G36220 | 1.10 | 3.31E-07 |
| 4CL | AT1G51680 | 1.83 | 6.56E-20 |
| LysoPL2 | AT1G52760 | 1.09 | 1.35E-11 |
| cinnamate-4-hydroxylase | AT2G30490 | 1.73 | 5.16E-20 |
| CCOAMT | AT1G67980 | 8.94 | 4.28E-169 |
| CCR2 | AT1G80820 | 3.10 | 4.05E-22 |
| ALDH2C4 | AT3G24503 | 1.75 | 2.32E-29 |
| CAD | AT4G37990 | 2.94 | 8.01E-06 |
| FAH1 | AT4G36220 | 1.10 | 3.31E-07 |
| Peroxidase | AT1G14540 | 7.98 | 1.27E-65 |

**C**

Col-0

Mock Tre$_{Mi}$31

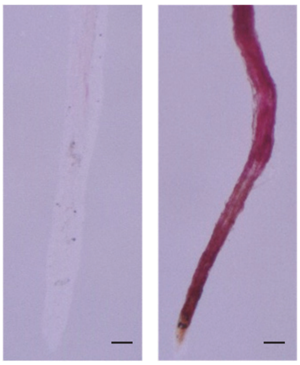

**D**

### Indole glucosinolate biosynthesis

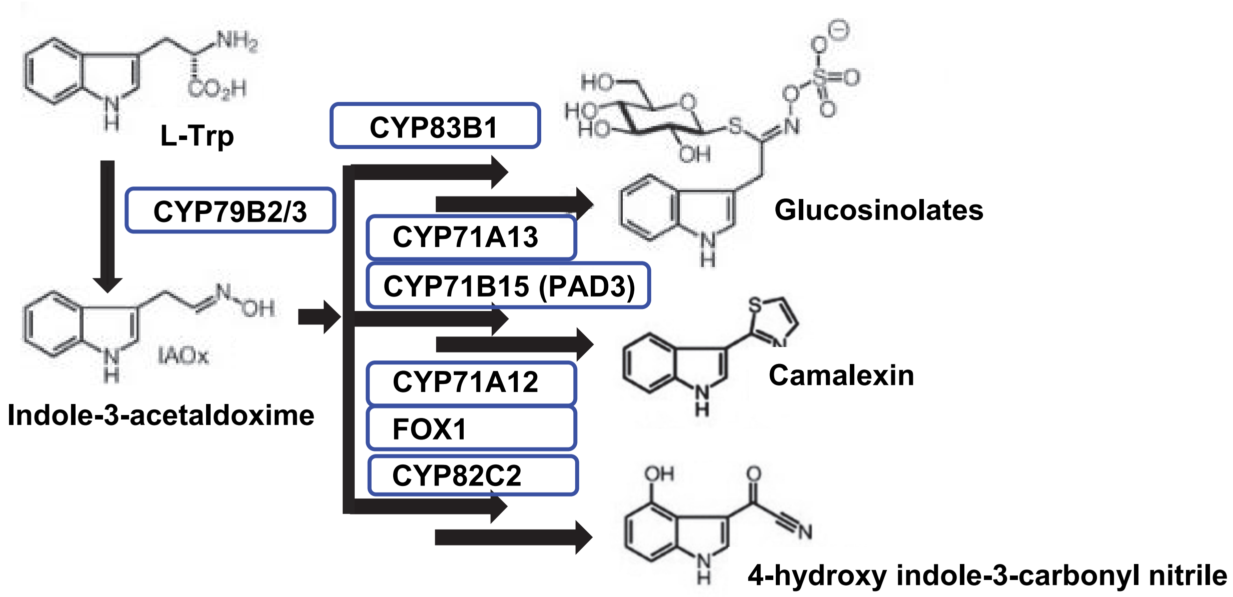


Expression upon Tre$_{Mi}$31 treatment for 12 hours

| | Gene ID | logFC | adj_pvalue |
|---|---|---|---|
| CYP79B2 | AT4G39950 | 3.38 | 6.19E-42 |
| CYP83B1 | AT4G31500 | 2.10 | 2.06E-34 |
| CYP71A13 | AT2G30770 | 6.44 | 2.57E-82 |
| CYP71B15 (PAD3) | AT3G26830 | 6.82 | 7.67E-121 |
| CYP71A12 | AT2G30750 | 9.13 | 2.99E-196 |
| FOX1 | AT1G26380 | 9.09 | 9.26E-165 |
| CYP82C2 | AT4G31970 | 10.71 | 3.91E-52 |

**Fig. S10. Pathway analysis of genes specifically upregulated upon treatment with Tre$_{Mi}$31.**
(**A** and **B**) KEGG (Kyoto Encyclopedia of Genes and Genomes) pathway analysis of genes involved in phenylalanine, tyrosine, and tryptophan biosynthesis (**A**) and phenylpropanoid biosynthesis (**B**), whose expressions are specifically induced by the treatment with 30 µM Tre$_{Mi}$31 for 12 hours (Table S7). Enzymes with expression induced by Tre$_{Mi}$31 are shown in blue boxes. (**C**) Tre$_{Mi}$31 induces lignin accumulation in Col-0 seedlings. Lignin was stained using phloroglucinol upon treatment with 50 µM Tre$_{Mi}$31 for 3 days. Black bars represent 100 µm. Experiments were repeated three times with consistent results. (**D**) Pathway analysis of genes involved in indole glucosinolate biosynthesis whose expressions are specifically induced by the treatment with 30 µM Tre$_{Mi}$31 for 12 hours. Enzymes with expression induced by Tre$_{Mi}$31 are shown in blue boxes.

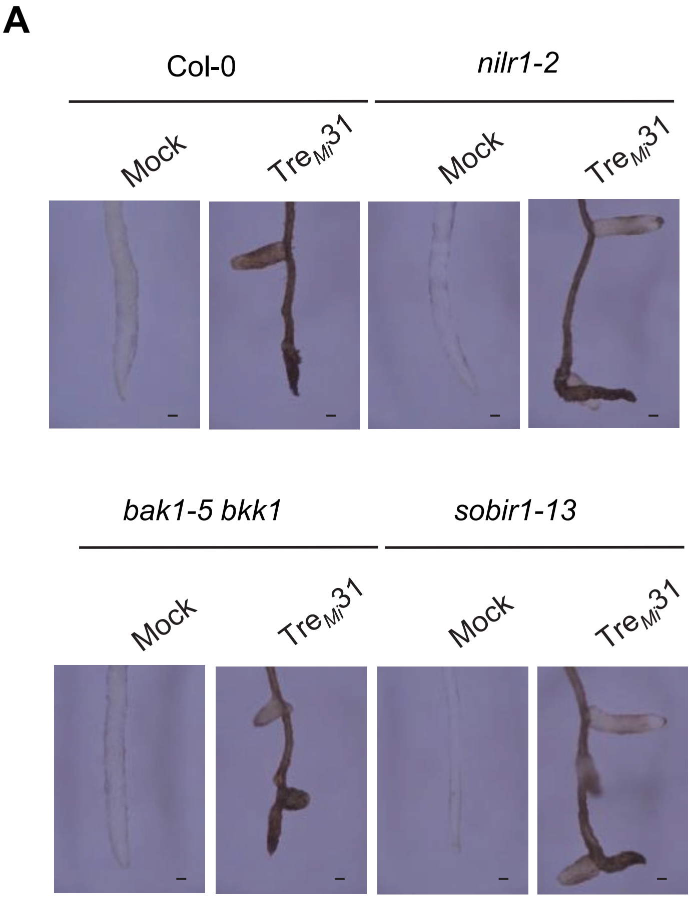


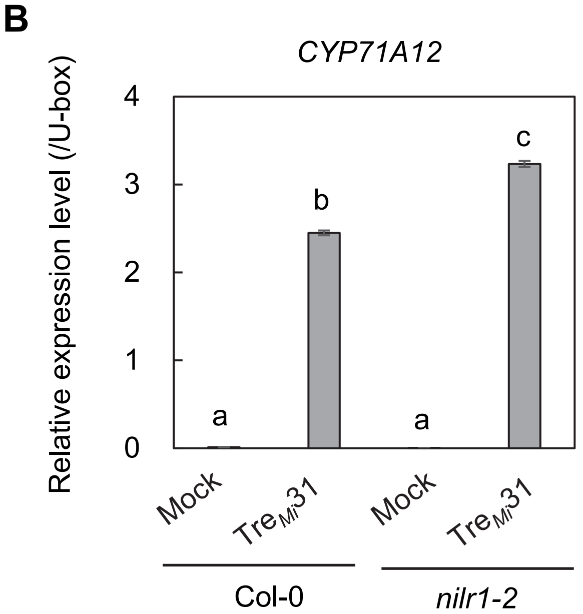


**Fig. S11. NILR1, BAK1, and SOBIR1 are not required for Tre$_{Mi}$31-induced responses.**
(**A**) *nilr1-2*, *bak1-5 bkk1*, and *sorbir1-13* mutants induce root pigmentation similar to Col-0 upon treatment with 50 µM Tre$_{Mi}$31. Black bars represent 100 µm. (**B**) *nilr1-2* mutant induces *CYP71A12* expression similar to Col-0 upon treatment with 50 µM Tre$_{Mi}$31 for 6 hours. Transcript levels of *CYP71A12* in the seedlings were measured by RT-qPCR after normalization to the *U-box* housekeeping gene transcript (*AT5G15400*). Values are presented as mean ± standard error (SE) of three technical replicates, with different letters indicating significant differences ($p \leq 0.0001$, one-way ANOVA with Tukey's post hoc test). All the experiments were repeated three times with consistent results.

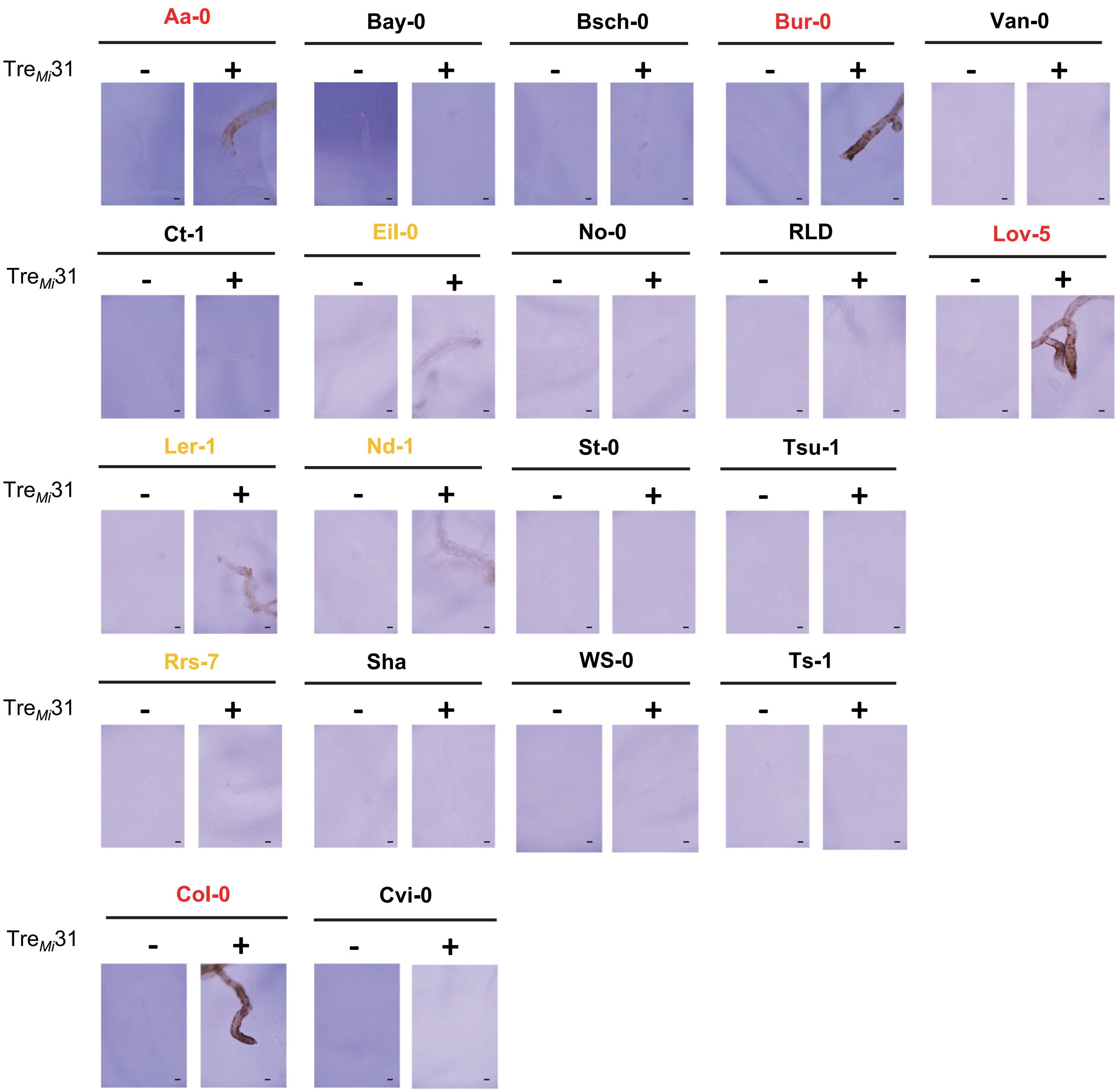


**Fig. S12. Variation in root pigmentation induced by 50 µM Tre$_{Mi}$31 across different *Arabidopsis* accessions.**

Black bars represent 100 µm. Highly sensitive accessions are indicated in red, while moderately sensitive accessions are shown in yellow.

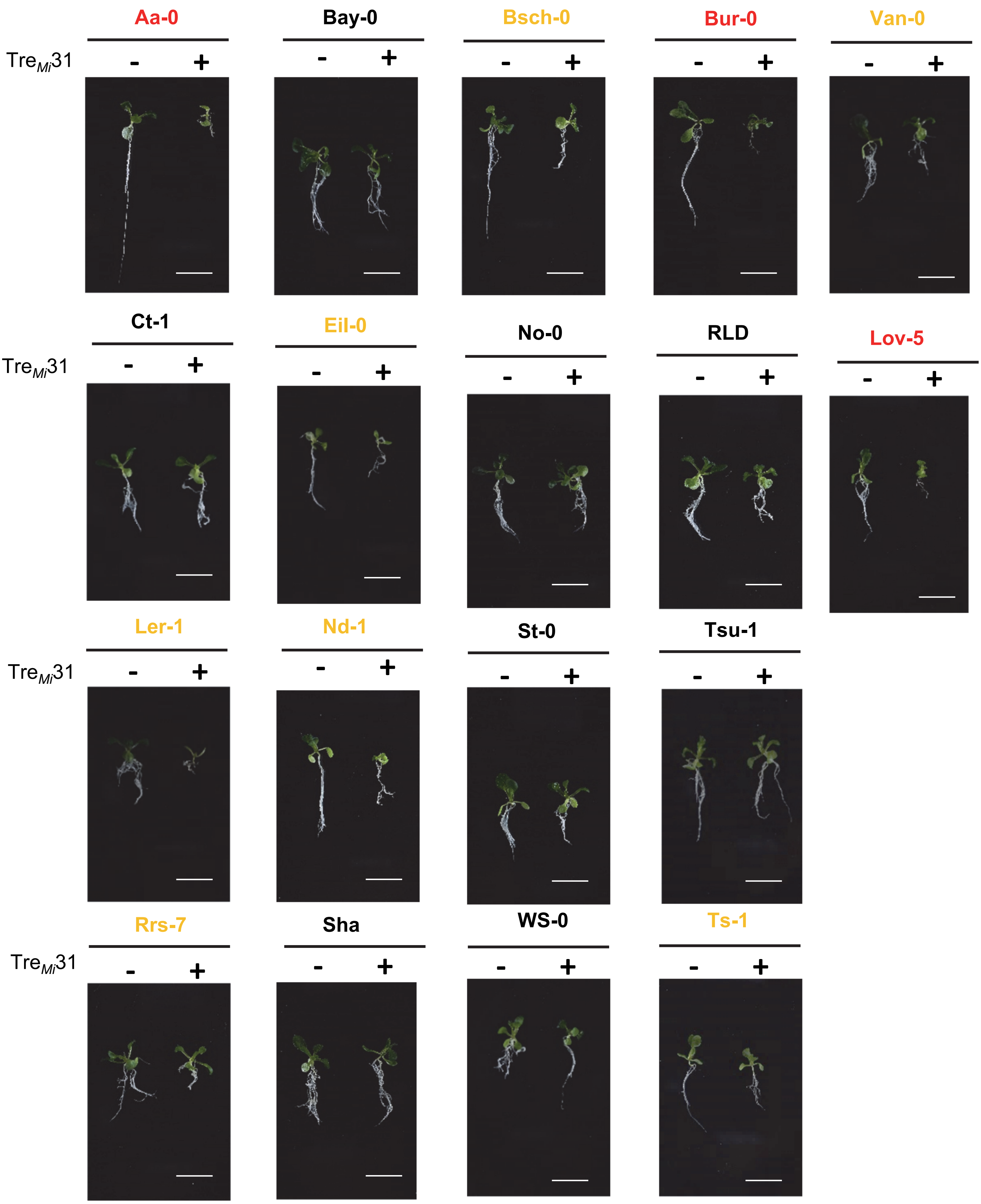


**Fig. S13. Variation in seedling growth inhibition induced by 50 µM Tre$_{Mi}$31 across different *Arabidopsis* accessions.**

White bars represent 1 cm. Highly sensitive accessions are indicated in red, while moderately sensitive accessions are shown in yellow.

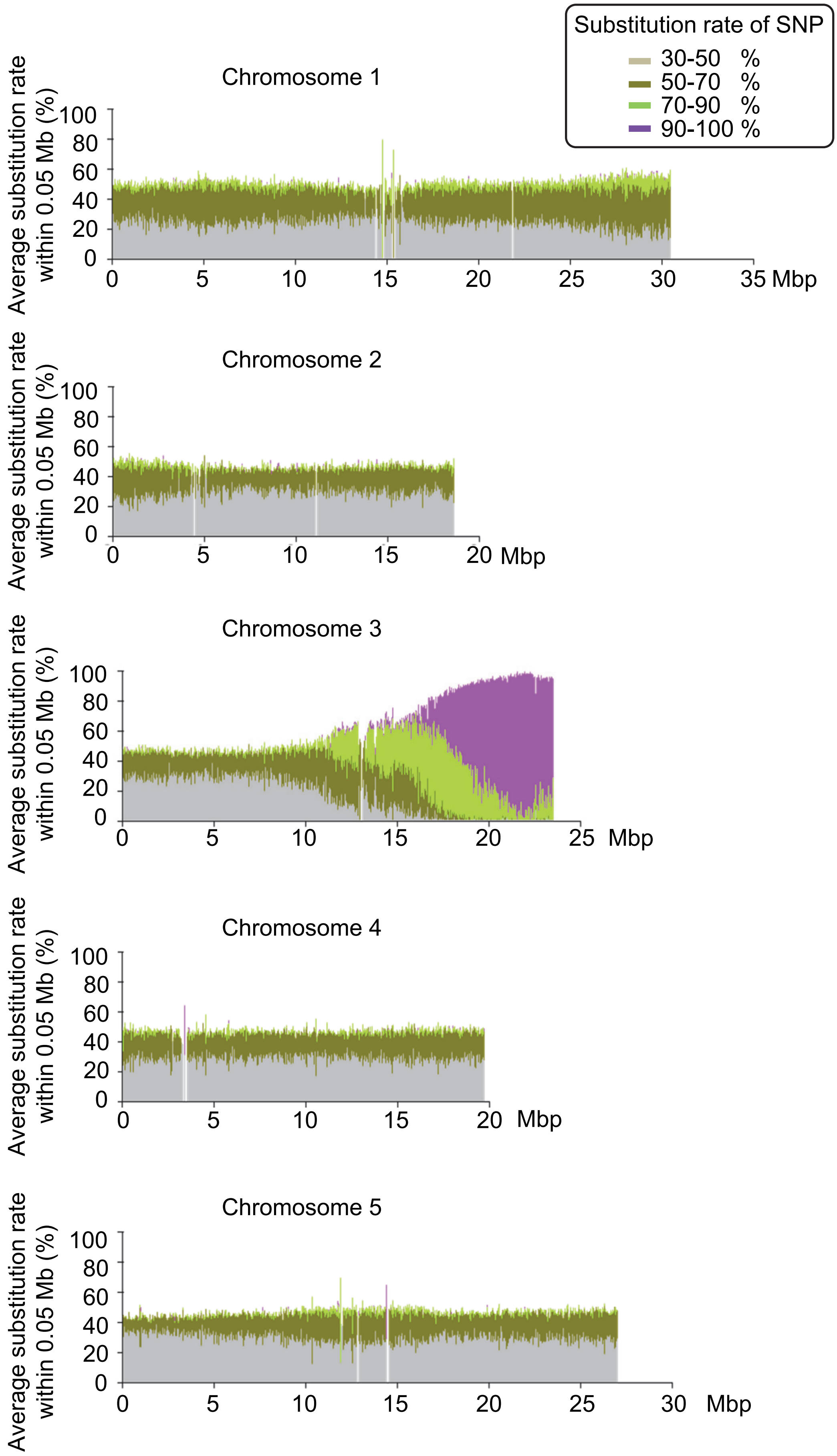


**Fig. S14. Mapping of the chromosomal region required for Tre$_{Mi}$31 sensitivity.**
Genomic sequencing of F2 populations with insensitivity from Col-0 × Cvi-0 crosses identified a genomic region associated with Tre$_{Mi}$31 insensitivity in Cvi-0. Significant accumulations of SNPs from Cvi-0 were detected near the end of chromosome 3. The substitution ratio of SNPs (Col-0 to Cvi-0) was categorized into four levels (30-50%, 50-70%, 70-90%, and 90-100%) and color-coded accordingly. The vertical axis represents the average substitution ratio of SNPs (Cvi-0 / (Col-0 + Cvi-0)) within a 0.05 Mb window.

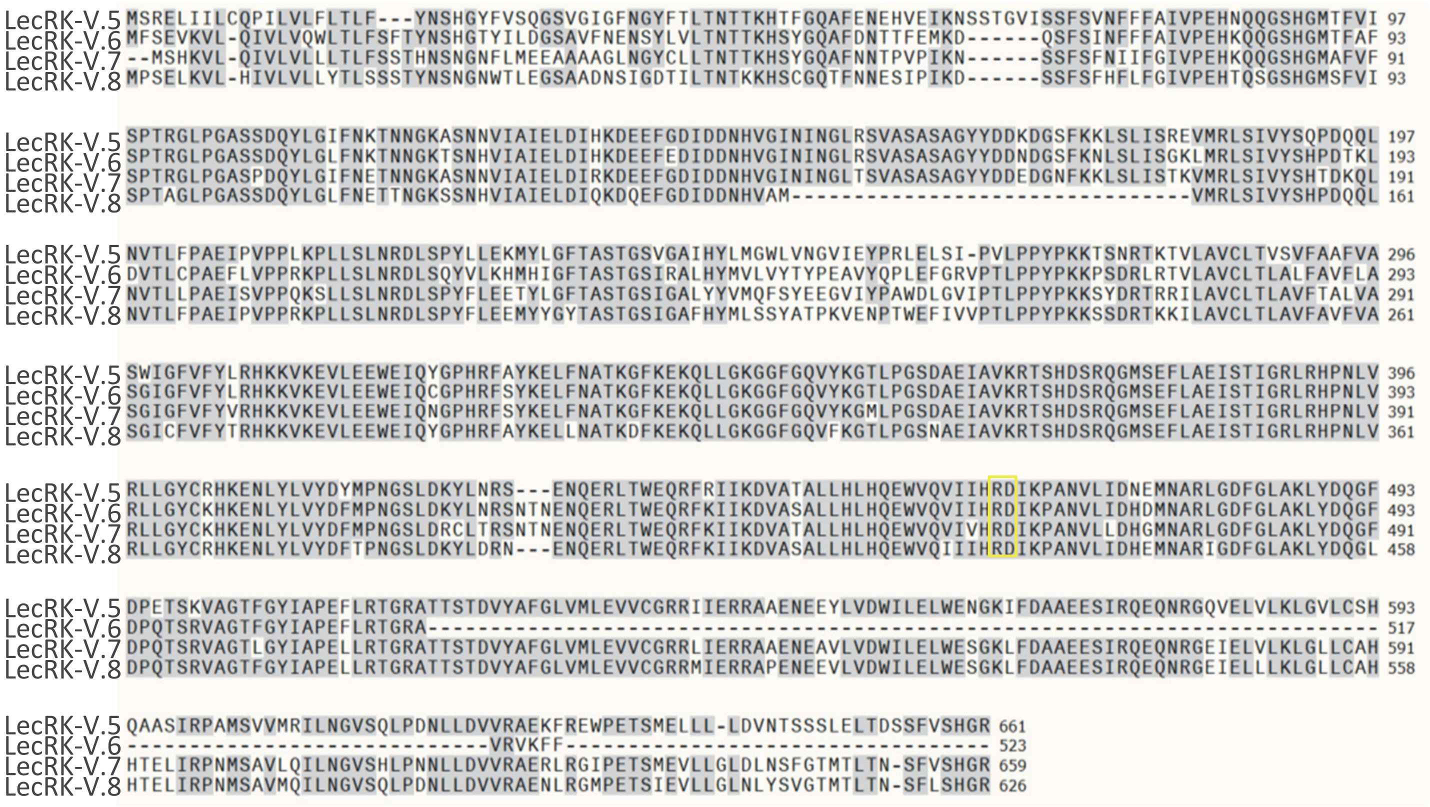


**Fig. S15. Amino acid sequence alignment of LecRK-V.5, V.6, V.7, and V.8.**

The conserved arginine (R) and aspartic acid (D) in the catalytic loop are shown in the yellow box.

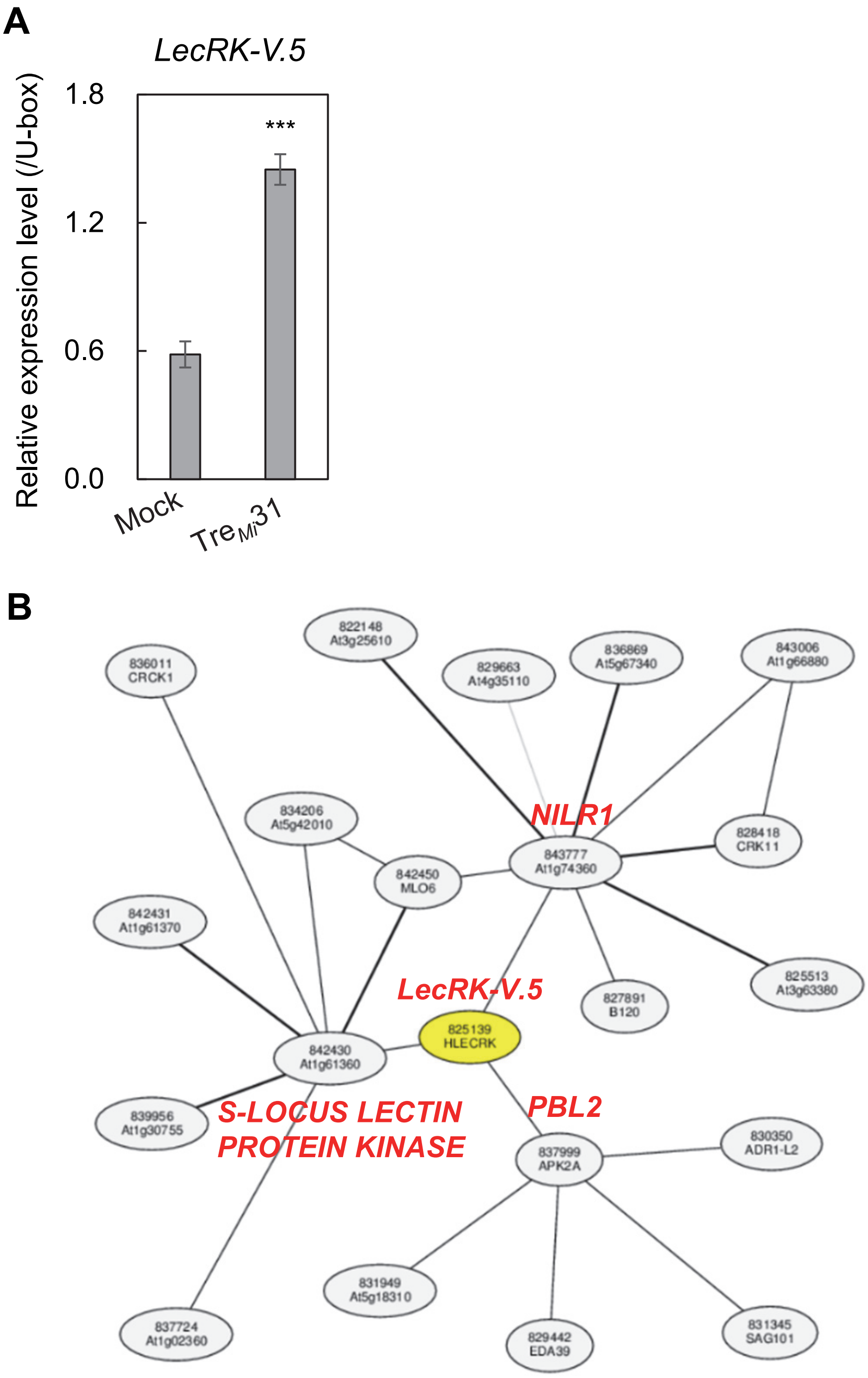


**Genes directly connected with *LecRK-V.5* on the network in ATTED-II**

| Coexpression z score | Locus | Function |
|---|---|---|
| 6.9 | AT1G74360 | NILR1 (NEMATODE-INDUCED LRR-RLK 1), Leucine-rich repeat protein kinase family protein |
| 6.9 | AT1G61360 | S-locus lectin protein kinase family protein |
| 6.1 | AT1G14370 | protein kinase 2A/PBL2 (PBS1-LIKE 2), |

**Fig. S16. *LecRK-V.5* is transcriptionally upregulated by Tre$_{Mi}$31 and co-expressed with *NILR1*.** (**A**) Tre$_{Mi}$31 induces the expression of *LecRK-V.5*. Transcript levels of *LecRK-V.5* in seedlings upon treatment with 40 µM Tre$_{Mi}$31 for 6 hours were measured by RT-qPCR and normalized to the *U-box* housekeeping gene transcript (*AT5G15400*). Values are presented as mean ± SE of three biological replicates, with different letters indicating significant differences ($^{***}$p ≤ 0.001, Student's t-test). (**B**) Co-expression analysis of *LecRK-V.5* using ATTED-II shows that *NILR1* is the most co-expressed gene in *Arabidopsis*.

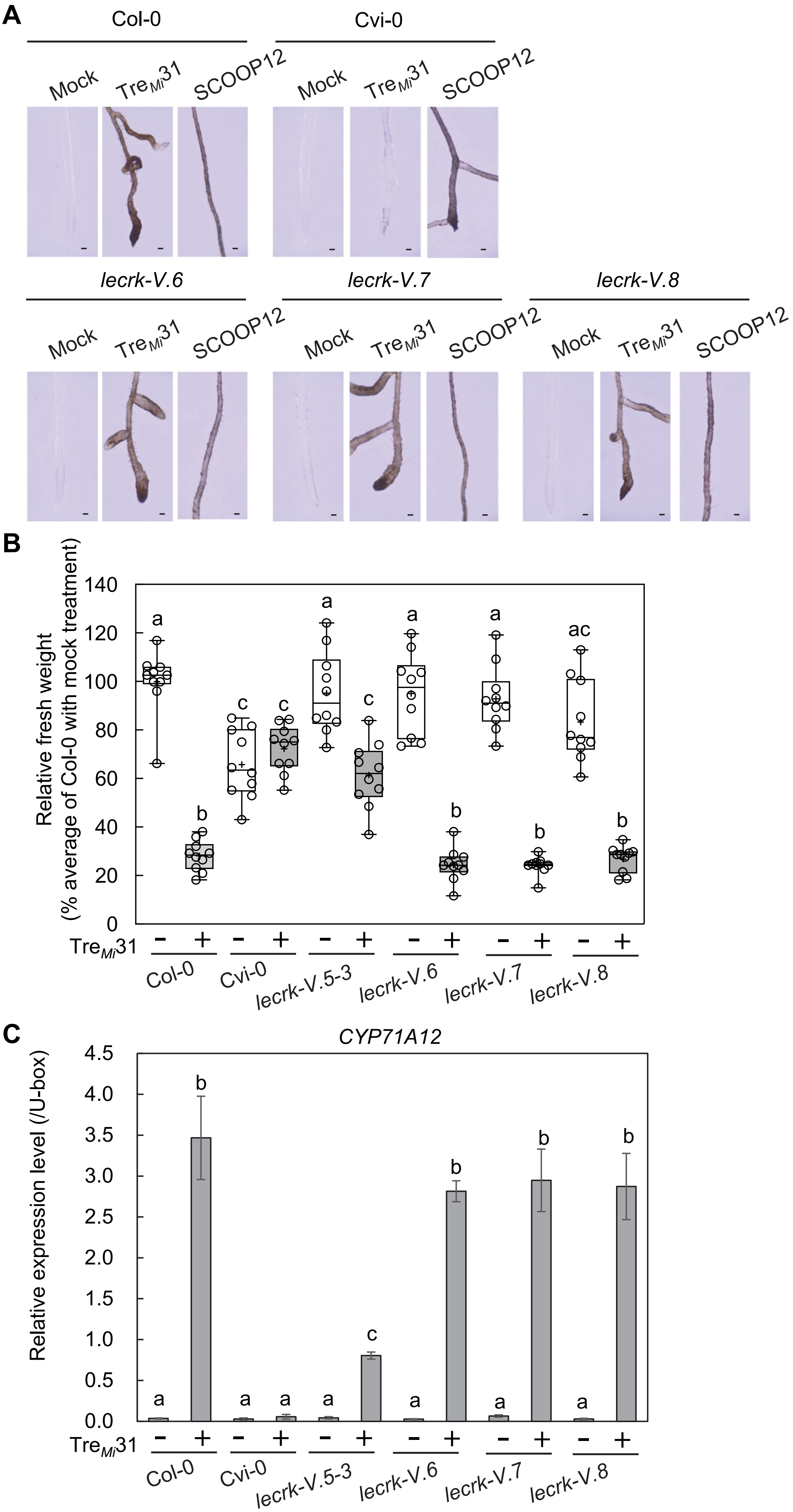

A
Col-0
Cvi-0
Mock
Tre$_{Mi}$31
SCOOP12
lecrk-V.6
lecrk-V.7
lecrk-V.8
B
Relative fresh weight
(% average of Col-0 with mock treatment)
140
120
100
80
60
40
20
0
a
b
c
c
a
c
a
b
a
b
ac
b
Tre$_{Mi}$31
− + − + − + − + − + − +
Col-0
Cvi-0
lecrk-V.5-3
lecrk-V.6
lecrk-V.7
lecrk-V.8
C
CYP71A12
Relative expression level (/U-box)
4.5
4.0
3.5
3.0
2.5
2.0
1.5
1.0
0.5
0.0
a
b
a
a
a
c
a
b
a
b
a
b
Tre$_{Mi}$31
− + − + − + − + − + − +
Col-0
Cvi-0
lecrk-V.5-3
lecrk-V.6
lecrk-V.7
lecrk-V.8

**Fig. S17. T-DNA insertions in *LecRK-V.6*, *LecRK-V.7*, and *LecRK-V.8* do not affect Tre$_{Mi}$31-induced responses.**

(**A** to **C**) *lecrk-V.6*, *lecrk-V.7*, and *lecrk-V.8* mutants exhibit root pigmentation (**A**), growth inhibition (**B**), and *CYP71A12* expression (**C**) in response to 50 μM Tre$_{Mi}$31, similar to Col-0. Black bars in (**A**) represent 100 μm. In the box plot (**B**), the 25th-75th percentiles are shown, with the median indicated by a central line, the mean by a black cross, and whiskers representing the full range. Each open circle represents one data point from 10 samples (with one seedling per sample). Different letters denote significant differences ($p \leq 0.05$, one-way ANOVA with Tukey's post hoc test). Experiments were repeated three times with consistent results. In the bar chart (**C**), transcript levels of *CYP71A12* in the seedlings upon treatment with 50 μM Tre$_{Mi}$31 for 6 hours were measured by RT-qPCR after normalization to the *U-box* housekeeping gene transcript (*AT5G15400*). Values are presented as mean ± standard error (SE) of three biological replicates, with different letters indicating significant differences ($p \leq 0.05$, one-way ANOVA with Tukey's post hoc test).

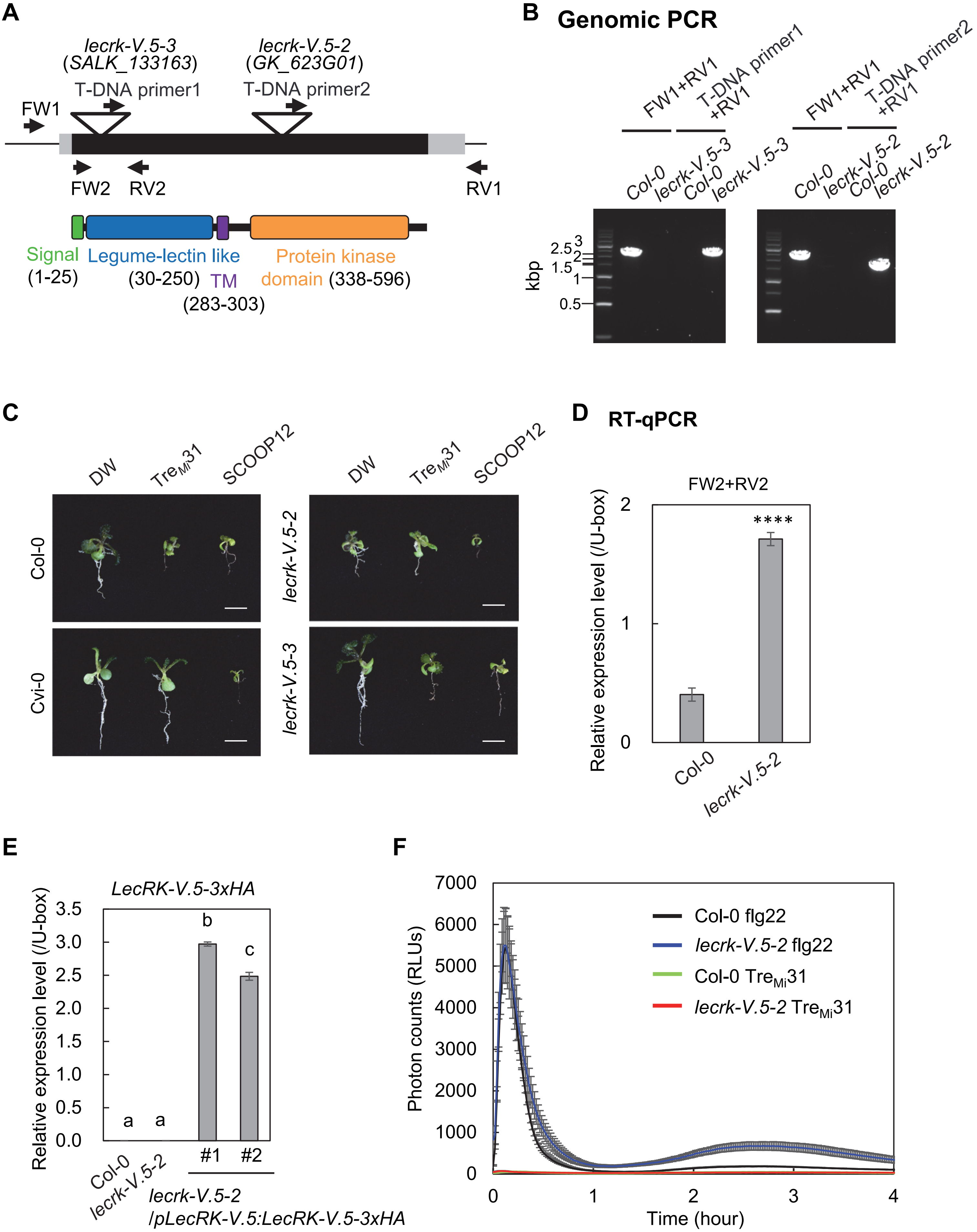
A
lecrk-V.5-3
(SALK_133163)
T-DNA primer1
lecrk-V.5-2
(GK_623G01)
T-DNA primer2
FW1
FW2
RV2
RV1
Signal
(1-25)
Legume-lectin like
(30-250)
TM
(283-303)
Protein kinase
domain (338-596)
B
Genomic PCR
FW1+RV1
T-DNA primer1
+RV1
FW1+RV1
T-DNA primer2
+RV1
Col-0
lecrk-V.5-3
Col-0
lecrk-V.5-3
Col-0
lecrk-V.5-2
Col-0
lecrk-V.5-2
kbp
3
2.5
2
1.5
1
0.5
C
DW
Tre$_{Mi}$31
SCOOP12
DW
Tre$_{Mi}$31
SCOOP12
Col-0
Cvi-0
lecrk-V.5-2
lecrk-V.5-3
D
RT-qPCR
FW2+RV2
Relative expression level (/U-box)
2
1
0
****
Col-0
lecrk-V.5-2
E
LecRK-V.5-3xHA
Relative expression level (/U-box)
3.5
3.0
2.5
2.0
1.5
1.0
0.5
0.0
b
c
a
a
Col-0
lecrk-V.5-2
#1
#2
lecrk-V.5-2
/pLecRK-V.5:LecRK-V.5-3xHA
F
Photon counts (RLUs)
7000
6000
5000
4000
3000
2000
1000
0
Col-0 flg22
lecrk-V.5-2 flg22
Col-0 Tre$_{Mi}$31
lecrk-V.5-2 Tre$_{Mi}$31
0
1
2
3
4
Time (hour)

**Fig. S18. *LecRK-V.5* is required for Tre$_{Mi}$31 signaling.**
(**A**) Schematic representation of the *LecRK-V.5* gene, showing the positions of T-DNA insertions and primers used for genotyping and RT-qPCR. The *LecRK-V.5* gene has no introns, with exons and untranslated regions depicted in black and gray, respectively. (**B**) Genotyping of *lecrk-V.5-2* and *lecrk-V.5-3* mutants. (**C**) *lecrk-V.5-3* mutants exhibit reduced seedling growth inhibition upon treatment with 50 µM Tre$_{Mi}$31 compared to Col-0, while *lecrk-V.5-2* mutants and Cvi-0 show no responses. Two µM SCOOP12 induces seedling growth inhibition in all genotypes. White bars represent 1 cm. (**D**) Transcript levels of truncated *LecRK-V.5* in *lecrk-V.5-2* mutants were measured by RT-qPCR after normalization to the *U-box* housekeeping gene transcript (*AT5G15400*). Values are presented as mean ± SE of three technical replicates, with asterisks indicating significant differences ($^{****}p \leq 0.0001$, Student's t-test). (**E**) Transcript levels of *LecRK-V.5-3xHA* in the complementation lines of *lecrk-V.5-2 /pLecRK-V.5:LecRK-V.5-3xHA* quantified by RT-qPCR, normalized to the *U-box* housekeeping gene transcript (*AT5G15400*). Values are presented as mean ± SE of three technical replicates, with different letters denoting significant differences ($p \leq 0.0001$, one-way ANOVA with Tukey's post hoc test). (**F**) Treatment with 50 µM Tre$_{Mi}$31 does not induce ROS production in Col-0. Flg22 was used as a positive control. Time course of ROS production was measured by a luminol-based assay, with results shown in relative luminescence units (RLUs). Experiments were repeated three times with consistent results.

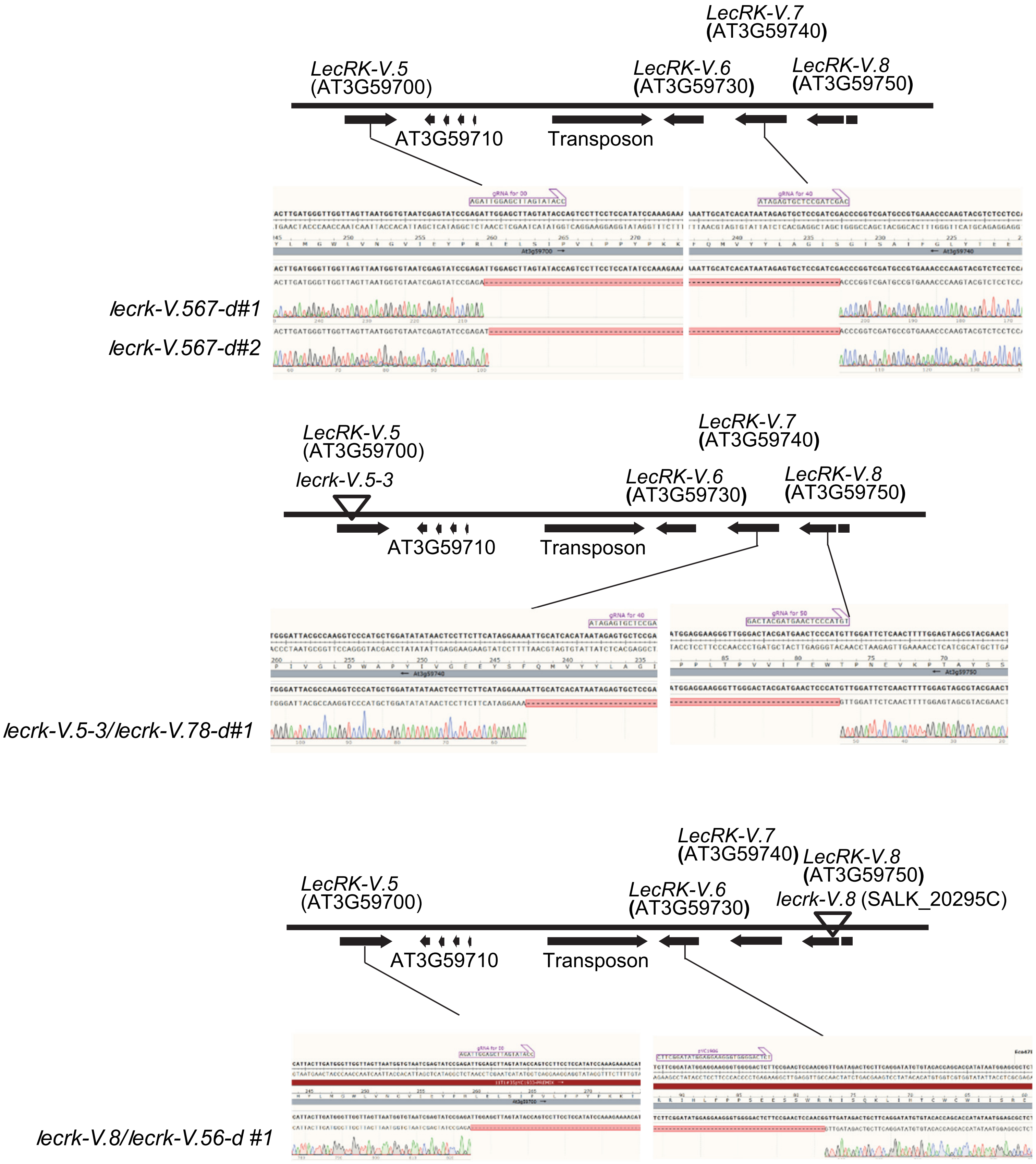


**Fig. S19. Confirmation of deletion of genes in CRISPR lines, *lecrk-V.567-d*, *lecrk-V.5-3/lecrk-V.78-d*, and *lecrk-V.8/lecrk-V.56-d* lines by Sanger sequencing.**

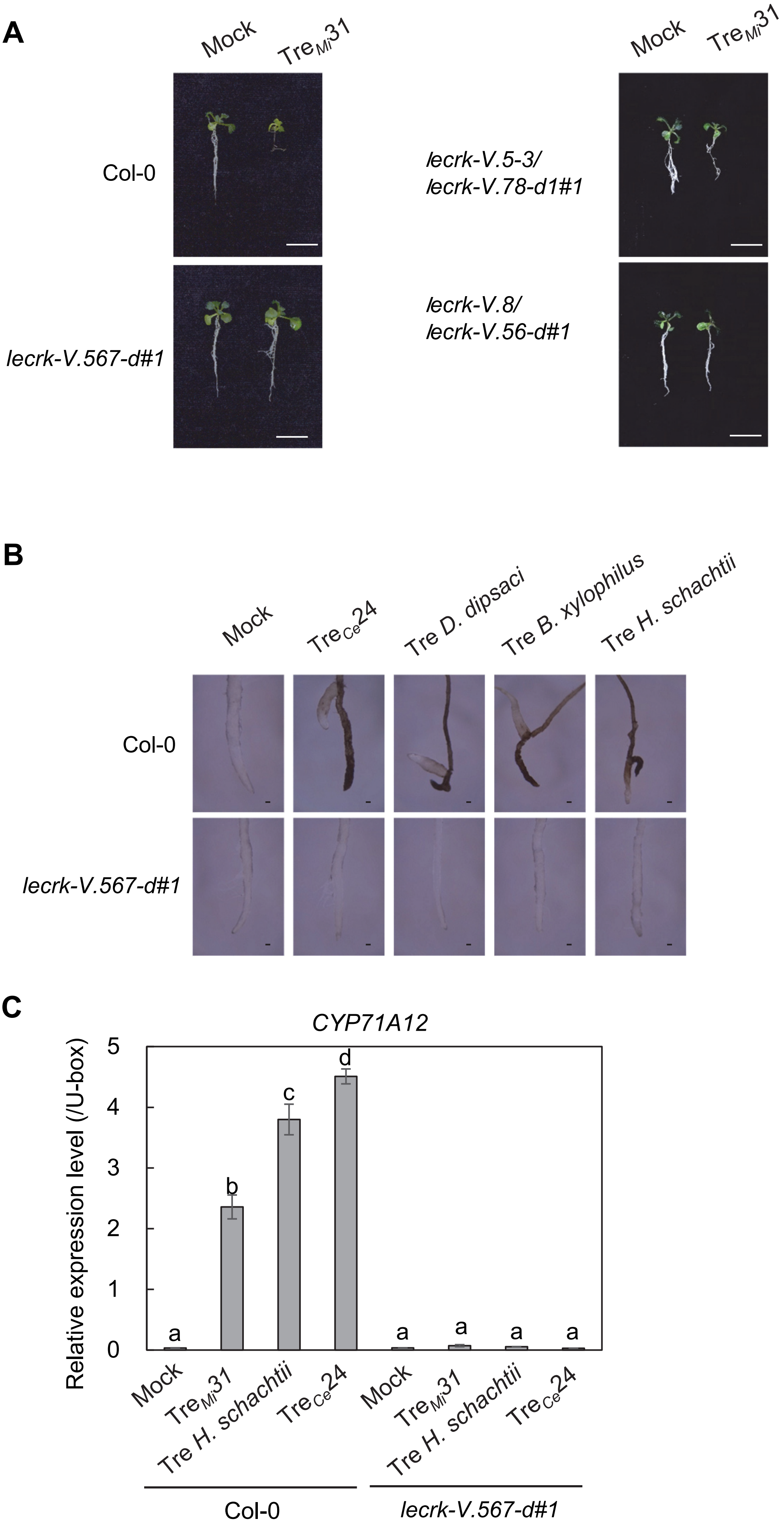

A
Mock
Tre$_{Mi}$31
Col-0
lecrk-V.567-d#1
lecrk-V.5-3/
lecrk-V.78-d1#1
lecrk-V.8/
lecrk-V.56-d#1
B
Mock
Tre$_{Ce}$24
Tre D. dipsaci
Tre B. xylophilus
Tre H. schachtii
Col-0
lecrk-V.567-d#1
C
CYP71A12
Relative expression level (/U-box)
0
1
2
3
4
5
a
b
c
d
a
a
a
a
Mock
Tre$_{Mi}$31
Tre H. schachtii
Tre$_{Ce}$24
Col-0
lecrk-V.567-d#1

**Fig. S20. *LecRK-V.5* and *LecRK-V.6* are involved in Tre$_{Mi}$31 recognition.**

(**A**) CRISPR deletion line *lecrk-V.5-3/lecrk-V.78-d* shows reduced seedling growth inhibition upon treatment with 50 µM Tre$_{Mi}$31, while *lecrk-V.567-d* and *lecrk-V.8/lecrk-V.56-d* lines do not induce the responses. White bars represent 1 cm. (**B**) *lecrk-V.567-d*#1 line does not induce root pigmentation upon treatment with 50 µM trehalose-derived peptides from PPNs. Black bars represent 100 µm. (**C**) *lecrk-V.567-d*#1 line does not induce *CYP71A12* expression in response to 50 µM Tre$_{Ce}$24 and the trehalose-derived peptide from *H. schachtii* (Tre *H. schachtii*). Transcript levels of *CYP71A12* in the seedlings upon treatment with 50 µM Tre$_{Ce}$24, Tre$_{Mi}$31, or Tre *H. schachtii* were measured by RT-qPCR after normalization to the *U-box* housekeeping gene transcript (*AT5G15400*). Values are presented as mean ± SE of three technical replicates, with different letters indicating significant differences ($p \leq 0.05$, one-way ANOVA with Tukey's post hoc test). Experiments were repeated three times with consistent results.

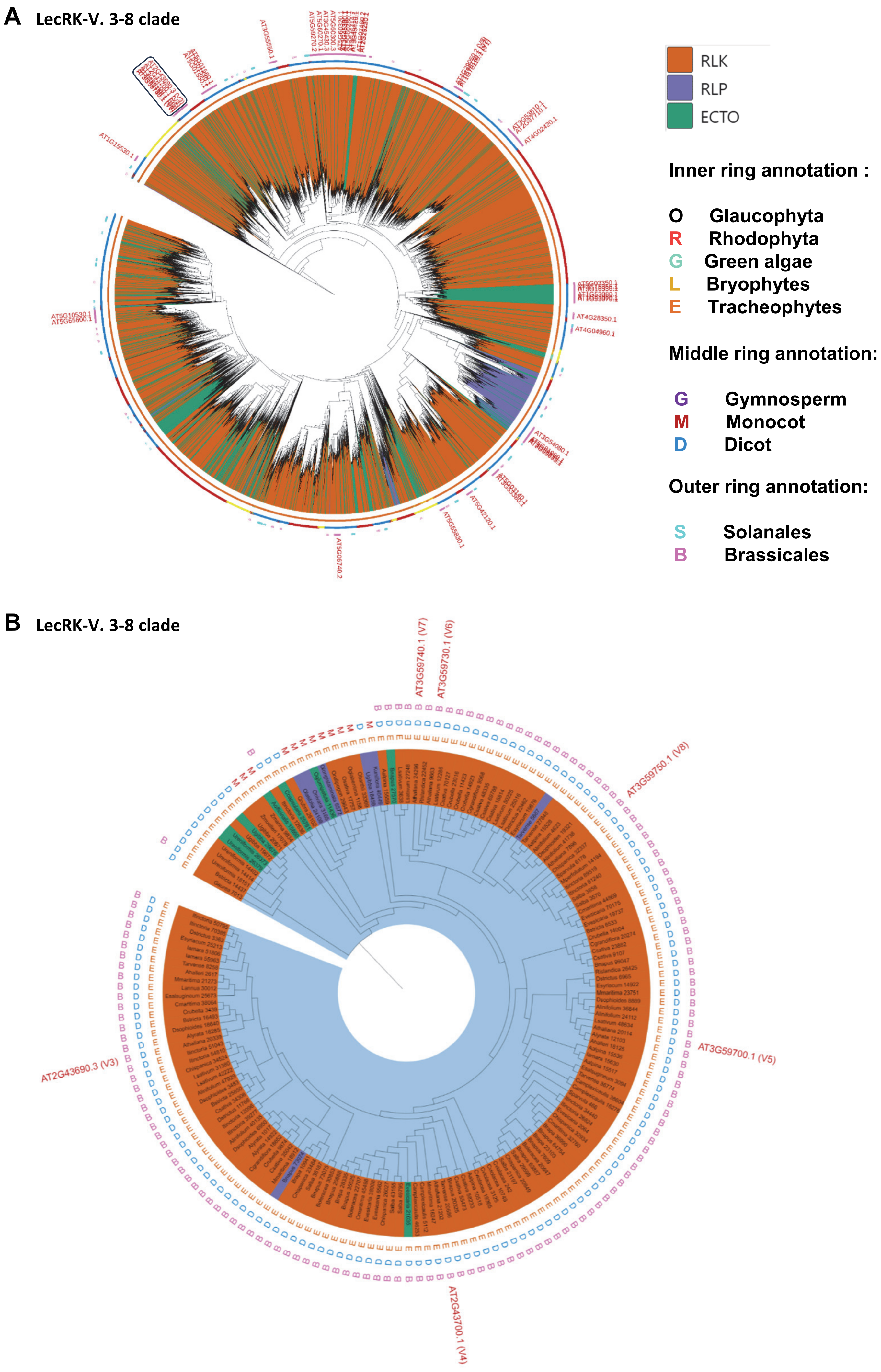

A
LecRK-V. 3-8 clade
RLK
RLP
ECTO
Inner ring annotation :
O Glaucophyta
R Rhodophyta
G Green algae
L Bryophytes
E Tracheophytes
Middle ring annotation:
G Gymnosperm
M Monocot
D Dicot
Outer ring annotation:
S Solanales
B Brassicales
AT1G15530.1
AT3G55550.1
AT4G02420.1
AT4G28350.1
AT4G04960.1
AT5G10530.1
AT5G65600.1
AT3G54080.1
AT5G42120.1
AT5G55830.1
AT5G06740.2
B
LecRK-V. 3-8 clade
AT3G59740.1 (V7)
AT3G59730.1 (V6)
AT3G59750.1 (V8)
AT3G59700.1 (V5)
AT2G43690.3 (V3)
AT2G43700.1 (V4)

**Fig. S21. Phylogenetic analysis reveals that *LecRK-V.5, LecRK-V.6, LecRK-V.7,* and *LecRK-V.8* cluster together and are conserved only in Brassicales.**

(**A**) Ectodomain phylogenetic tree of all L-lectins (14,465 members) across 350 species from Glaucophyta, red algae, green algae, Bryophytes, and Tracheophytes (*14, 15*), shows that *LecRK-V.3* to *LecRK-V.8* clusters within the same clade. (**B**) Ectodomain phylogenetic tree of *LecRK-V.3* to *LecRK-V.8*. Branches: Orange-RLK, Purple-RLP, Green-ectodomain-only proteins (lacking kinase and transmembrane regions). Inner ring: Black-Glaucophyta, Red-Rhodophyta, Light green-green algae, Light orange-bryophytes, and Orange-tracheophytes. Middle ring: Yellow-basal angiosperms, Red-monocots, Blue-dicots, and Purple-gymnosperms. Outer ring: Pink-Brassicales, Cyan-Solanales. *Arabidopsis* members are labeled.

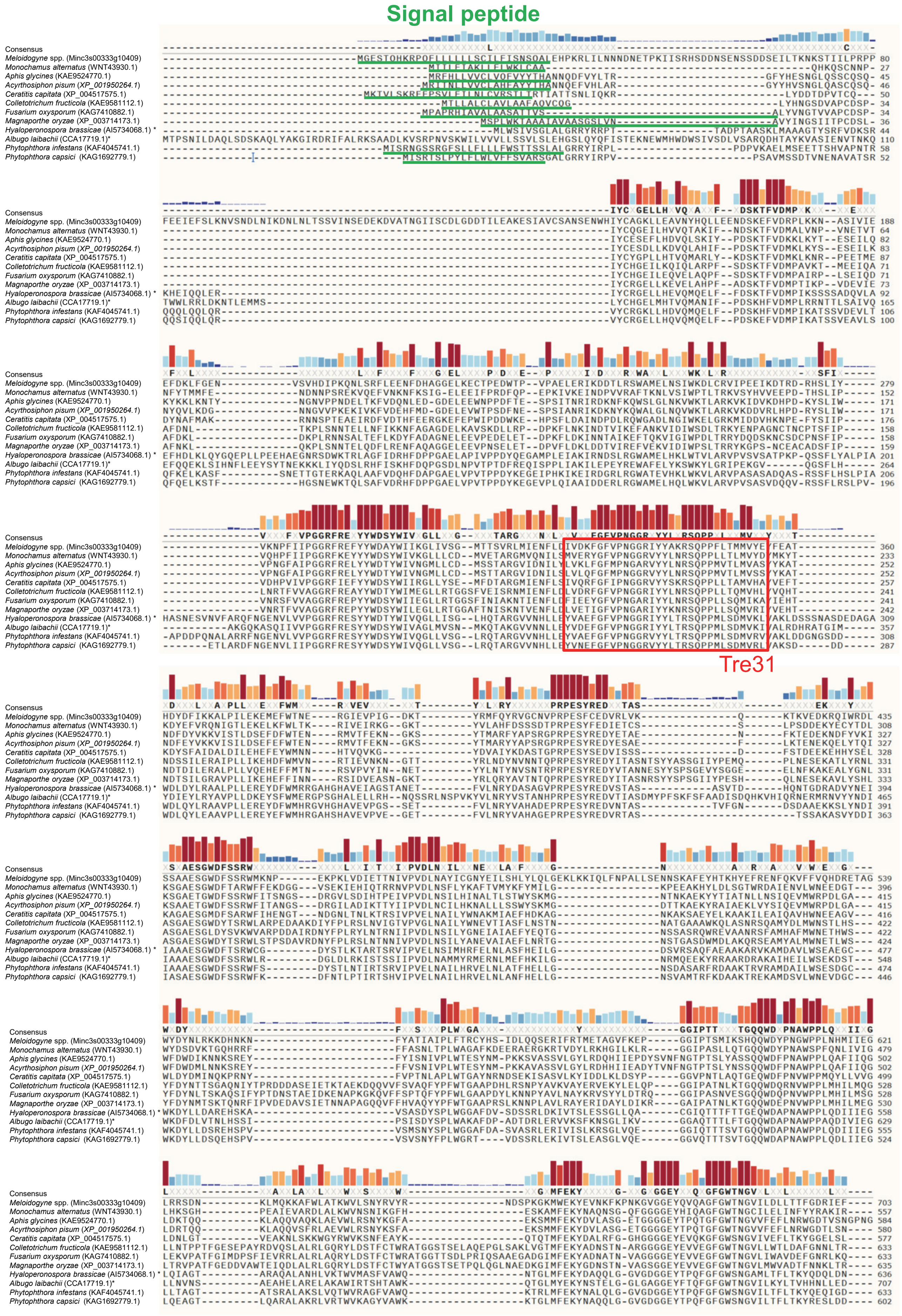
Signal peptide
Tre31
Consensus
Meloidogyne spp. (Minc3s00333g10409)
Monochamus alternatus (WNT43930.1)
Aphis glycines (KAE9524770.1)
Acyrthosiphon pisum (XP_001950264.1)
Ceratitis capitata (XP_004517575.1)
Colletotrichum fructicola (KAE9581112.1)
Fusarium oxysporum (KAG7410882.1)
Magnaporthe oryzae (XP_003714173.1)
Hyaloperonospora brassicae (AI5734068.1) *
Albugo laibachii (CCA17719.1)*
Phytophthora infestans (KAF4045741.1)
Phytophthora capsici (KAG1692779.1)

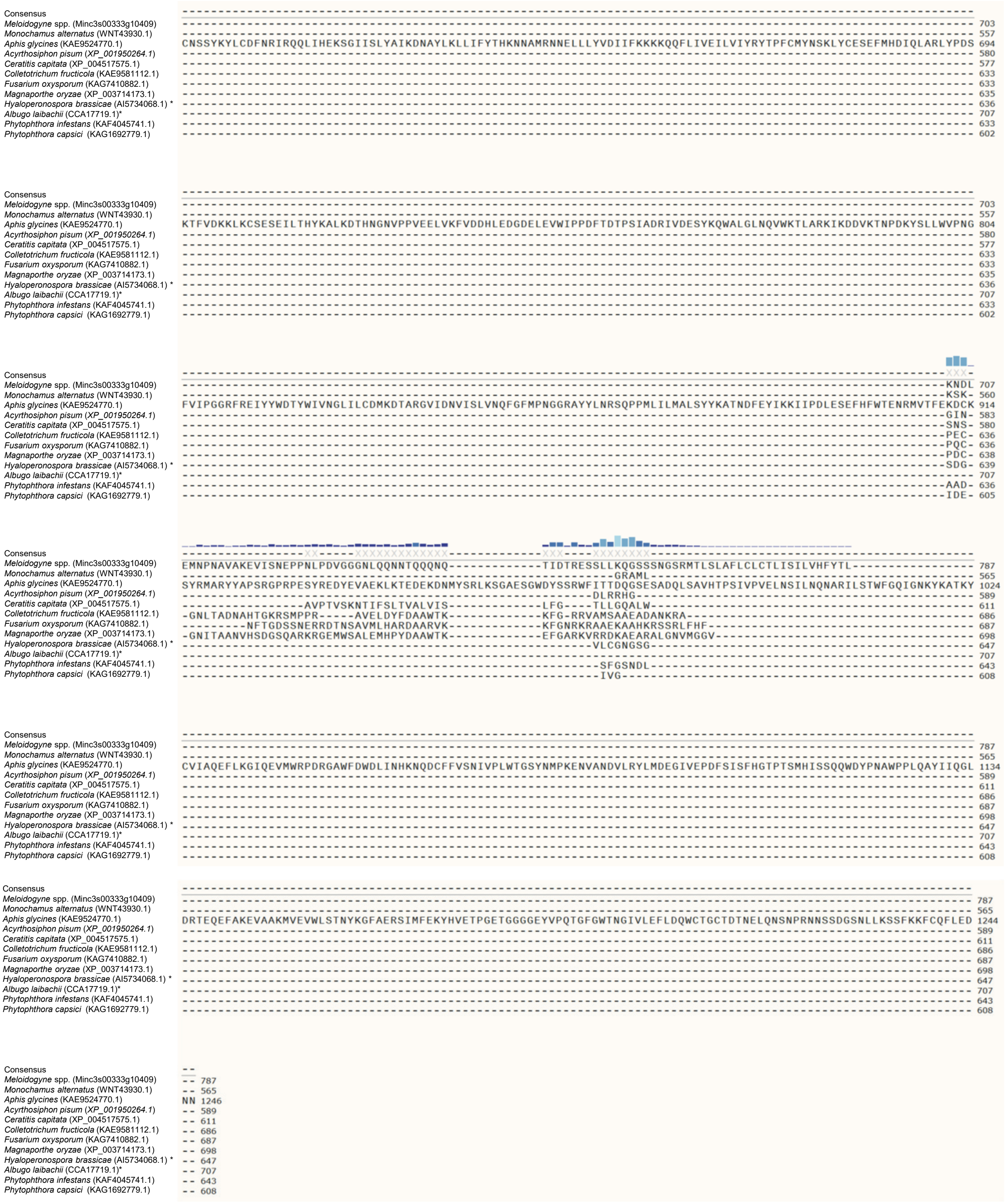


**Fig. S22. Amino acid sequence alignment of secreted trehalase proteins of pathogenic insects and fungi.** Regions corresponding to predicted signal peptides by SignalP-6.0 are highlighted with green lines. The 31 amino acid residues of PPNs (Tre31) are shown in a red box. The degree of conservation of each residue among sequences is displayed as colored bars above the alignment.

**A**

*cerk1 fls2 pCYP71A12:GUS*

| Name | Sequence | 50 µM |
|---|---|---|
| Tre *Aphis glycines* | VKLFGFMPNGARVYYLNRSQPPMVTLMV | 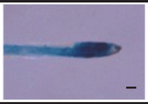 |
| Tre *Acyrthosiphon pisum* | VLQFGFMPNGGRVYYLNRSQPPMVTLMV | 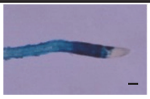 |
| Tre *Ceratitis capitata* | VQRFGFIPNGGRVYYSKRSQPPLLTAMV | 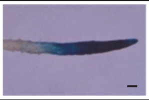 |
| Tre *Colletotrichum fructicola* | VDRFGFVPNGGRIYYLNRSQPPLLTQMV | 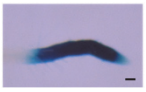 |

**B**

*cerk1 fls2 pCYP71A12:GUS*

| Name | Sequence | 50 µM |
|---|---|---|
| $Tre_{Ce}19$ | VDKYGFVPNGGRVYYLQRS | 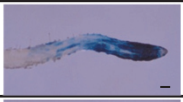 |
| $Tre_{Ce}19$_K3E | VDEYGFVPNGGRVYYLQRS | 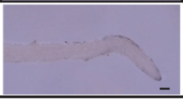 |

**C**

*cerk1 fls2 pCYP71A12:GUS*

| Name | Sequence | 50 µM |
|---|---|---|
| $Tre_{Mi}19$ | VDKFGFVPNGGRIYYAKRS | 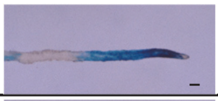 |
| $Tre_{Mi}19$_K3E | VDEFGFVPNGGRIYYAKRS | 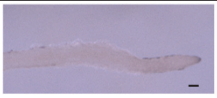 |

**D**

Col-0 *lecrk-V.567-d#1*

Tre *A. glycines*

Tre *A. pisum*

Tre *C. capitata*

Tre *C. fructicola*

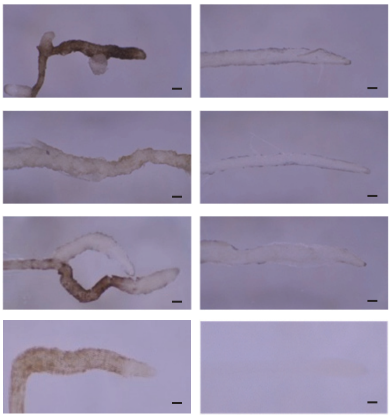

**Fig. S23. Characterization of trehalase peptides from plant pathogenic fungi and insects.**
(**A**) Trehalase peptides from plant pathogenic insects and fungi induce *CYP71A12* expression in *cerk1 fls2 pCYP71A12:GUS* line. (**B** and **C**) Charge-reverse substitution at the third lysine residue to glutamic acid (K3E) in $Tre_{Ce}19$ and $Tre_{Mi}19$ results in loss of MAMP activity. (**D**) *lecrk-V.567-d*#1 line does not induce root pigmentation in response to 50 µM trehalase-derived peptides from plant pathogenic insects and fungi. Black bars represent 100 µm. Experiments were repeated three times with consistent results.

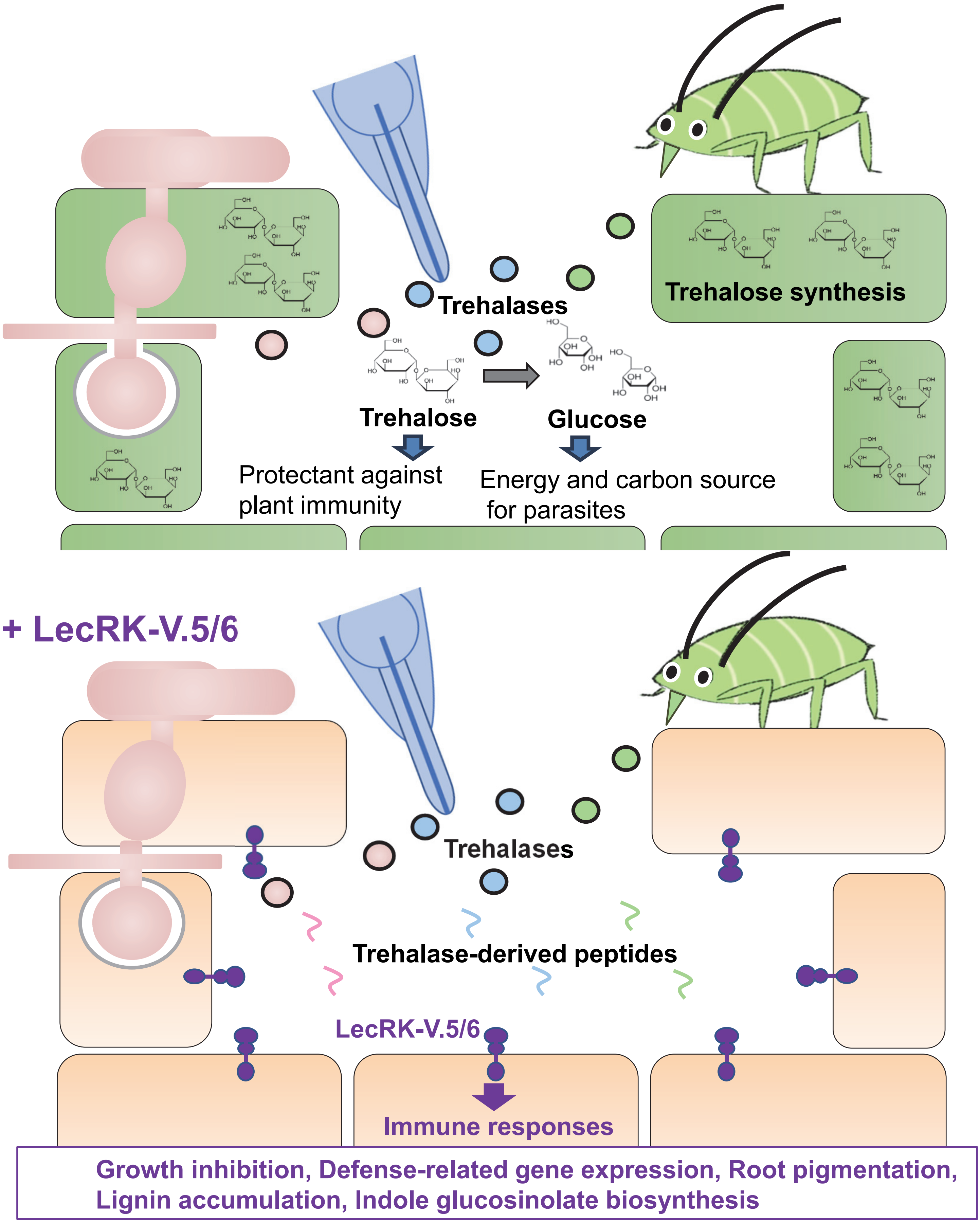


**Fig. S24. Model of *Arabidopsis* Col-0 and PPN interaction through apoplastic trehalases and *LecRK-V.5* and *LecRK-V.6*.**

Trehalase proteins are produced during the migratory stage of some PPNs and are likely secreted into the apoplast, where they catalyze the conversion of trehalose into glucose. Similarly, trehalase proteins may also be secreted by certain fungal pathogens and insect parasites (*17*). Trehalose serves as an energy and carbon source for parasites and acts as a protectant against host immune defenses, while trehalase may facilitate glucose absorption by these organisms. In the Col-0 accession of *Arabidopsis*, these secreted trehalases or their peptide fragments are recognized, likely through LecRK-V.5 and LecRK-V.6. This recognition of trehalase-derived peptides activates immune responses, including growth inhibition, defense-related gene expression, root pigmentation, lignin accumulation, and indole glucosinolate biosynthesis.

**Other Supplementary Materials for this manuscript include the following:**

**Table S1. Genes upregulated or downregulated after treatment with *C. elegans* extract compared to mock treatment in Col-0.**

**Table S2. Genes upregulated or downregulated after treatment with *M. incognita* extract compared to mock treatment in Col-0.**

**Table S3. Gene Ontology (GO) enrichment analysis of genes upregulated or downregulated after treatment with *M. incognita* or *C. elegans* extract.**

**Table S4. List of genes and isoforms with differential expression after treatment with *M. incognita* extract, *C. elegans* extract, $Tre_{Mi}31$, flg22, and chitin.**

**Table S5. List of MAMP candidate proteins identified by chromatography purification and LC-MS/MS analyses.**

**Table S6. List of identified peptides of *Ce*TRE3 by LC-MS/MS analyses.**

**Table S7. Genes upregulated or downregulated after treatment with $Tre_{Mi}31$.**

**Table S8. Gene Ontology (GO) enrichment analysis of genes upregulated or downregulated after treatment with $Tre_{Mi}31$.**

**Table S9. Root pigmentation phenotypes of Recombinant inbred lines (RILs).**

**Table S10. SNPs analyses of RILs by Sanger sequencing.**

**Table S11. Phenotypes of T-DNA insertion mutants of the genes in the narrowed-down region.**

**Table S12. Genes upregulated or downregulated after treatment with $Tre_{Mi}31$ in Col-0 and *lecrk V.5-3* mutant.**

**Table S13. Primers that were used in this paper.**

### References


1. B. T. Townsley, M. F. Covington, Y. Ichihashi, K. Zumstein, N. R. Sinha, BrAD-seq: Breath Adapter Directional sequencing: a streamlined, ultra-simple and fast library preparation protocol for strand specific mRNA library construction. *Frontiers in Plant Science* **6**, 366 (2015).
2. B. Langmead, C. Trapnell, M. Pop, S. L. Salzberg, Ultrafast and memory-efficient alignment of short DNA sequences to the human genome. *Genome Biol* **10**, R25 (2009).
3. T. Suzuki *et al.*, The DROL1 subunit of U5 snRNP in the spliceosome is specifically required to splice AT-AC-type introns in Arabidopsis. *Plant J* **109**, 633-648 (2022).
4. T. Hulsen, J. de Vlieg, W. Alkema, BioVenn - a web application for the comparison and visualization of biological lists using area-proportional Venn diagrams. *BMC Genomics* **9**, 488 (2008).
5. M. Safaeizadeh, T. Boller, C. Becker, Comparative RNA-seq analysis of Arabidopsis thaliana response to AtPep1 and flg22, reveals the identification of PP2-B13 and ACLP1 as new members in pattern-triggered immunity. *PLoS One* **19**, e0297124 (2024).
6. J. Wan *et al.*, A LysM receptor-like kinase plays a critical role in chitin signaling and fungal resistance in Arabidopsis. *Plant Cell* **20**, 471-481 (2008).
7. S. Babicki *et al.*, Heatmapper: web-enabled heat mapping for all. *Nucleic Acids Res* **44**, W147-153 (2016).
8. K. Sato *et al.*, Transcriptomic Analysis of Resistant and Susceptible Responses in a New Model Root-Knot Nematode Infection System Using Solanum torvum and Meloidogyne arenaria. *Front Plant Sci* **12**, 680151 (2021).
9. K. Kano, S. Noda, S. Sato, K. Kuwata, E. Mishiro-Sato, An efficient in-gel digestion method on small amounts of protein sample from large intact gel pieces. *SEPARATION SCIENCE PLUS* **6**, 2200121 (2023).
10. Y. Kadota *et al.*, Quantitative phosphoproteomic analysis reveals common regulatory mechanisms between effector- and PAMP-triggered immunity in plants. *New Phytol* **221**, 2160–2175 (2019).
11. H. Tsutsui, T. Higashiyama, pKAMA-ITACHI Vectors for Highly Efficient CRISPR/Cas9-Mediated Gene Knockout in Arabidopsis thaliana. *Plant Cell Physiol* **58**, 46-56 (2017).
12. H. Nishiyama *et al.*, Protocol for root-knot nematode culture by a hydroponic system and nematode inoculation to Arabidopsis. *Nematological Research (Japanese Journal of Nematology)* **45**, 45-49 (2015).
13. Y. Goto *et al.*, Exogenous Treatment with Glutamate Induces Immune Responses in Arabidopsis. *Mol Plant Microbe Interact* **33**, 474-487 (2020).
14. B. P. M. Ngou, M. Wyler, M. W. Schmid, Y. Kadota, K. Shirasu, Evolutionary trajectory of pattern recognition receptors in plants. *Nat Commun* **15**, 308 (2024).
15. B. P. M. Ngou, R. Heal, M. Wyler, M. W. Schmid, J. D. G. Jones, Concerted expansion and contraction of immune receptor gene repertoires in plant genomes. *Nat Plants* **8**, 1146-1152 (2022).
16. T. Obayashi, H. Hibara, Y. Kagaya, Y. Aoki, K. Kinoshita, ATTED-II v11: A Plant Gene Coexpression Database Using a Sample Balancing Technique by Subagging of Principal Components. *Plant Cell Physiol* **63**, 869-881 (2022).
17. Y. Zhang, J. Fan, J. Sun, F. Francis, J. Chen, Transcriptome analysis of the salivary glands of the grain aphid, Sitobion avenae. *Sci Rep* **7**, 15911 (2017).
